# GTO: a toolkit to unify pipelines in genomic and proteomic research

**DOI:** 10.1101/2020.01.07.882845

**Authors:** João R. Almeida, Armando J. Pinho, José L. Oliveira, Olga Fajarda, Diogo Pratas

## Abstract

**Summary:** Next-generation sequencing triggered the production of a massive volume of publicly available data and the development of new specialised tools. These tools are dispersed over different frameworks, making the management and analyses of the data a challenging task. Additionally, new targeted tools are needed, given the dynamics and specificities of the field. We present GTO, a comprehensive toolkit designed to unify pipelines in genomic and proteomic research, which combines specialised tools for analysis, simulation, compression, development, visualisation, and transformation of the data. This toolkit combines novel tools with a modular architecture, being an excellent platform for experimental scientists, as well as a useful resource for teaching bioinformatics inquiry to students in life sciences.

**Availability and implementation:** GTO is implemented in C language and it is available, under the MIT license, at http://bioinformatics.ua.pt/gto.

**Supplementary information:** Supplementary data are available at publisher’s Web site.

## 1. Introduction

Next-generation sequencing (NGS) has become an essential tool in genetic and genomic analysis with substantial impact in the fields of biomedicine and anthropology. The advantages of NGS over traditional methods include its multiplex capability and analytical resolution, making it a time and cost-efficient approach for fast clinical and forensic screening [1]. The development of efficient bioinformatics tools is essential to assess and analyse the large volumes of sequencing data produced by next-generation sequencers. However, more important than that, are the computational methods that unify the existing tools, given the notable pace at which these tools are made available.

Toolkits are sets of tools that combine multiple features in a custom-based manner as some examples show, both in genomics [2] and proteomics [3]. Developing a toolkit requires a specific architecture, namely, taking into account the purpose and technologies, accessibility, compatibility, portability, interoperability, and usability. Moreover, the implementation needs to consider efficiency, while maintaining affordable computational resources and the absence of dependencies (standalone use).

We contribute with GTO, a standalone toolkit to unify pipelines operating both at genomic (FASTQ, FASTA, SEQ) and proteomic levels, with an open license and free of any dependency. This toolkit includes information theory-based tools for reference-free and reference-based data compression applied to data analysis. Among many applications, this toolkit supports the creation of workflows for identification of metagenomic composition in FASTQ reads, detection and visualisation of genomic rearrangements, mapping and visualisation of variation, localisation of low complexity regions, or simulation of sequences with specific SNP and structural variant rates. The toolkit was designed for Unix/Linux-based systems, built for ultra-fast computations. It supports pipes for easy integration with the sub-programs as well as external tools. GTO works as *LEGOs^™^*, since it allows the construction of multiple pipelines with many combinations. We support the toolkit with a detailed manual and a website with several examples for fast learning.

Due to the variety and distribution of the given tools and their tight interconnection using the command line with pipes, the toolkit is an excellent platform for scientists as well as for empowering students to progress to the scientific aspects of bioinformatics analysis efficiently. Therefore, without the need to install multiple programs, dependencies, and read different manuals or licenses, it is possible to maintain an easy-to-follow connection with all the phases of each pipeline application.

## 2. The Toolkit

GTO is a powerful toolkit composed of more than 75 tools with particular focus on genomics and proteomics, following an integrative and flexible design between the tools. These tools aim key features such as they are easy to use, to compile, to improve and specially designed for work in Unix/Linux command line. These tools can be used isolated, or combined as one, forming execution workflows. This is technically possible due to the two streams used for the computation, namely the standard input and output. Furthermore, the tools’ aggregation is possible with mechanisms for inter-process communication using message passing, provided by the Unix operating system. This creates a chain of processes in which the output of each process is passed directly as input to its subsequent, as shown in the following example:

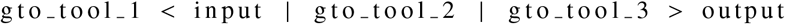

GTO includes tools for information display, randomisation, edition, conversion, extraction, search, calculation, compression, simulation and visualisation. The toolkit is prepared to deal with extensive datasets, typically in the scale of Gigabytes or Terabytes (but not limited). Furthermore, the toolkit contains three main groups of tools according to its characteristic: Genomics, Proteomics, and General purpose. The genomics group is subdivided in: FASTQ, FASTA, SEQ (genomic sequences); while the proteomics contains AA (amino acid); the general-purpose tools can be applied to any format sequence.

### 2.1. Genomics

The toolkit allows the data conversion between different formats aiming the FASTQ or FASTA sequences. It also provides features for filtering and randomise DNA sequences, as well as for analytic purposes followed by simulations of generation and alteration nature. The SEQ cluster works directly with the DNA sequences without any standard format. These tools allow the data extraction, summarising, classification and mathematical operations in the information theory field. Among many examples, which are better described at the supporting website and manual, the toolkit allows the preparations of the reads, namely filtering and trimming, the automatic construction of nucleotide reference databases, and comparative genomics.

### 2.2. Proteomics

The toolkit has a specific cluster of tools designed to group, compress, and analyse amino acid sequences. These tools allow proteomic analysis based on the amino acids properties, such as electric charge (positive and negative), uncharged side chains, hydrophobic side chains and special cases. The toolkit permits to find approximate amino acid sequences and perform comparative proteomics analysis.

### 2.3. General Purpose

This set of tools are complementary to the genomics and proteomics tools, in which they were not designed to work in a specific field, but to assist the pipelines composed by the previously described subsets. These tools provide operations in the symbolical domain, including reversion, segmentation, and permutation; while in the numerical domain it contains tools that contain low-pass filters (with multiple window types), sum, min and max operations over streams.

## 3. Validation

All the tools in the toolkit were tested with synthetic sequences aiming the individual validation. Therefore, the documented examples are easily replicable with the written tests. Besides the application of these tools in controlled environments, the toolkit was also used in several research workflows both as a primary and auxiliary tool. Several complete workflows are available in the repository, under the pipelines folder while an extensive description of the tool can be found at the manual. Further, we include some pipeline examples.

Another workflow example is the computation of bi-directional complexity profiles in any genomic or proteomic sequence [4]. These profiles can localise specific features in the sequences, namely low and high complexity sequences, inverted repeats regions, tandem duplications, among others. The construction of these profiles follow a pipeline constituted by many transformations (e.g. reversing, segmenting, inverting) and well as the use of specific low-pass filters after data compression applications [5]. Figure 1 depicts the complexity profiles of four human *Herpesvirus* whole genomes using the same scale, where redundant regions are highlighted with blue (below a Bps of one).

**Figure 1:**
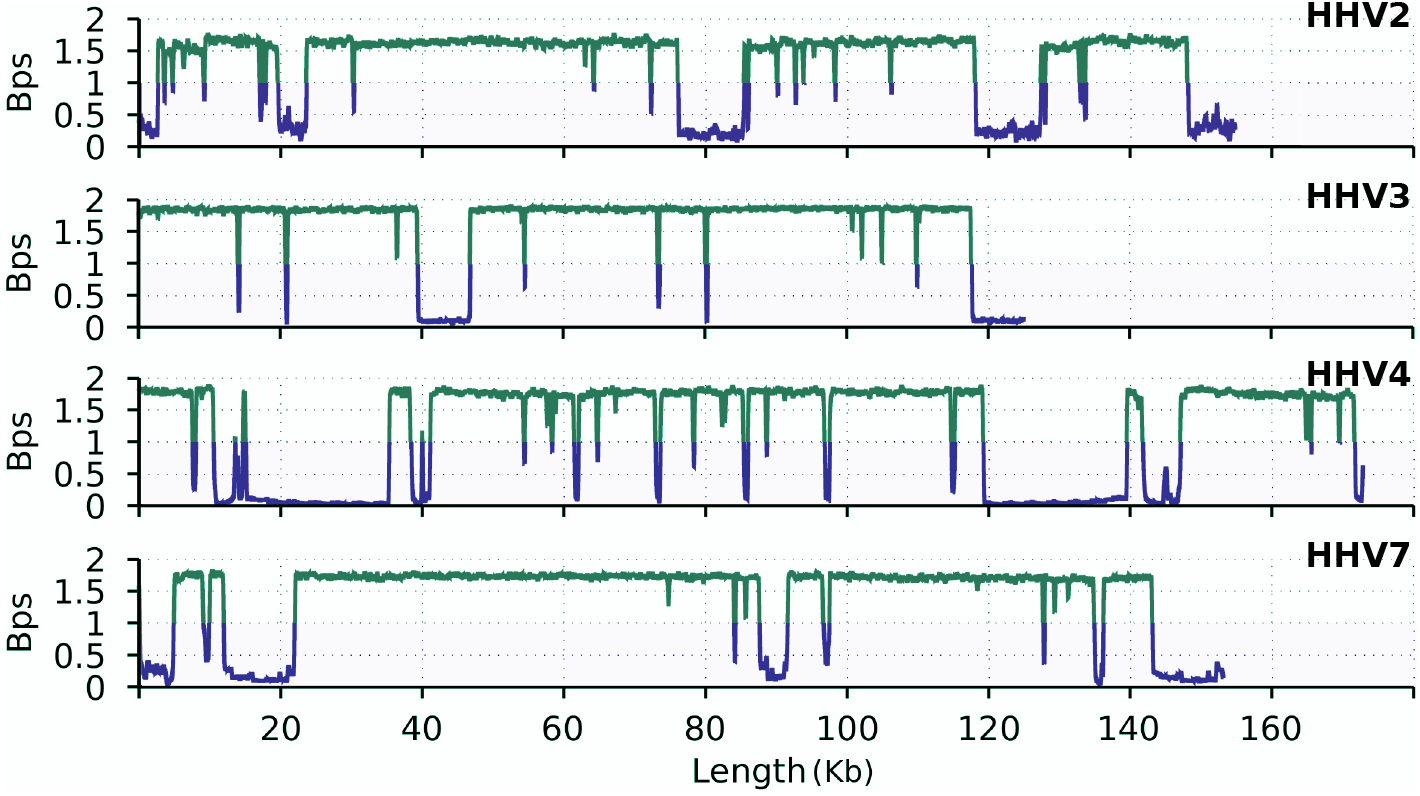
Bi-directional complexity profiles of four types of human *Herpesvirus* (HHV2, HHV3, HHV4 and HHV7) generated with GTO using the pipeline: `gto_complexity_profile_regions.sh`. Complexity values below one are highlighted with blue colour while the others with green. Bps stands for bits per symbol where lower values represent redundancy. The length is in Kb (Kilobases) and all profiles use the same scale.

GTO uses GeCo2 [6] and AC [7] compressors to estimate the local complexity of DNA and amino acid sequences, respectively. However, GTO is not limited to use these data compressors. For example, new models can be tested under this framework, namely with extended alphabets [8]. In general, any data compressor able to output local estimations can be used in the pipeline as an alternative [9].

Analogous to the complexity profiles for DNA sequences, an example using amino acid sequences is given in Figure 2. This example depicts a bi-directional complexity profile for the largest human protein sequence, titin. There are several regions with low complexity that usually are associated with specific characteristics, namely loops [10].

**Figure 2:**
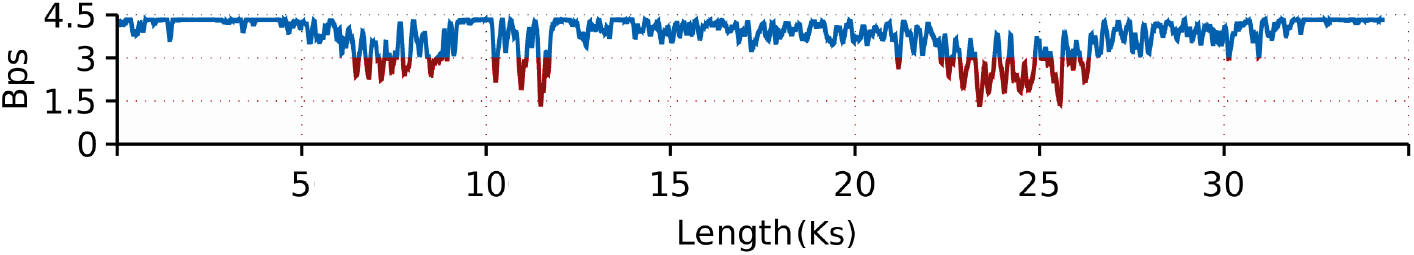
Bi-directional complexity profiles of human titin protein generated with GTO using the pipeline: `gto_proteins_complexity_profile_regions.sh`. Complexity values below three are highlighted with a red colour while the others with blue. Bps stands for bits per symbol where lower values represent redundancy. The length is in Ks (Kilosymbols).

Another example workflow is in the domain of comparative genomics, namely to map and visualise rearrangements. This workflow is completely automatic from the input of the sequences to the generation of an SVG image with the associated and transformed regions corresponding to the rearrangements. The pipeline applies smash technology [11, 12] for mapping the rearrangements using an alignment-free methodology [13]. For proving the efficiency of the mapping pipeline, we use another pipeline to generate two identical FASTA files with simulated rearrangements between them (`gto_simulate_rearragements.sh`). After, we load the two FASTA files into the mapping pipeline (`gto_map_rearrangements.sh`) where it outputs two files, one with the mapping positions and an SVG image depicting the mapped positions as it can be seen in Figure 3. All the rearrangements have been efficiently mapped with GTO according to the ground truth (< 1 second of computational time).

**Figure 3:**
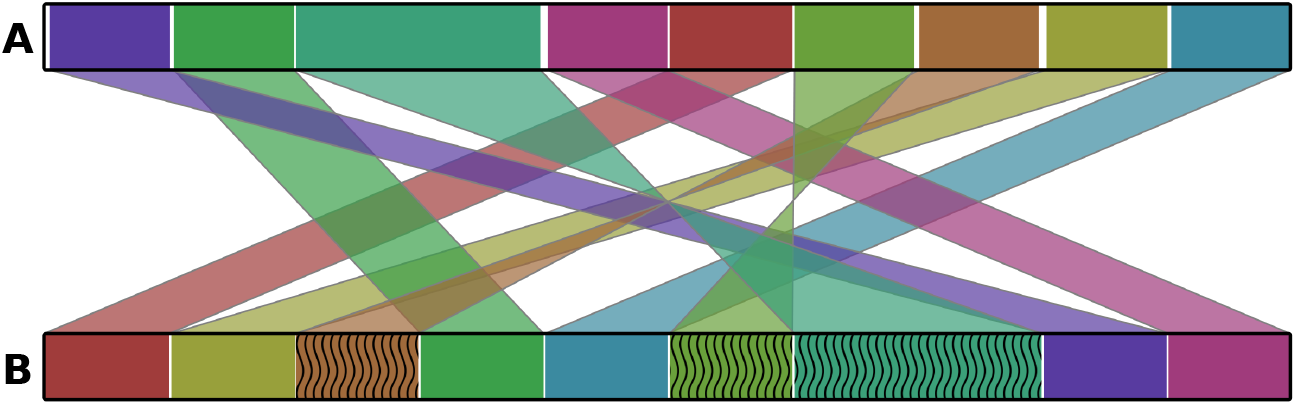
Rearrangements map generated with GTO using the pipeline: `gto_map_rearrangements.sh`. The length of both sequences (A and B) is 5 MB. Wave pattern stands for inverted repeated regions.

Analogous to the rearrangements map pipeline, for mapping at proteomic level, we consider the NAV2 HUMAN Neuron navigator 2 and the neuron navigator 2 isoform X15 of *Macaca mulatta* proteins. Although there are many examples under the proteome evolution [14], these are protein sequences under the consideration of identical scale [15]. Additionally, we shuffled the *Macaca mulatta* proteins using a block size of 300 amino acids. Figure 4 depicts the proteins map after running the pipeline (`gto_map_rearrangements_proteins.sh`). Despite a low level of dissimilarity of the sequences with an additional pseudo-random permutation of blocks of 300 symbols, all the regions have been efficiently mapped with GTO (< 1 second of computational time).

**Figure 4:**
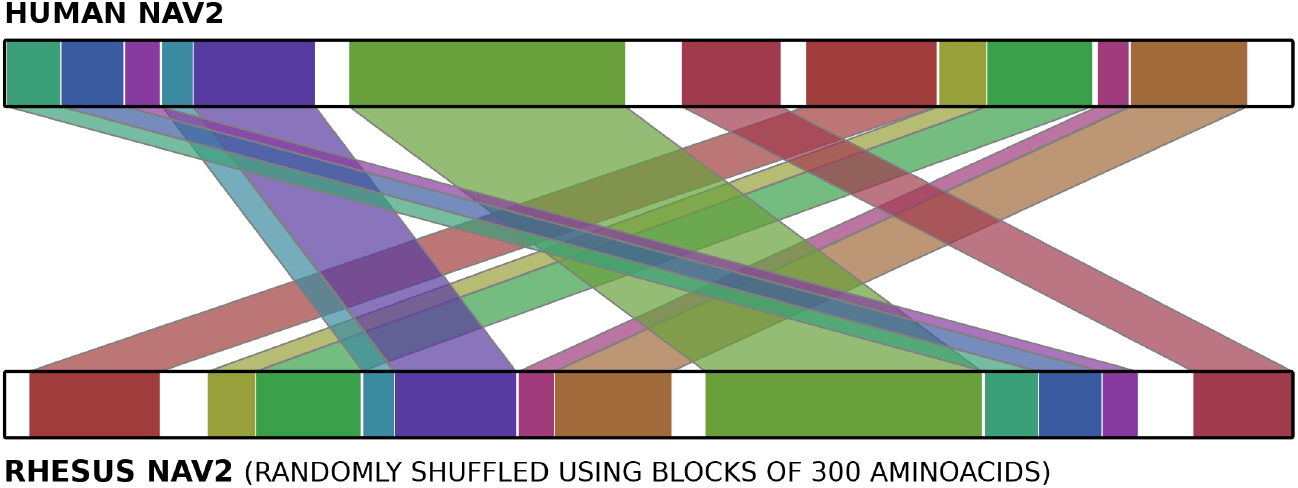
Rearrangements map generated with GTO using the pipeline: `gto_map_rearrangements_proteins.sh`.

A final workflow example is the full automatic metagenomic identification of viral (or any other) content in FASTQ reads. This includes the filtering and trimming of the reads, mapping, and sensitive identification of the most representative genomes, under a ranking of abundance. In this particular example, we generate a semi-synthetic viral dataset containing several real viruses with applied degrees of substitutions and block permutations shuffled with synthetic noisy DNA. This dataset is generated using the `gto_create_viral_dataset.sh` pipeline.

The intention is to perform a metagenomic analysis on this dataset without informing the program what organisms are contained in the sample since the program needs to infer the results. Then, we compare the results with the ground truth. If the results are similar to the ground truth, then the methodology is validated. For the purpose, GTO uses falcon-meta technology [16, 17] that relies on assembly-free and alignment-free comparison of each reference according to the whole reads. The dataset contains synthetic reads (uniform distribution) merged with the following viruses with the respective modifications:

- B19V: two *Parvovirus*, one with 1% of editions and other with permuted blocks of 500 bases (GID: AY386330.1);
- HHV2: one human *Herpesvirus* 2 with permuted blocks of size 100 bases (GID: JN561323.2);
- HHV3: one human *Herpesvirus* 3 (GID: X04370.1);
- HHV4: two human *Herpesvirus* 4, one with permuted blocks of 300 bases (GID: DQ279927.1);
- TTV: one human *Torque teno* virus with 5% of editions (GID: AB041963.1);
- HPV: one human *Papillomavirus* with 5% of editions and permuted blocks of 300 bases (GID: MG921180.1).

After the merging of all FASTA sequences, ART [18] was used to generate the paired end FASTQ reads. In the meanwhile, another workflow example was used to create the viral database (`gto_build_dbs.sh`). Then, the pipeline (`gto_metagenomics.sh`) ran obtaining the top output presented in Table 1.

**Table 1:** The eight most representative reference sequences according to the RS (Relative Similarity). The ID stands for the order of the top output, the length for the size of the reference genome, and the GID for the sequence global identifier.

| ID | Length | RS (%) | Reference GID | Virus name |
| --- | --- | --- | --- | --- |
| 1 | 124884 | 97.767 | X04370.1 | HHV3 |
| 2 | 5596 | 96.603 | AY386330.1 | B19V |
| 3 | 172764 | 94.143 | DQ279927.1 | HHV4 |
| 4 | 154675 | 81.400 | JN561323.2 | HHV2 |
| 5 | 154746 | 80.153 | Z86099.2 | HHV2 |
| 6 | 2785 | 78.300 | AB041963.1 | TTV |
| 7 | 7372 | 71.445 | MG921180.1 | HPV |
| 8 | 549 | 47.591 | AY034056.1 | PHV3-BALF1-gene |

We can conclude that despite the noise, editions, and permutations applied to real data, all the viruses have been efficiently identified with GTO, including the exact genotype (< 1.5 minutes of computational time).

## 4. Conclusion

We contribute with GTO, a toolkit to unify research pipelines, composed of distinct tools aiming at efficient combinations of them towards specific workflows. GTO’s efficient performance is due to the use of low-level programming languages, which increases the processing speed and decreases the RAM of addressing genomics and proteomics data. The source code and binaries of GTO are available in the GitHub repository and the Cobilab channel of Conda. Detailed information for each tool and the toolkit installation is also available at http://bioinformatics.ua.pt/gto.

## Supporting information

GTO Manual

## Funding

This work has received support from the NETDIAMOND project (POCI-01-0145-FEDER-016385), co-funded by Centro 2020 program, Portugal 2020, European Union. João Almeida is supported by FCT - Foundation for Science and Technology (national funds), grant SFRH/BD/147837/2019.

## Notes

https://bioinformatics.ua.pt/gto/

## References

[1] E. R. Mardis, DNA sequencing technologies: 2006–2016, Nature Protocols 12 (2017) 213.

[2] G. A. Van der Auwera, M. O. Carneiro, C. Hartl, R. Poplin, G. Del Angel, A. Levy-Moonshine, T. Jordan, K. Shakir, D. Roazen, J. Thibault, et al., From FASTQ data to high-confidence variant calls: the genome analysis toolkit best practices pipeline, Current Protocols in Bioinformatics 43 (2013) 11–10.

[3] D. Kessner, M. Chambers, R. Burke, D. Agus, P. Mallick, Proteowizard: open source software for rapid proteomics tools development, Bioinformatics 24 (2008) 2534–2536.

[4] A. J. Pinho, S. P. Garcia, D. Pratas, P. J. Ferreira, DNA sequences at a glance, PloS one 8 (2013) e79922.

[5] A. J. Pinho, D. Pratas, P. J. Ferreira, S. P. Garcia, Symbolic to numerical conversion of dna sequences using finite-context models, in: 2011 19th European Signal Processing Conference, IEEE, pp. 2024–2028.

[6] D. Pratas, M. Hosseini, A. J. Pinho, GeCo2: an optimized tool for lossless compression and analysis of DNA sequences, in: International Conference on Practical Applications of Computational Biology & Bioinformatics, Springer, pp. 137–145.

[7] M. Hosseini, D. Pratas, A. J. Pinho, AC: a compression tool for amino acid sequences, Interdisciplinary Sciences: Computational Life Sciences 11 (2019) 68–76.

[8] J. M. Carvalho, S. Brás, D. Pratas, J. Ferreira, S. C. Soares, A. J. Pinho, Extended-alphabet finite-context models, Pattern Recognition Letters 112(2018)49–55.

[9] M. Hosseini, D. Pratas, A. Pinho, A survey on data compression methods for biological sequences, Information 7 (2016) 56.

[10] G. Agüero-Chapin, H. González-Díaz, R. Molina, J. Varona-Santos, E. Uriarte, Y. Gonález-Díaz, Novel 2D maps and coupling numbers for protein sequences. The first QSAR study of polygalacturonases; isolation and prediction of a novel sequence from Psidium guajava L., FEBS letters 580 (2006) 723–730.

[11] D. Pratas, R. M. Silva, A. J. Pinho, P. J. Ferreira, An alignment-free method to find and visualise rearrangements between pairs of dna sequences, Scientific reports 5 (2015) 10203.

[12] M. Hosseini, D. Pratas, B. Morgenstern, A. J. Pinho, Smash++: an alignment-free and memory-efficient tool to find genomic rearrangements, bioRxiv (2019).

[13] A. Zielezinski, H. Z. Girgis, G. Bernard, C.-A. Leimeister, K. Tang, T. Dencker, A. K. Lau, S. Röhling, J. Choi, M. S. Waterman, et al., Benchmarking of alignment-free sequence comparison methods, BioRxiv (2019) 611137.

[14] S. K. Forslund, M. Kaduk, E. L. Sonnhammer, Evolution of protein domain architectures, in: Evolutionary Genomics, Springer, 2019, pp. 469–504.

[15] B. Rost, Twilight zone of protein sequence alignments, Protein engineering 12 (1999) 85–94.

[16] D. Pratas, A. J. Pinho, R. M. Silva, J. M. Rodrigues, M. Hosseini, T. Caetano, P. J. Ferreira, FALCON: a method to infer metagenomic composition of ancient DNA, BioRxiv (2018) 267179.

[17] D. Pratas, A. J. Pinho, Metagenomic composition analysis of sedimentary ancient DNA from the Isle of Wight, in: 2018 26th European Signal Processing Conference (EUSIPCO), IEEE, pp. 1177–1181.

[18] W. Huang, L. Li, J. R. Myers, G. T. Marth, ART: a next-generation sequencing read simulator, Bioinformatics 28 (2011) 593–594.

