## Supplementary material for "GTO: a toolkit to unify pipelines in genomic and proteomic research": GTO Manual

GTØ

### The Genomics Toolkit

João Rafael Almeida [1,2,3]
Armando José Pinho [1,2]
José Luís Oliveira [1,2]
Olga Fajarda [1,2]
Diogo Pratas [1,2,4]

[1]Institute of Electronics and Informatics Engineering of Aveiro, University of Aveiro, Aveiro, Portugal
[2]Department of Electronics, Telecommunications and Informatics, University of Aveiro, Aveiro, Portugal
[3]Department of Information and Communications Technologies, University of A Coruña, A Coruña, Spain
[4]Department of Virology, University of Helsinki, Helsinki, Finland

Version 1.4

**Bibliography** **107**

### Chapter 1

### Introduction

Recent advances in DNA sequencing, specifically in next-generation sequencing (NGS), revolutionised the field of genomics, making possible the generation of large amounts of sequencing data very rapidly and at substantially low cost [1]. This new technology also brought with it several challenges, namely in what concerns the analysis, storage, and transmission of the generated sequences [2, 3]. As a consequence, several specialised tools were developed throughout the years in order to deal with these challenges.

Firstly, the storage of the raw data generated by NGS experiments is possible by using several file formats, the FASTQ and FASTA are the most commonly used [4]. FASTQ is an extension of the FASTA format, that besides the nucleotide sequence, also stores associated per base quality score and it is considered the standard format for sequencing data storage and exchange [5].

Regarding the analysis and manipulation of these sequencing data files many software applications emerged, including `fqtools` [6], `FASTX-Toolkit` [7], `GALAXY` [8], `GATK` [9], `MEGA` [10], `SeqKit` [11], among others. `Fqtools` is a suite of tools to view, manipulate and summarise FASTQ data. This software also identifies invalid FASTQ files [6]. `GALAXY`, in its turn, is an open, web-based scientific platform for analysing genomic data [12]. This platform integrates several specialised sets of tools, e.g. for manipulating FASTQ files [13]. `FASTX-Toolkit` is a collection of command-line tools to process FASTA and FASTQ files. This toolkit is available in two forms: as a command-line, or integrated into the web-based platform GALAXY [7]. `SeqKit` is another toolkit used to process FASTA and FASTQ files and is available for all major operating systems [11]. The Genome Analysis Toolkit (`GATK`) was designed as a structured programming framework to simplify the development of analysis tools. However, nowadays, it is a suite of tools focused on variant discovering and genotyping [14]. More towards the evolutionary perspectives, Molecular Evolutionary Genetics Analysis (`MEGA`) software provides tools to analyse DNA and protein sequences statistically [15]. Several of these frameworks lack on variety, namely the ability to perform multiple tasks using only one toolkit.

Compression is another important aspect when dealing with high-throughput sequencing data, as it reduces storage space and accelerates data transmission. A survey on DNA compressors and amino acid sequence compression can be found in [16]. Currently, the DNA sequence compressors HiRGC [17], iDoComp [18], GeCo [19], and GDC [20] are considered to have the best performance [21]. Of these four approaches, GeCo is the only one that can be used for reference-free and reference-based compression. Furthermore, GeCo can be used as an analysis tool to determine absolute measures for many distance computations and local measures [19].

Amino acid sequences are known to be very hard to compress [22], however, Hosseini et al. [23] recently developed AC, a state-of-the-art for lossless amino acid sequence compression. In [24] the authors compared the performance of AC, in terms of bit-rate, to several general-purpose lossless compressors and several protein compressors, using different proteomes. They concluded that in average AC provides the best bit-rates.

Another relevant subject is genomic data simulation. Read simulations tools are fundamental for the development, testing and evaluation of methods and computational tools [25, 26]. Despite the availability of a large number of real sequence reads, read simulation data is necessary due to the inability to know the ground truth of real data [27]. Escalona *et al.* [28], recently, reviewed 23 NGS simulation tools. XS [29], a FASTQ read simulation tool, stands out in relation to the other 22 simulation tools because it is the only one that does not need a reference sequence. Furthermore, XS is the only open-source tool for simulation of FASTQ reads produced by the four most used sequencing machines, Roche-454, Illumina, ABI SOLiD and Ion Torrent.

Although a large number of tools are available for analysing, compressing, and simulation, these tools are specialised in only a specific task. Besides, in many cases the output of one tool cannot be used directly as input for another tool, e.g. the output of a simulation tool cannot always be used directly as input for an analysis tool. Thus, unique software that includes several specialised tools is necessary.

In this document, we describe `GTO`, a complete toolkit for genomics and proteomics, namely for FASTQ, FASTA and SEQ formats, with many complementary tools. The toolkit is for Unix-based systems, built for ultra-fast computations. `GTO` supports pipes for easy integration with the sub-programs belonging to `GTO` as well as external tools. `GTO` works as *LEGOs*, since it allows the construction of multiple pipelines with many combinations.

`GTO` includes tools for information display, randomisation, edition, conversion, extraction, search, calculation, compression, simulation and visualisation. `GTO` is prepared to deal with very large datasets, typically in the scale of Gigabytes or Terabytes (but not limited). The complete toolkit is an optimised command-line version, using the prefix "`gto_`" followed by the suffix with the respective name of the program. `GTO` is implemented in `C` language and it is available, under the MIT license, at:

```
http://bioinformatics.ua.pt/gto
```

#### 1.1 Installation

To install `GTO` through the GitHub repository:

```
git clone https://github.com/cobilab/gto.git
cd gto/src/
make
```

Or by installing them directly using the Cobilab channel from Conda:

```
conda install -c cobilab gto --yes
```

#### 1.2 Testing

The examples provided in this document are available in the repository. Therefore, each example can be easily reproduced, which it will also test and validate each tool. To replicate those tests, it can be done in two different ways:

- Running one test for a specific tool:

  ```
  cd gto/tester/gto_{tool}
  sh runExample.sh
  ```

- Running the batch of tests for all the tools:

  ```
  cd gto/tester/
  sh runAllTests.sh
  ```

Some of these tests require internet connection to download external files and it will create new files.

#### 1.3 License

The license is **MIT**. In resume, it is a short and simple permissive license with conditions only requiring preservation of copyright and license notices. Licensed works, modifications, and larger works may be distributed under different terms and without source code.

**Permissions**:

- commercial use;
- modification;
- distribution;
- private use.

**Limitations**:

- liability;
- warranty.

**Conditions**:

- License and copyright notice.

For details on the license, consult: `https://opensource.org/licenses/MIT`.

### Chapter 2

### FASTQ tools

The toolkit has a set of tools dedicated to manipulating FASTQ files. Some of these tools allow the data conversion to/from different formats, i. e., there are tools designed to convert a FASTQ file into a sequence or a FASTA/Multi-FASTA format, or converting DNA in some of those formats to FASTQ.

There are also tools for data manipulation in this format, which are designed to exclude 'N', remove low quality scored reads, following different metrics and randomize DNA sequences. Succeeding the manipulation, it is also possible to perform analyses over these files, simulations and mutations. The current available tools for FASTQ format analysis and manipulation include:

1. `gto_fastq_to_fasta`: it converts a FASTQ file format to a pseudo FASTA file.
2. `gto_fastq_to_mfasta`: it converts a FASTQ file format to a pseudo Multi-FASTA file.
3. `gto_fastq_exclude_n`: it discards the FASTQ reads with the minimum number of "N" symbols.
4. `gto_fastq_extract_quality_scores`: it extracts all the quality-scores from FASTQ reads.
5. `gto_fastq_info`: it analyses the basic information of FASTQ file format.
6. `gto_fastq_maximum_read_size`: it filters the FASTQ reads with the length higher than the value defined.
7. `gto_fastq_minimum_quality_score`: it discards reads with average quality-score below of the defined.
8. `gto_fastq_minimum_read_size`: it filters the FASTQ reads with the length smaller than the value defined.
9. `gto_fastq_rand_extra_chars`: it substitues in the FASTQ files, the DNA sequence the outside ACGT chars by random ACGT symbols.
10. `gto_fastq_from_seq`: it converts a genomic sequence to pseudo FASTQ file format.
11. `gto_fastq_mutate`: it creates a synthetic mutation of a FASTQ file given specific rates of mutations, deletions and additions.

12. gto_fastq_split: it splits Paired End files according to the direction of the strand ('/1' or '/2').

13. gto_fastq_pack: it packages each FASTQ read in a single line.

14. gto_fastq_unpack: it unpacks the FASTQ reads packaged using the gto_fastq_pack tool.

15. gto_fastq_quality_score_info: it analyses the quality-scores of a FASTQ file.

16. gto_fastq_quality_score_min: it analyses the minimal quality-scores of a FASTQ file.

17. gto_fastq_quality_score_max: it analyses the maximal quality-scores of a FASTQ file.

18. gto_fastq_cut: it cuts read sequences in a FASTQ file.

19. gto_fastq_minimum_local_quality_score_forward: it filters the reads considering the quality score average of a defined window size of bases.

20. gto_fastq_minimum_local_quality_score_reverse: it filters the reverse reads, considering the average window size score defined by the bases.

21. gto_fastq_xs: it is a skilled FASTQ read simulation tool, flexible, portable and tunable in terms of sequence complexity.

22. gto_fastq_clust_reads: it agroups reads and creates an index file.

23. gto_fastq_complement: it replaces the ACGT bases with their complements in a FASTQ file format.

24. gto_fastq_reverse: it reverses the ACGT bases order for each read in a FASTQ file format.

25. gto_fastq_variation_map: it identifies the variation that occours in the sequences relative to the reads or a set of reads.

26. gto_fastq_variation_filter: it filters and segments the regions of singularity from the output of gto_fastq_variation_map.

27. gto_fastq_variation_visual: it depites the regions of singularity using the output from gto_fastq_variation_ into an SVG image.

28. gto_fastq_falcon: it measures similarity between any FASTQ file, independently from the size, against any multi-FASTA database.

#### 2.1 Program gto_fastq_to_fasta

The gto_fastq_to_fasta converts a FASTQ file format to a pseudo FASTA file. However, it does not align the sequence. Also, it extracts the sequence and adds a pseudo header.
For help type:

```
./gto_fastq_to_fasta -h
```

In the following subsections, we explain the input and output paramters.

##### Input parameters

The `gto_fastq_to_fasta` program needs two streams for the computation, namely the input and output standard. The input stream is a FASTQ file.
The attribution is given according to:

```
Usage: ./gto_fastq_to_fasta [options] [[--] args]
   or: ./gto_fastq_to_fasta [options]

It converts a FASTQ file format to a pseudo FASTA file.
It does NOT align the sequence.
It extracts the sequence and adds a pseudo header.

    -h, --help            show this help message and exit

Basic options
    < input.fastq         Input FASTQ file format (stdin)
    > output.fasta        Output FASTA file format (stdout)

Example: ./gto_fastq_to_fasta < input.fastq > output.fasta
```

An example of such an input file is:

```
@SRR001666.1 071112_SLXA-EAS1_s_7:5:1:817:345 length=72
GGGTGATGGCCGCTGCCGATGGCGTCAAATCCCACCAAGTTACCCTTAACAACTTAAGGGTTTTCAAATAGA
+SRR001666.1 071112_SLXA-EAS1_s_7:5:1:817:345 length=72
IIIIIIIIIIIIIIIIIIIIIIIIIIIIII9IG9ICIIIIIIIIIIIIIIIIIIIIDIIIIIII>IIIIII/
@SRR001666.2 071112_SLXA-EAS1_s_7:5:1:801:338 length=72
GTTCAGGGATACGACGTTTGTATTTTAAGAATCTGAAGCAGAAGTCGATGATAATACGCGTCGTTTTATCAT
+SRR001666.2 071112_SLXA-EAS1_s_7:5:1:801:338 length=72
IIIIIIIIIIIIIIIIIIIIIIIIIIIIIIII6IBIIIIIIIIIIIIIIIIIIIIIIIGII>IIIII-I)8I
```

##### Output

The output of the `gto_fastq_to_fasta` program a FASTA file.
Using the input above, an output example for this is the following:

```
> Computed with Fastq2Fasta
GGGTGATGGCCGCTGCCGATGGCGTCAAATCCCACCAAGTTACCCTTAACAACTTAAGGGTTTTCAAATAGA
GTTCAGGGATACGACGTTTGTATTTTAAGAATCTGAAGCAGAAGTCGATGATAATACGCGTCGTTTTATCAT
```

#### 2.2 Program gto_fastq_to_mfasta

The `gto_fastq_to_mfasta` onverts a FASTQ file format to a pseudo Multi-FASTA file. However, it does not align the sequence. Also, it extracts the sequence and adds a pseudo header.

For help type:

```
./gto_fastq_to_mfasta -h
```

In the following subsections, we explain the input and output paramters.

##### Input parameters

The `gto_fastq_to_mfasta` program needs two streams for the computation, namely the input and output standard. The input stream is a FASTQ file.
The attribution is given according to:

```
Usage: ./gto_fastq_to_mfasta [options] [[--] args]
   or: ./gto_fastq_to_mfasta [options]

It converts a FASTQ file format to a pseudo Multi-FASTA file.
It does NOT align the sequence.
It extracts the sequence and adds each header in a Multi-FASTA format.

    -h, --help             show this help message and exit

Basic options
    < input.fastq          Input FASTQ file format (stdin)
    > output.mfasta        Output Multi-FASTA file format (stdout)

Example: ./gto_fastq_to_mfasta < input.fastq > output.mfasta
```

An example of such an input file is:

```
@SRR001666.1 071112_SLXA-EAS1_s_7:5:1:817:345 length=72
GGGTGATGGCCGCTGCCGATGGCGTCAAATCCCACCAAGTTACCCTTAACAACTTAAGGGTTTTCAAATAGA
+SRR001666.1 071112_SLXA-EAS1_s_7:5:1:817:345 length=72
IIIIIIIIIIIIIIIIIIIIIIIIIIIIII9IG9ICIIIIIIIIIIIIIIIIIIIIDIIIIIII>IIIIII/
@SRR001666.2 071112_SLXA-EAS1_s_7:5:1:801:338 length=72
GTTCAGGGATACGACGTTTGTATTTTAAGAATCTGAAGCAGAAGTCGATGATAATACGCGTCGTTTTATCAT
+SRR001666.2 071112_SLXA-EAS1_s_7:5:1:801:338 length=72
IIIIIIIIIIIIIIIIIIIIIIIIIIIIIIII6IBIIIIIIIIIIIIIIIIIIIIIIIGII>IIIII-I)8I
```

##### Output

The output of the `gto_fastq_to_mfasta` program a Multi-FASTA file.
Using the input above, an output example for this is the following:

```
>SRR001666.1 071112_SLXA-EAS1_s_7:5:1:817:345 length=72
GGGTGATGGCCGCTGCCGATGGCGTCAAATCCCACCAAGTTACCCTTAACAACTTAAGGGTTTTCAAATAGA
>SRR001666.2 071112_SLXA-EAS1_s_7:5:1:801:338 length=72
GTTCAGGGATACGACGTTTGTATTTTAAGAATCTGAAGCAGAAGTCGATGATAATACGCGTCGTTTTATCAT
```

#### 2.3 Program gto_fastq_exclude_n

The `gto_fastq_exclude_n` discards the FASTQ reads with the minimum number of "N" symbols. Also, if present, it will erase the second header (after +).
For help type:

```
./gto_fastq_exclude_n -h
```

In the following subsections, we explain the input and output paramters.

##### Input parameters

The `gto_fastq_exclude_n` program needs two streams for the computation, namely the input and output standard. The input stream is a FASTQ file.
The attribution is given according to:

```
Usage: ./gto_fastq_exclude_n [options] [[--] args]
   or: ./gto_fastq_exclude_n [options]

It discards the FASTQ reads with the minimum number of "N" symbols.
If present, it will erase the second header (after +).

    -h, --help            show this help message and exit

Basic options
    -m, --max=<int>       The maximum of of "N" symbols in the read
    < input.fastq         Input FASTQ file format (stdin)
    > output.fastq        Output FASTQ file format (stdout)

Example: ./gto_fastq_exclude_n -m <max> < input.fastq > output.fastq

Console output example :
<FASTQ non-filtered reads>
Total reads    : value
Filtered reads : value
```

An example of such an input file is:

```
@SRR001666.1 071112_SLXA-EAS1_s_7:5:1:817:345 length=72
GNNTGATGGCCGCTGCCGATGGCGNANAATCCCACCAANATACCCTTAACAACTTAAGGGTTNTCAAATAGA
+
IIIIIIIIIIIIIIIIIIIIIIIIIIIII9IG9ICIIIIIIIIIIIIIIIIIIIIDIIIIIII>IIIIII/
@SRR001666.2 071112_SLXA-EAS1_s_7:5:1:801:338 length=72
NTTCAGGGATACGACGNTTGTATTTTAAGAATCTGNAGCAGAAGTCGATGATAATACGCGNCGTTTTATCAN
+
IIIIIIIIIIIIIIIIIIIIIIIIIIIIIII6IBIIIIIIIIIIIIIIIIIIIIIIIGII>IIIII-I)8I
```

##### Output

The output of the `gto_fastq_exclude_n` program is a set of all the filtered FASTQ reads, followed by the execution report. The execution report only appears in the console.

Using the input above with the max value as 5, an output example for this is the following:

```
@SRR001666.2 071112_SLXA-EAS1_s_7:5:1:801:338 length=72
NTTCAGGGATACGACGNTTGTATTTTAAGAATCTGNAGCAGAAGTCGATGATAATACGCGNCGTTTTATCAN
+
IIIIIIIIIIIIIIIIIIIIIIIIIIIIII6IBIIIIIIIIIIIIIIIIIIIIIIIIGII>IIIII-I)8I
Total reads    : 2
Filtered reads : 1
```

#### 2.4 Program gto_fastq_extract_quality_scores

The `gto_fastq_extract_quality_scores` extracts all the quality-scores from FASTQ reads.
For help type:

```
./gto_fastq_extract_quality_scores -h
```

In the following subsections, we explain the input and output paramters.

##### Input parameters

The `gto_fastq_extract_quality_scores` program needs two streams for the computation, namely the input and output standard. The input stream is a FASTQ file.
The attribution is given according to:

```
Usage: ./gto_fastq_extract_quality_scores [options] [[--] args]
   or: ./gto_fastq_extract_quality_scores [options]

It extracts all the quality-scores from FASTQ reads.

    -h, --help            show this help message and exit

Basic options
    < input.fastq         Input FASTQ file format (stdin)
    > output.fastq        Output FASTQ file format (stdout)

Example: ./gto_fastq_extract_quality_scores < input.fastq > output.fastq

Console output example:
<FASTQ quality scores>
Total reads          : value
Total Quality-Scores : value
```

An example of such an input file is:

```
@SRR001666.1 071112_SLXA-EAS1_s_7:5:1:817:345 length=72
GGGTGATGGCCGCTGCCGATGGCGTCAAATCCCACCAAGTTACCCTTAACAACTTAAGGGTTTTCAAATAGA
+SRR001666.1 071112_SLXA-EAS1_s_7:5:1:817:345 length=72
IIIIIIIIIIIIIIIIIIIIIIIIIIIIII9IG9ICIIIIIIIIIIIIIIIIIIIIDIIIIIII>IIIIII/
@SRR001666.2 071112_SLXA-EAS1_s_7:5:1:801:338 length=72
GTTCAGGGATACGACGTTTGTATTTTAAGAATCTGAAGCAGAAGTCGATGATAATACGCGTCGTTTTATCAT
```

```
+SRR001666.2 071112_SLXA-EAS1_s_7:5:1:801:338 length=72
IIIIIIIIIIIIIIIIIIIIIIIIIIIIII6IBIIIIIIIIIIIIIIIIIIIIIIIGII>IIIII-I)8I
```

##### Output

The output of the `gto_fastq_extract_quality_scores` program is a set of all the quality scores from the FASTQ reads, followed by the execution report. The execution report only appears in the console.
Using the input above, an output example for this is the following:

```
IIIIIIIIIIIIIIIIIIIIIIIIIIIIII9IG9ICIIIIIIIIIIIIIIIIIIIIDIIIIIII>IIIIII/
IIIIIIIIIIIIIIIIIIIIIIIIIIIIII6IBIIIIIIIIIIIIIIIIIIIIIIIGII>IIIII-I)8I
Total reads          : 2
Total Quality-Scores : 144
```

#### 2.5 Program gto_fastq_info

The `gto_fastq_info` analyses the basic information of FASTQ file format.
For help type:

```
./gto_fastq_info -h
```

In the following subsections, we explain the input and output paramters.

##### Input parameters

The `gto_fastq_info` program needs two streams for the computation, namely the input and output standard. The input stream is a FASTQ file.
The attribution is given according to:

```
Usage: ./gto_fastq_info [options] [[--] args]
   or: ./gto_fastq_info [options]

It analyses the basic information of FASTQ file format.

    -h, --help            show this help message and exit

Basic options
    < input.fastq         Input FASTQ file format (stdin)
    > output              Output read information (stdout)

Example: ./gto_fastq_info < input.fastq > output

Output example:
Total reads     : value
Max read length : value
Min read length : value
Min QS value    : value
Max QS value    : value
QS range        : value
```

An example of such an input file is:

```
@SRR001666.1 071112_SLXA-EAS1_s_7:5:1:817:345 length=72
GGGTGATGGCCGCTGCCGATGGCGTCAAATCCCACCAAGTTACCCTTAACAACTTAAGGGTTTTCAAATAGA
+SRR001666.1 071112_SLXA-EAS1_s_7:5:1:817:345 length=72
IIIIIIIIIIIIIIIIIIIIIIIIIIIIII9IG9ICIIIIIIIIIIIIIIIIIIIIDIIIIIII>IIIIII/
@SRR001666.2 071112_SLXA-EAS1_s_7:5:1:801:338 length=72
GTTCAGGGATACGACGTTTGTATTTTAAGAATCTGAAGCAGAAGTCGATGATAATACGCGTCGTTTTATCAT
+SRR001666.2 071112_SLXA-EAS1_s_7:5:1:801:338 length=72
IIIIIIIIIIIIIIIIIIIIIIIIIIIIIIII6IBIIIIIIIIIIIIIIIIIIIIIIIGII>IIIII-I)8I
```

##### Output

The output of the `gto_fastq_info` program is a set of information related to the file read.
Using the input above, an output example for this is the following:

```
Total reads     : 2
Max read length : 72
Min read length : 72
Min QS value    : 41
Max QS value    : 73
QS range        : 33
```

#### 2.6 Program gto_fastq_maximum_read_size

The `gto_fastq_maximum_read_size` filters the FASTQ reads with the length higher than the value defined. For help type:

```
./gto_fastq_maximum_read_size -h
```

In the following subsections, we explain the input and output paramters.

##### Input parameters

The `gto_fastq_maximum_read_size` program needs two streams for the computation, namely the input and output standard. The input stream is a FASTQ file.
The attribution is given according to:

```
Usage: ./gto_fastq_maximum_read_size [options] [[--] args]
   or: ./gto_fastq_maximum_read_size [options]

It filters the FASTQ reads with the length higher than the value defined.
If present, it will erase the second header (after +).

    -h, --help            show this help message and exit

Basic options
```

```
    -s, --size=<int>      The maximum read length
    < input.fastq         Input FASTQ file format (stdin)
    > output.fastq        Output FASTQ file format (stdout)

Example: ./gto_fastq_maximum_read_size -s <size> < input.fastq > output.fastq

Console output example :
<FASTQ non-filtered reads>
Total reads    : value
Filtered reads : value
```

An example of such an input file is:

```
@SRR001666.1 071112_SLXA-EAS1_s_7:5:1:817:345 length=60
GGGTGATGGCCGCTGCCGATGGCGTCAAATCCCACCAAGTTACCCTTAACAACTTAAGGG
+
IIIIIIIIIIIIIIIIIIIIIIIIIIIIII9IG9ICIIIIIIIIIIIIIIIIIIIIDIII
@SRR001666.2 071112_SLXA-EAS1_s_7:5:1:801:338 length=72
GTTCAGGGATACGACGTTTGTATTTTAAGAATCTGAAGCAGAAGTCGATGATAATACGCGTCGTTTTATCAT
+
IIIIIIIIIIIIIIIIIIIIIIIIIIIIIIII6IBIIIIIIIIIIIIIIIIIIIIIIIGII>IIIII-I)8I
```

##### Output

The output of the `gto_fastq_maximum_read_size` program is a set of all the filtered FASTQ reads, followed by the execution report. The execution report only appears in the console.
Using the input above with the size values as 60, an output example for this is the following:

```
@SRR001666.1 071112_SLXA-EAS1_s_7:5:1:817:345 length=60
GGGTGATGGCCGCTGCCGATGGCGTCAAATCCCACCAAGTTACCCTTAACAACTTAAGGG
+
IIIIIIIIIIIIIIIIIIIIIIIIIIIIII9IG9ICIIIIIIIIIIIIIIIIIIIIDIII
Total reads    : 2
Filtered reads : 1
```

#### 2.7 Program gto_fastq_minimum_quality_score

The `gto_fastq_minimum_quality_score` discards reads with average quality-score below of the defined.
For help type:

```
./gto_fastq_minimum_quality_score -h
```

In the following subsections, we explain the input and output paramters.

##### Input parameters

The `gto_fastq_minimum_quality_score` program needs two streams for the computation, namely the input and output standard. The input stream is a FASTQ file.
The attribution is given according to:

```
Usage: ./gto_fastq_minimum_quality_score [options] [[--] args]
   or: ./gto_fastq_minimum_quality_score [options]

It discards reads with average quality-score below value.

    -h, --help             show this help message and exit

Basic options
    -m, --min=<int>        The minimum average quality-score (Value 25 or 30 is commonly used)
    < input.fastq          Input FASTQ file format (stdin)
    > output.fastq         Output FASTQ file format (stdout)

Example: ./gto_fastq_minimum_quality_score -m <min> < input.fastq > output.fastq

Console output example:
<FASTQ non-filtered reads>
Total reads    : value
Filtered reads : value
```

An example of such an input file is:

```
@SRR001666.1 071112_SLXA-EAS1_s_7:5:1:817:345 length=72
GGGTGATGGCCGCTGCCGATGGCGTCAAATCCCACCAAGTTACCCTTAACAACTTAAGGGTTTTCAAATAGA
+SRR001666.1 071112_SLXA-EAS1_s_7:5:1:817:345 length=72
IIIIIIIIIIIIIIIIIIIIIIIIIIIIII9IG9ICIIIIIIIIIIIIIIIIIIIIDIIIIIII>IIIIII/
@SRR001666.2 071112_SLXA-EAS1_s_7:5:1:801:338 length=72
GTTCAGGGATACGACGTTTGTATTTTAAGAATCTGAAGCAGAAGTCGATGATAATACGCGTCGTTTTATCAT
+SRR001666.2 071112_SLXA-EAS1_s_7:5:1:801:338 length=72
54599<>77977==6=?I6IBI::3334423552167799>>><<<@@A@BBCDGGBFFH>IIIII-I)8I
```

##### Output

The output of the `gto_fastq_minimum_quality_score` program is a set of all the filtered FASTQ reads, followed by the execution report.
Using the input above with the minimum averge value as 30, an output example for this is the following:

```
@SRR001666.1 071112_SLXA-EAS1_s_7:5:1:817:345 length=72
GGGTGATGGCCGCTGCCGATGGCGTCAAATCCCACCAAGTTACCCTTAACAACTTAAGGGTTTTCAAATAGA
+
IIIIIIIIIIIIIIIIIIIIIIIIIIIIII9IG9ICIIIIIIIIIIIIIIIIIIIIDIIIIIII>IIIIII/
Total reads    : 2
Filtered reads : 1
```

#### 2.8 Program gto_fastq_minimum_read_size

The `gto_fastq_minimum_read_size` filters the FASTQ reads with the length smaller than the value defined.
For help type:

```
./gto_fastq_minimum_read_size -h
```

In the following subsections, we explain the input and output paramters.

##### Input parameters

The `gto_fastq_minimum_read_size` program needs two streams for the computation, namely the input and output standard. The input stream is a FASTQ file.
The attribution is given according to:

```
Usage: ./gto_fastq_minimum_read_size [options] [[--] args]
   or: ./gto_fastq_minimum_read_size [options]

It filters the FASTQ reads with the length smaller than the value defined.
If present, it will erase the second header (after +).

    -h, --help            show this help message and exit

Basic options
    -s, --size=<int>      The minimum read length
    < input.fastq         Input FASTQ file format (stdin)
    > output.fastq        Output FASTQ file format (stdout)

Example: ./gto_fastq_minimum_read_size -s <size> < input.fastq > output.fastq

Console output example:
<FASTQ non-filtered reads>
Total reads    : value
Filtered reads : value
```

An example of such an input file is:

```
@SRR001666.1 071112_SLXA-EAS1_s_7:5:1:817:345 length=60
GGGTGATGGCCGCTGCCGATGGCGTCAAATCCCACCAAGTTACCCTTAACAACTTAAGGG
+SRR001666.1 071112_SLXA-EAS1_s_7:5:1:817:345 length=60
IIIIIIIIIIIIIIIIIIIIIIIIIIIIII9IG9ICIIIIIIIIIIIIIIIIIIIIDIII
@SRR001666.2 071112_SLXA-EAS1_s_7:5:1:801:338 length=72
GTTCAGGGATACGACGTTTGTATTTTAAGAATCTGAAGCAGAAGTCGATGATAATACGCGTCGTTTTATCAT
+SRR001666.2 071112_SLXA-EAS1_s_7:5:1:801:338 length=72
IIIIIIIIIIIIIIIIIIIIIIIIIIIIIIII6IBIIIIIIIIIIIIIIIIIIIIIIIGII>IIIII-I)8I
```

##### Output

The output of the `gto_fastq_minimum_read_size` program is a set of all the filtered FASTQ reads, followed by the execution report. The execution report only appears in the console.
Using the input above with the size values as 65, an output example for this is the following:

```
@SRR001666.2 071112_SLXA-EAS1_s_7:5:1:801:338 length=72
GTTCAGGGATACGACGTTTGTATTTTAAGAATCTGAAGCAGAAGTCGATGATAATACGCGTCGTTTTATCAT
+
IIIIIIIIIIIIIIIIIIIIIIIIIIIIIIII6IBIIIIIIIIIIIIIIIIIIIIIIIIGII>IIIII-I)8I
Total reads    : 2
Filtered reads : 1
```

#### 2.9 Program gto_fastq_rand_extra_chars

The `gto_fastq_rand_extra_chars` substitues in the FASTQ files, the DNA sequence the outside ACGT chars by random ACGT symbols.
For help type:

```
./gto_fastq_rand_extra_chars -h
```

In the following subsections, we explain the input and output paramters.

##### Input parameters

The `gto_fastq_rand_extra_chars` program needs two streams for the computation, namely the input and output standard. The input stream is a FASTQ file.
The attribution is given according to:

```
Usage: ./gto_fastq_rand_extra_chars [options] [[--] args]
   or: ./gto_fastq_rand_extra_chars [options]

It substitues in the FASTQ files, the DNA sequence the outside ACGT chars by random ACGT symbols.

    -h, --help            show this help message and exit

Basic options
    < input.fastq         Input FASTQ file format (stdin)
    > output.fastq        Output FASTQ file format (stdout)

Example: ./gto_fastq_rand_extra_chars < input.fastq > output.fastq
```

An example of such an input file is:

```
@SRR001666.1 071112_SLXA-EAS1_s_7:5:1:817:345 length=72
GNNTGATGGCCGCTGCCGATGGCGNANAATCCCACCAANATACCCTTAACAACTTAAGGGTTNTCAAATAGA
+SRR001666.1 071112_SLXA-EAS1_s_7:5:1:817:345 length=72
IIIIIIIIIIIIIIIIIIIIIIIIIIIIIII9IG9ICIIIIIIIIIIIIIIIIIIIIDIIIIIII>IIIIII/
@SRR001666.2 071112_SLXA-EAS1_s_7:5:1:801:338 length=72
NTTCAGGGATACGACGNTTGTATTTTAAGAATCTGNAGCAGAAGTCGATGATAATACGCGNCGTTTTATCAN
+SRR001666.2 071112_SLXA-EAS1_s_7:5:1:801:338 length=72
IIIIIIIIIIIIIIIIIIIIIIIIIIIIIIII6IBIIIIIIIIIIIIIIIIIIIIIIIIGII>IIIII-I)8I
```

##### Output

The output of the `gto_fastq_rand_extra_chars` program is a FASTQ file.
Using the input above, an output example for this is the following:

```
@SRR001666.1 071112_SLXA-EAS1_s_7:5:1:817:345 length=72
GTGTGATGGCCGCTGCCGATGGCGCATAATCCCACCAACATACCCTTAACAACTTAAGGGTTTTCAAATAGA
+SRR001666.1 071112_SLXA-EAS1_s_7:5:1:817:345 length=72
IIIIIIIIIIIIIIIIIIIIIIIIIIIIII9IG9ICIIIIIIIIIIIIIIIIIIIIDIIIIIII>IIIIII/
@SRR001666.2 071112_SLXA-EAS1_s_7:5:1:801:338 length=72
GTTCAGGGATACGACGATTGTATTTTAAGAATCTGCAGCAGAAGTCGATGATAATACGCGCCGTTTTATCAG
+SRR001666.2 071112_SLXA-EAS1_s_7:5:1:801:338 length=72
IIIIIIIIIIIIIIIIIIIIIIIIIIIIII6IBIIIIIIIIIIIIIIIIIIIIIIIGII>IIIII-I)8I
```

#### 2.10 Program gto_fastq_from_seq

The `gto_fastq_from_seq` converts a genomic sequence to pseudo FASTQ file format.
For help type:

```
./gto_fastq_from_seq -h
```

In the following subsections, we explain the input and output paramters.

##### Input parameters

The `gto_fastq_from_seq` program needs two streams for the computation, namely the input and output standard. The input stream is a sequence group file.
The attribution is given according to:

```
Usage: ./gto_fastq_from_seq [options] [[--] args]
   or: ./gto_fastq_from_seq [options]

It converts a genomic sequence to pseudo FASTQ file format.

    -h, --help            show this help message and exit

Basic options
    < input.seq           Input sequence file (stdin)
    > output.fastq        Output FASTQ file format (stdout)

Optional options
    -n, --name=<str>      The read's header
    -l, --lineSize=<int>  The maximum of chars for line

Example: ./gto_fastq_from_seq -l <lineSize> -n <name> < input.seq > output.fastq
```

An example of such an input file is:

```
ACAAGACGGCCTCCTGCTGCTGCTGCTCTCCGGGGCCACGGCCCTGGAGGGTCCACCGCTGCCCTGCTGCCATTGTCCCC
GGCCCCACCTAAGGAAAAGCAGCCTCCTGACTTTCCTCGCTTGGGCCGAGACAGCGAGCATATGCAGGAAGCGGCAGGAA
GTGGTTTGAGTGGACCTCCGGGCCCCTCATAGGAGAGGAAGCTCGGGAGGTGGCCAGGCGGCAGGAAGCAGGCCAGTGCC
GCGAATCCGCGCGCCGGGACAGAATCTCCTGCAAAGCCCTGCAGGAACTTCTTCTGGAAGACCTTCTCCACCCCCCCAGC
TAAAACCTCACCCATGAATGCTCACGCAAGTTTAATTACAGACCTGAAACAAGATGCCATTGTCCCCCGGCCTCCTGCTG
CTGCTGCTCTCCGGGGCCACGGCCACCGCTGCCCTGCCCCTGGAGGGTGGCCCCACCGGCCGAGACAGCGAGCATATGCA
GGAAGCGGCAGGAATAAGGAAAAGCAGCCTCCTGACTTTCCTCGCTTGGTGGTTTGAGTGGACCTCCCAGGCCAGTGCCG
GGCCCCTCATAGGAGAGGAAGCTCGGGAGGTGGCCAGGCGGCAGGAAGGCGCACCCCCCCAGCAATCCGCGCGCCGGGAC
AGAATGCCCTGCAGGAACTTCTTCTGGAAGACCTTCTCCTCCTGCAAATAAAACCTCACCCATGAATGCTCACGCAAGTT
TAATTACAGACCTGAA
```

#### Output

The output of the `gto_fastq_from_seq` program is a pseudo FASTQ file.
An example, using the size line as 80 and the read's header as "SeqToFastq", for the input, is:

```
@SeqToFastq1
ACAAGACGGCCTCCTGCTGCTGCTGCTCTCCGGGGCCACGGCCCTGGAGGGTCCACCGCTGCCCTGCTGCCATTGTCCCC
+
FFFFFFFFFFFFFFFFFFFFFFFFFFFFFFFFFFFFFFFFFFFFFFFFFFFFFFFFFFFFFFFFFFFFFFFFFFFFFFFF
@SeqToFastq2
GGCCCCACCTAAGGAAAAGCAGCCTCCTGACTTTCCTCGCTTGGGCCGAGACAGCGAGCATATGCAGGAAGCGGCAGGAA
+
FFFFFFFFFFFFFFFFFFFFFFFFFFFFFFFFFFFFFFFFFFFFFFFFFFFFFFFFFFFFFFFFFFFFFFFFFFFFFFFF
@SeqToFastq3
GTGGTTTGAGTGGACCTCCGGGCCCCTCATAGGAGAGGAAGCTCGGGAGGTGGCCAGGCGGCAGGAAGCAGGCCAGTGCC
+
FFFFFFFFFFFFFFFFFFFFFFFFFFFFFFFFFFFFFFFFFFFFFFFFFFFFFFFFFFFFFFFFFFFFFFFFFFFFFFFF
@SeqToFastq4
GCGAATCCGCGCGCCGGGACAGAATCTCCTGCAAAGCCCTGCAGGAACTTCTTCTGGAAGACCTTCTCCACCCCCCCAGC
+
FFFFFFFFFFFFFFFFFFFFFFFFFFFFFFFFFFFFFFFFFFFFFFFFFFFFFFFFFFFFFFFFFFFFFFFFFFFFFFFF
@SeqToFastq5
TAAAACCTCACCCATGAATGCTCACGCAAGTTTAATTACAGACCTGAAACAAGATGCCATTGTCCCCCGGCCTCCTGCTG
+
FFFFFFFFFFFFFFFFFFFFFFFFFFFFFFFFFFFFFFFFFFFFFFFFFFFFFFFFFFFFFFFFFFFFFFFFFFFFFFFF
@SeqToFastq6
CTGCTGCTCTCCGGGGCCACGGCCACCGCTGCCCTGCCCCTGGAGGGTGGCCCCACCGGCCGAGACAGCGAGCATATGCA
+
FFFFFFFFFFFFFFFFFFFFFFFFFFFFFFFFFFFFFFFFFFFFFFFFFFFFFFFFFFFFFFFFFFFFFFFFFFFFFFFF
@SeqToFastq7
GGAAGCGGCAGGAATAAGGAAAAGCAGCCTCCTGACTTTCCTCGCTTGGTGGTTTGAGTGGACCTCCCAGGCCAGTGCCG
+
FFFFFFFFFFFFFFFFFFFFFFFFFFFFFFFFFFFFFFFFFFFFFFFFFFFFFFFFFFFFFFFFFFFFFFFFFFFFFFFF
@SeqToFastq8
GGCCCCTCATAGGAGAGGAAGCTCGGGAGGTGGCCAGGCGGCAGGAAGGCGCACCCCCCCAGCAATCCGCGCGCCGGGAC
+
FFFFFFFFFFFFFFFFFFFFFFFFFFFFFFFFFFFFFFFFFFFFFFFFFFFFFFFFFFFFFFFFFFFFFFFFFFFFFFFF
@SeqToFastq9
AGAATGCCCTGCAGGAACTTCTTCTGGAAGACCTTCTCCTCCTGCAAATAAAACCTCACCCATGAATGCTCACGCAAGTT
+
FFFFFFFFFFFFFFFFFFFFFFFFFFFFFFFFFFFFFFFFFFFFFFFFFFFFFFFFFFFFFFFFFFFFFFFFFFFFFFFF
```

```
@SeqToFastq10
TAATTACAGACCTGAA
+
FFFFFFFFFFFFFFFF
```

#### 2.11 Program gto_fastq_mutate

The **gto_fastq_mutate** creates a synthetic mutation of a FASTQ file given specific rates of mutations, deletions and additions. All these paramenters are defined by the user, and their are optional.
For help type:

```
./gto_fastq_mutate -h
```

In the following subsections, we explain the input and output paramters.

##### Input parameters

The **gto_fastq_mutate** program needs two streams for the computation, namely the input and output standard. However, optional settings can be supplied too, such as the starting point to the random generator, and the edition, deletion and insertion rates. Also, the user can choose to use the ACGTN alphabet in the synthetic mutation. The input stream is a FASTQ File.
The attribution is given according to:

```
Usage: ./gto_fastq_mutate [options] [[--] args]
   or: ./gto_fastq_mutate [options]

Creates a synthetic mutation of a FASTQ file given specific rates of mutations,
deletions and additions

    -h, --help                  show this help message and exit

Basic options
    < input.fasta               Input FASTQ file format (stdin)
    > output.fasta              Output FASTQ file format (stdout)

Optional
    -s, --seed=<int>            Starting point to the random generator
    -m, --mutation-rate=<dbl>   Defines the mutation rate (default 0.0)
    -d, --deletion-rate=<dbl>   Defines the deletion rate (default 0.0)
    -i, --insertion-rate=<dbl>  Defines the insertion rate (default 0.0)
    -a, --ACGTN-alphabet        When active, the application uses the ACGTN alphabet

Example: ./gto_fastq_mutate -s <seed> -m <mutation rate> -d <deletion rate> -i
<insertion rate> -a < input.fastq > output.fastq
```

An example of such an input file is:

```
@SRR001666.1 071112_SLXA-EAS1_s_7:5:1:817:345 length=72
GGGTGATGGCCGCTGCCGATGGCGTCAAATCCCACCAAGTTACCCTTAACAACTTAAGGGTTTTCAAATAGA
+SRR001666.1 071112_SLXA-EAS1_s_7:5:1:817:345 length=72
IIIIIIIIIIIIIIIIIIIIIIIIIIIIII9IG9ICIIIIIIIIIIIIIIIIIIIIDIIIIIII>IIIIII/
@SRR001666.2 071112_SLXA-EAS1_s_7:5:1:801:338 length=72
GTTCAGGGATACGACGTTTGTATTTTAAGAATCTGAAGCAGAAGTCGATGATAATACGCGTCGTTTTATCAT
+SRR001666.2 071112_SLXA-EAS1_s_7:5:1:801:338 length=72
IIIIIIIIIIIIIIIIIIIIIIIIIIIIIIII6IBIIIIIIIIIIIIIIIIIIIIIIIGII>IIIII-I)8I
```

##### Output

The output of the `gto_fastq_mutate` program is a FASTQ file whith the synthetic mutation of input file. Using the input above with the seed value as 1 and the mutation rate as 0.5, an output example for this is the following:

```
@SRR001666.1 071112_SLXA-EAS1_s_7:5:1:817:345 length=72
GGACTTTGAGGTGTGGCGATAGACTGAAAACACTTCAGGGTAAAATCACTCGCAAAAGTGCTATGGTTATGG
+SRR001666.1 071112_SLXA-EAS1_s_7:5:1:817:345 length=72
IIIIIIIIIIIIIIIIIIIIIIIIIIIIII9IG9ICIIIIIIIIIIIIIIIIIIIIDIIIIIII>IIIIII/
@SRR001666.2 071112_SLXA-EAS1_s_7:5:1:801:338 length=72
GTTCAGAGCCTTTACCGTAGGGGTGTAAGATTTTATACAAAAAGTCCAGGTCAAGAGGAATCGGACAACCGA
+SRR001666.2 071112_SLXA-EAS1_s_7:5:1:801:338 length=72
IIIIIIIIIIIIIIIIIIIIIIIIIIIIIIII6IBIIIIIIIIIIIIIIIIIIIIIIIGII>IIIII-I)8I
```

#### 2.12 Program gto_fastq_split

The `gto_fastq_split` splits Paired End files according to the direction of the strand ('/1' or '/2'). It writes by default singleton reads as forward stands.
For help type:

```
./gto_fastq_split -h
```

In the following subsections, we explain the input and output paramters.

##### Input parameters

The `gto_fastq_split` program needs a stream for the computation, namely the input standard. The input stream is a FASTQ file.
The attribution is given according to:

```
Usage: ./gto_fastq_split [options] [[--] args]
   or: ./gto_fastq_split [options]

It writes by default singleton reads as forward stands.

    -h, --help            show this help message and exit
```

```
Basic options
    -f, --forward=<str>   Output forward file
    -r, --reverse=<str>   Output reverse file
    < input.fastq         Input FASTQ file format (stdin)
    > output              Output read information (stdout)

Example: ./gto_fastq_split -f <output_forward.fastq> -r <output_reverse.fastq> < input.fastq > output

Output example :
Total reads      : value
Singleton reads  : value
Forward reads    : value
Reverse reads    : value
```

An example of such an input file is:

```
@SRR001666.1 071112_SLXA-EAS1_s_7:5:1:817:345 length=72 1
GNNTGATGGCCGCTGCCGATGGCGNANAATCCCACCAANATACCCTTAACAACTTAAGGGTTNTCAAATAGA
+
IIIIIIIIIIIIIIIIIIIIIIIIIIIIII9IG9ICIIIIIIIIIIIIIIIIIIIIDIIIIIII>IIIIII/
@SRR001666.2 071112_SLXA-EAS1_s_7:5:1:801:338 length=72 2
NTTCAGGGATACGACGNTTGTATTTTAAGAATCTGNAGCAGAAGTCGATGATAATACGCGNCGTTTTATCAN
+
IIIIIIIIIIIIIIIIIIIIIIIIIIIIIIII6IBIIIIIIIIIIIIIIIIIIIIIIIGII>IIIII-I)8I
```

##### Output

The output of the `gto_fastq_split` program is a set of information related to the file read.
Using the input above, an output example for this is the following:

```
Total reads     : 2
Singleton reads : 0
Forward reads   : 65536
Reverse reads   : 1
```

Also, this program generates two FASTQ files, with the reverse and forward reads.
An example of the forward reads, for the input, is:

```
@SRR001666.1 071112_SLXA-EAS1_s_7:5:1:817:345 length=72 1
GNNTGATGGCCGCTGCCGATGGCGNANAATCCCACCAANATACCCTTAACAACTTAAGGGTTNTCAAATAGA
+
IIIIIIIIIIIIIIIIIIIIIIIIIIIIII9IG9ICIIIIIIIIIIIIIIIIIIIIDIIIIIII>IIIIII/
```

#### 2.13 Program gto_fastq_pack

The `gto_fastq_pack` packages each FASTQ read in a single line. It can show the read score first or the dna sequence, depending on the execution mode.
For help type:

```
./gto_fastq_pack -h
```

In the following subsections, we explain the input and output paramters.

##### Input parameters

The `gto_fastq_pack` program needs two streams for the computation, namely the input and output standard. The input stream is a FASTQ file.
The attribution is given according to:

```
Usage: ./gto_fastq_pack [options] [[--] args]
   or: ./gto_fastq_pack [options]

It packages each FASTQ read in a single line.

    -h, --help             show this help message and exit

Basic options
    < input.fastq          Input FASTQ file format (stdin)
    > output.fastqpack     Output packaged FASTQ file format (stdout)

Optional
    -s, --scores           When active, the application show the scores first

Example: ./gto_fastq_pack -s < input.fastq > output.fastqpack
```

An example of such an input file is:

```
@SRR001666.1 071112_SLXA-EAS1_s_7:5:1:817:345 length=72
GNNTGATGGCCGCTGCCGATGGCGNANAATCCCACCAANATACCCTTAACAACTTAAGGGTTNTCAAATAGA
+
IIIIIIIIIIIIIIIIIIIIIIIIIIII9IG9ICIIIIIIIIIIIIIIIIIIIIDIIIIIII>IIIIII/
@SRR001666.2 071112_SLXA-EAS1_s_7:5:1:801:338 length=72
NTTCAGGGATACGACGNTTGTATTTTAAGAATCTGNAGCAGAAGTCGATGATAATACGCGNCGTTTTATCAN
+
IIIIIIIIIIIIIIIIIIIIIIIIIIIIIII6IBIIIIIIIIIIIIIIIIIIIIIIIGII>IIIII-I)8I
```

##### Output

The output of the `gto_fastq_pack` program is a packaged FASTQ file.
Using the input above, an output example for this is the following:

```
GNNTGATGGCCGCTGCCGATGGCGNANAATCCCACCAANATACCCTTAACAACTTAAGGGTTNTCAAATAGA
IIIIIIIIIIIIIIIIIIIIIIIIIIII9IG9ICIIIIIIIIIIIIIIIIIIIIDIIIIIII>IIIIII/
SRR001666.1 071112_SLXA-EAS1_s_7:5:1:817:345 length=72+ 0
NTTCAGGGATACGACGNTTGTATTTTAAGAATCTGNAGCAGAAGTCGATGATAATACGCGNCGTTTTATCAN
IIIIIIIIIIIIIIIIIIIIIIIIIIIIIII6IBIIIIIIIIIIIIIIIIIIIIIIIGII>IIIII-I)8I
SRR001666.2 071112_SLXA-EAS1_s_7:5:1:801:338 length=72+ 1
```

Another example for the same input, but using the scores first (flag ”s”), is:

```
IIIIIIIIIIIIIIIIIIIIIIIIIIIIIIII9IG9ICIIIIIIIIIIIIIIIIIIIIIDIIIIIII>IIIIII/
GNNTGATGGCCGCTGCCGATGGCGNANAATCCCACCAANATACCCTTAACAACTTAAGGGTTNTCAAATAGA
SRR001666.1 071112_SLXA-EAS1_s_7:5:1:817:345 length=72+ 0
IIIIIIIIIIIIIIIIIIIIIIIIIIIIIIII6IBIIIIIIIIIIIIIIIIIIIIIIIGII>IIIII-I)8I
NTTCAGGGATACGACGNTTGTATTTTAAGAATCTGNAGCAGAAGTCGATGATAATACGCGNCGTTTTATCAN
SRR001666.2 071112_SLXA-EAS1_s_7:5:1:801:338 length=72+ 1
```

#### 2.14 Program gto_fastq_unpack

The `gto_fastq_unpack` unpacks the FASTQ reads packaged using the `gto_fastq_pack` tool.
For help type:

```
./gto_fastq_unpack -h
```

In the following subsections, we explain the input and output paramters.

##### Input parameters

The `gto_fastq_unpack` program needs two streams for the computation, namely the input and output standard. The input stream is a packaged FASTQ file.
The attribution is given according to:

```
Usage: ./gto_fastq_unpack [options] [[--] args]
   or: ./gto_fastq_unpack [options]

It unpacks the FASTQ reads packaged using the gto_fastq_pack tool.

    -h, --help             show this help message and exit

Basic options
    < input.fastq          Input FASTQ file format (stdin)
    > output.fastq         Output FASTQ file format (stdout)

Optional
    -s, --scores           When active, the application show the scores first

Example: ./gto_fastq_unpack -s < input.fastqpack > output.fastq
```

An example of such an input file is:

```
GNNTGATGGCCGCTGCCGATGGCGNANAATCCCACCAANATACCCTTAACAACTTAAGGGTTNTCAAATAGA
IIIIIIIIIIIIIIIIIIIIIIIIIIIIIIII9IG9ICIIIIIIIIIIIIIIIIIIIIIDIIIIIII>IIIIII/
SRR001666.1 071112_SLXA-EAS1_s_7:5:1:817:345 length=72+ 0
NTTCAGGGATACGACGNTTGTATTTTAAGAATCTGNAGCAGAAGTCGATGATAATACGCGNCGTTTTATCAN
IIIIIIIIIIIIIIIIIIIIIIIIIIIIIIII6IBIIIIIIIIIIIIIIIIIIIIIIIGII>IIIII-I)8I
SRR001666.2 071112_SLXA-EAS1_s_7:5:1:801:338 length=72+ 1
```

##### Output

The output of the `gto_fastq_unpack` program is a FASTQ file.
Using the input above, an output example for this is the following:

```
@SRR001666.1 071112_SLXA-EAS1_s_7:5:1:817:345 length=72
GNNTGATGGCCGCTGCCGATGGCGNANAATCCCACCAANATACCCTTAACAACTTAAGGGTTNTCAAATAGA
+
IIIIIIIIIIIIIIIIIIIIIIIIIIIIII9IG9ICIIIIIIIIIIIIIIIIIIIIDIIIIIII>IIIIII/
@SRR001666.2 071112_SLXA-EAS1_s_7:5:1:801:338 length=72
NTTCAGGGATACGACGNTTGTATTTTAAGAATCTGNAGCAGAAGTCGATGATAATACGCGNCGTTTTATCAN
+
IIIIIIIIIIIIIIIIIIIIIIIIIIIIIIII6IBIIIIIIIIIIIIIIIIIIIIIIIGII>IIIII-I)8I
```

#### 2.15 Program gto_fastq_quality_score_info

The `gto_fastq_quality_score_info` analyses the quality-scores of a FASTQ file.
For help type:

```
./gto_fastq_quality_score_info -h
```

In the following subsections, we explain the input and output paramters.

##### Input parameters

The `gto_fastq_quality_score_info` program needs two streams for the computation, namely the input and output standard. The input stream is a FASTQ file.
The attribution is given according to:

```
Usage: ./gto_fastq_quality_score_info [options] [[--] args]
   or: ./gto_fastq_quality_score_info [options]

It analyses the quality-scores of a FASTQ file.

    -h, --help            show this help message and exit

Basic options
    < input.fastq         Input FASTQ file format (stdin)
    > output              Output read information (stdout)

Optional
    -m, --max=<int>       The lenght of the maximum window

Example: ./gto_fastq_quality_score_info -m <max> < input.fastq > output

Output example :
Total reads     : value
Max read length : value
Min read length : value
Min QS value    : value
```

```
Max QS value    : value
QS range        : value
```

An example of such an input file is:

```
@111 071112_SLXA-EAS1_s_7:5:1:817:345 length=72 1
GNNTGATGGCCGCTGCCGATGGCGNANAATCCCACCAANATACCCTTAACAACTTAAGGGTTNTCAAATAGA
+111
IIIIIIIIIIIIIIIIIIIIIIIIIIIIII9IG9ICIIIIIIIIIIIIIIIIIIIIDIIIIIII>IIIIII/
@222 071112_SLXA-EAS1_s_7:5:1:801:338 length=72 2
NTTCAGGGATACGACGNTTGTATTTTAAGAATCTGNAGCAGAAGTCGATGATAATACGCGNCGTTTTATCAN
+222
IIIIIIIIIIIIIIIIIIIIIIIIIIIIIIII6IBIIIIIIIIIIIIIIIIIIIIIIIIIGII>IIIII-I)8I
```

##### Output

The output of the `gto_fastq_quality_score_info` program is a set of information related to the file read. Using the input above with the max window value as 30, an output example for this is the following:

```
Total reads     : 2
Max read length : 72
Min read length : 72
Min QS value    : 41
Max QS value    : 73
QS range        : 33
 1  2  3  4  5  6  7  8  9 10 11 12 13 14 15 16 17 18 19 20 21 22 23 24 25 26 27 28 29 30
--+--+--+--+--+--+--+--+--+--+--+--+--+--+--+--+--+--+--+--+--+--+--+--+--+--+--+--+--+--+
73 73 73 73 73 73 73 73 73 73 73 73 73 73 73 73 73 73 73 73 73 73 73 73 73 73 73 73 73 73
```

#### 2.16 Program gto_fastq_quality_score_max

The `gto_fastq_quality_score_max` analyses the maximal quality-scores of a FASTQ file.
For help type:

```
./gto_fastq_quality_score_max -h
```

In the following subsections, we explain the input and output paramters.

##### Input parameters

The `gto_fastq_quality_score_max` program needs two streams for the computation, namely the input and output standard. The input stream is a FASTQ file.
The attribution is given according to:

```
Usage: ./gto_fastq_quality_score_max [options] [[--] args]
   or: ./gto_fastq_quality_score_max [options]

It analyses the maximal quality-scores of a FASTQ file.
```

```
    -h, --help             show this help message and exit

Basic options
    < input.fastq          Input FASTQ file format (stdin)
    > output               Output read information (stdout)

Optional
    -m, --max=<int>        The maximum window length (default 40)

Example: ./gto_fastq_quality_score_max -m <max> < input.fastq > output
```

An example of such an input file is:

```
@111 071112_SLXA-EAS1_s_7:5:1:817:345 length=72 1
GNNTGATGGCCGCTGCCGATGGCGNANAATCCCACCAANATACCCTTAACAACTTAAGGGTTNTCAAATAGA
+
IIIIIIIIIII9IG9ICIIIIIIIIIIIABAAABCIIIIFFGIIAACBBIIIII6IBIIIIIII>IIIIII/
@222 071112_SLXA-EAS1_s_7:5:1:801:338 length=72 2
NTTCAGGGATACGACGNTTGTATTTTAAGAATCTGNAGCAGAAGTCGATGATAATACGCGNCGTTTTATCAN
+
IIIIIIIABAAABCIIIIFFGIIAACBBIIIII6IBIIIIIIIIIIIIIIIIIIIIIIIGII>IIIII-I)8I
```

##### Output

The output of the `gto_fastq_quality_score_max` program is a set of information related to the file read, considering the maximal quality scores.
Using the input above with the max window value as 30, an output example for this is the following:

```
 1  2  3  4  5  6  7  8  9 10 11 12 13 14 15 16 17 18 19 20 21 22 23 24 25 26 27 28 29 30
--+--+--+--+--+--+--+--+--+--+--+--+--+--+--+--+--+--+--+--+--+--+--+--+--+--+--+--+--+--+
73 73 73 73 73 73 73 73 73 73 73 73 66 73 73 73 73 73 73 73 73 73 73 73 73 73 73 73 73 73
```

#### 2.17 Program gto_fastq_quality_score_min

The `gto_fastq_quality_score_min` analyses the minimal quality-scores of a FASTQ file.
For help type:

```
./gto_fastq_quality_score_min -h
```

In the following subsections, we explain the input and output paramters.

##### Input parameters

The `gto_fastq_quality_score_min` program needs two streams for the computation, namely the input and output standard. The input stream is a FASTQ file.
The attribution is given according to:

```
Usage: ./gto_fastq_quality_score_min [options] [[--] args]
   or: ./gto_fastq_quality_score_min [options]

It analyses the minimal quality-scores of a FASTQ file.

    -h, --help            show this help message and exit

Basic options
    < input.fastq         Input FASTQ file format (stdin)
    > output              Output read information (stdout)

Optional
    -m, --max=<int>       The maximum window length (default 40)

Example: ./gto_fastq_quality_score_min -m <max> < input.fastq > output
```

An example of such an input file is:

```
@111 071112_SLXA-EAS1_s_7:5:1:817:345 length=72 1
GNNTGATGGCCGCTGCCGATGGCGNANAATCCCACCAANATACCCTTAACAACTTAAGGGTTNTCAAATAGA
+
IIIIIIIIIII9IG9ICIIIIIIIIIIIABAAABCIIIIFFGIIAACBBIIIII6IBIIIIIII>IIIIII/
@222 071112_SLXA-EAS1_s_7:5:1:801:338 length=72 2
NTTCAGGGATACGACGNTTGTATTTTAAGAATCTGNAGCAGAAGTCGATGATAATACGCGNCGTTTTATCAN
+
IIIIIIIABAAABCIIIIFFGIIAACBBIIIII6IBIIIIIIIIIIIIIIIIIIIIIIIGII>IIIII-I)8I
```

##### Output

The output of the `gto_fastq_quality_score_min` program is a set of information related to the file read, considering the minimum quality scores.
Using the input above with the max window value as 30, an output example for this is the following:

```
 1  2  3  4  5  6  7  8  9 10 11 12 13 14 15 16 17 18 19 20 21 22 23 24 25 26 27 28 29 30
--+--+--+--+--+--+--+--+--+--+--+--+--+--+--+--+--+--+--+--+--+--+--+--+--+--+--+--+--+--+
73 73 73 73 73 73 73 65 66 65 65 65 57 67 71 57 73 67 70 70 71 73 73 65 65 67 66 66 73 65
```

#### 2.18 Program gto_fastq_cut

The `gto_fastq_cut` cuts read sequences in a FASTQ file. It requires that the initial and end positions for the cut.
For help type:

```
./gto_fastq_cut -h
```

In the following subsections, we explain the input and output paramters.

##### Input parameters

The `gto_fastq_cut` program needs two streams for the computation, namely the input and output standard. The input stream is a FASTQ file.
The attribution is given according to:

```
Usage: ./gto_fastq_cut [options] [[--] args]
   or: ./gto_fastq_cut [options]

It cuts read sequences in a FASTQ file.

    -h, --help            show this help message and exit

Basic options
    -i, --initial=<int>   Starting position to the cut
    -e, --end=<int>       Ending position to the cut
    < input.fastq         Input FASTQ file format (stdin)
    > output.fastq        Output FASTQ file format (stdout)

Example: ./gto_fastq_cut -i <initial> -e <end> < input.fastq > output.fastq
```

An example of such an input file is:

```
@SRR001666.1 071112_SLXA-EAS1_s_7:5:1:817:345 length=72
GGGTGATGGCCGCTGCCGATGGCGTCAAATCCCACCAAGTTACCCTTAACAACTTAAGGGTTTTCAAATAGA
+SRR001666.1 071112_SLXA-EAS1_s_7:5:1:817:345 length=72
IIIIIIIIIIIIIIIIIIIIIIIIIIIIII9IG9ICIIIIIIIIIIIIIIIIIIIIDIIIIIII>IIIIII/
@SRR001666.2 071112_SLXA-EAS1_s_7:5:1:801:338 length=72
GTTCAGGGATACGACGTTTGTATTTTAAGAATCTGAAGCAGAAGTCGATGATAATACGCGTCGTTTTATCAT
+SRR001666.2 071112_SLXA-EAS1_s_7:5:1:801:338 length=72
IIIIIIIIIIIIIIIIIIIIIIIIIIIIIIII6IBIIIIIIIIIIIIIIIIIIIIIIIGII>IIIII-I)8I
```

##### Output

The output of the `gto_fastq_cut` program is a FASTQ file cut.
Using the initial value as 10 and the end value as 30, an example for this input, is:

```
@SRR001666.1 071112_SLXA-EAS1_s_7:5:1:817:345 length=72
CGCTGCCGATGGCGTCAAATC
+
IIIIIIIIIIIIIIIIIIII9
@SRR001666.2 071112_SLXA-EAS1_s_7:5:1:801:338 length=72
ACGACGTTTGTATTTTAAGAA
+
IIIIIIIIIIIIIIIIIIIII
```

#### 2.19 Program gto_fastq_minimum_local_quality_score_forward

The `gto_fastq_minimum_local_quality_score_forward` filters the reads considering the quality score average of a defined window size of bases.

For help type:

```
./gto_fastq_minimum_local_quality_score_forward -h
```

In the following subsections, we explain the input and output paramters.

##### Input parameters

The `gto_fastq_minimum_local_quality_score_forward` program needs two streams for the computation, namely the input and output standard. The input stream is a FASTQ file.
The attribution is given according to:

```
Usage: ./gto_fastq_minimum_local_quality_score_forward [options] [[--] args]
   or: ./gto_fastq_minimum_local_quality_score_forward [options]

It filters the reads considering the quality score average of a defined window size
of bases.

    -h, --help                show this help message and exit

Basic options
    -k, --windowsize=<int>    The window size of bases (default 5)
    -w, --minavg=<int>        The minimum average of quality score (default 25)
    -m, --minqs=<int>         The minimum value of the quality score (default 33)
    < input.fastq             Input FASTQ file format (stdin)
    > output.fastq            Output FASTQ file format (stdout)

Example: ./gto_fastq_minimum_local_quality_score_forward -k <windowsize> -w <minavg>
-m <minqs> < input.fastq > output.fastq

Console output example:
Minimum QS       : value
<FASTQ output>
Total reads      : value
Trimmed reads    : value
```

An example of such an input file is:

```
@SRR001666.1 071112_SLXA-EAS1_s_7:5:1:817:345 length=72
GGGTGATGGCCGCTGCCGATGGCGTCAAATCCCACCAAGTTACCCTTAACAACTTAAGGGTTTTCAAATAGA
+SRR001666.1 071112_SLXA-EAS1_s_7:5:1:817:345 length=72
IIIIIIIIIIIIIIIIIIIIIIIIIIIIII9IG9ICIIIIIIIIIIIIIIIIIIIIDIIIIIII>IIIIII/
@SRR001666.2 071112_SLXA-EAS1_s_7:5:1:801:338 length=72
GTTCAGGGATACGACGTTTGTATTTTAAGAATCTGAAGCAGAAGTCGATGATAATACGCGTCGTTTTATCAT
+SRR001666.2 071112_SLXA-EAS1_s_7:5:1:801:338 length=72
IIIIIIIIIIIIIIIIIIIIIIIIIIIIIIII6IBIIIIIIIIIIIIIIIIIIIIIIIGII>IIIII-I)8I
```

##### Output

The output of the `gto_fastq_minimum_local_quality_score_forward` program is a FASTQ file with the reads filtered following a quality score average of a defined window of bases. The execution report only appears in the console.

Using the input above with the default values, an output example for this is the following:

```
Minimum QS      : 33
@SRR001666.1 071112_SLXA-EAS1_s_7:5:1:817:345 length=72
GGGTGATGGCCGCTGCCGATGGCGTCAAATCCCACCAAGTTACCCTTAACAACTTAAGGGTTTTCAAATAGA
+
IIIIIIIIIIIIIIIIIIIIIIIIIIIIII9IG9ICIIIIIIIIIIIIIIIIIIIIDIIIIIII>IIIIII/
@SRR001666.2 071112_SLXA-EAS1_s_7:5:1:801:338 length=72
GTTCAGGGATACGACGTTTGTATTTTAAGAATCTGAAGCAGAAGTCGATGATAATACGCGTCGTTT
+
IIIIIIIIIIIIIIIIIIIIIIIIIIIIIII6IBIIIIIIIIIIIIIIIIIIIIIIIGII>IIII
Total reads     : 2
Trimmed reads   : 1
```

#### 2.20 Program gto_fastq_minimum_local_quality_score_reverse

The `gto_fastq_minimum_local_quality_score_reverse` filters the reverse reads, considering the quality score average of a defined window size of bases.

For help type:

```
./gto_fastq_minimum_local_quality_score_reverse -h
```

In the following subsections, we explain the input and output paramters.

##### Input parameters

The `gto_fastq_minimum_local_quality_score_reverse` program needs two streams for the computation, namely the input and output standard. The input stream is a FASTQ file.

The attribution is given according to:

```
Usage: ./gto_fastq_minimum_local_quality_score_reverse [options] [[--] args]
   or: ./gto_fastq_minimum_local_quality_score_reverse [options]

It filters the reverse reads, considering the quality score average of a defined
window size of bases.

    -h, --help                show this help message and exit

Basic options
    -k, --windowsize=<int>    The window size of bases (default 5)
    -w, --minavg=<int>        The minimum average of quality score (default 25)
    -m, --minqs=<int>         The minimum value of the quality score (default 33)
    < input.fastq             Input FASTQ file format (stdin)
    > output.fastq            Output FASTQ file format (stdout)

Example: ./gto_fastq_minimum_local_quality_score_reverse -k <windowsize> -w <minavg>
-m <minqs> < input.fastq > output.fastq
```

```
Console output example:
Minimum QS       : value
<FASTQ output>
Total reads      : value
Trimmed reads    : value
```

An example of such an input file is:

```
@SRR001666.1 071112_SLXA-EAS1_s_7:5:1:817:345 length=72
GGGTGATGGCCGCTGCCGATGGCGTCAAATCCCACCAAGTTACCCTTAACAACTTAAGGGTTTTCAAATAGA
+SRR001666.1 071112_SLXA-EAS1_s_7:5:1:817:345 length=72
IIIIIIIIIIIIIIIIIIIIIIIIIIIII9IG9ICIIIIIIIIIIIIIIIIIIIIDIIIIIII>IIIIII/
@SRR001666.2 071112_SLXA-EAS1_s_7:5:1:801:338 length=72
GTTCAGGGATACGACGTTTGTATTTTAAGAATCTGAAGCAGAAGTCGATGATAATACGCGTCGTTTTATCAT
+SRR001666.2 071112_SLXA-EAS1_s_7:5:1:801:338 length=72
IIIIIIIIIIIIIIIIIIIIIIIIIIIIIIII6IBIIIIIIIIIIIIIIIIIIIIIIIGII>IIIII-I)8I
```

##### Output

The output of the `gto_fastq_minimum_local_quality_score_reverse` program is a FASTQ file with the reads filtered following a quality score average of a defined window of bases. The execution report only appears in the console.
Using the input above with the default values, an output example for this is the following:

```
Minimum QS: 33
@SRR001666.1 071112_SLXA-EAS1_s_7:5:1:817:345 length=72
GGGTGATGGCCGCTGCCGATGGCGTCAAATCCCACCAAGTTACCCTTAACAACTTAAGGGTTTTCAAATAGA
+
IIIIIIIIIIIIIIIIIIIIIIIIIIIII9IG9ICIIIIIIIIIIIIIIIIIIIIDIIIIIII>IIIIII/
Total reads    : 2
Trimmed reads  : 1
```

#### 2.21 Program gto_fastq_xs

The `gto_fastq_xs` is a skilled FASTQ read simulation tool, flexible, portable (does not need a reference sequence) and tunable in terms of sequence complexity. XS handles Ion Torrent, Roche-454, Illumina and ABI-SOLiD simulation sequencing types. It has several running modes, depending on the time and memory available, and is aimed at testing computing infrastructures, namely cloud computing of large-scale projects, and testing FASTQ compression algorithms. Moreover, XS offers the possibility of simulating the three main FASTQ components individually (headers, DNA sequences and quality-scores). Quality-scores can be simulated using uniform and Gaussian distributions.
For help type:

```
./gto_fastq_xs -h
```

In the following subsections, we explain the input and output paramters.

##### Input parameters

The `gto_fastq_xs` program needs a FASTQ file to compute.

The attribution is given according to:

```
Usage: ./gto_fastq_xs   [OPTION]... [FILE]

System options:
 -h                        give this help
 -v                        verbose mode

Main FASTQ options:
 -t  <sequencingType>      type: 1=Roche-454, 2=Illumina, 3=ABI SOLiD, 4=Ion Torrent
 -hf <headerFormat>        header format: 1=Length appendix, 2=Pair End
 -i  n=<instrumentName>    the unique instrument name (use n= before name)
 -o                        use the same header in third line of the read
 -ls <lineSize>            static line (bases/quality scores) size
 -ld <minSize>:<maxSize>   dynamic line (bases/quality scores) size
 -n  <numberOfReads>       number of reads per file

DNA options:
 -f  <A>,<C>,<G>,<T>,<N>   symbols frequency
 -rn <numberOfRepeats>     repeats: number (default: 0)
 -ri <repeatsMinSize>      repeats: minimum size
 -ra <repeatsMaxSize>      repeats: maximum size
 -rm <mutationRate>        repeats: mutation frequency
 -rr                       repeats: use reverse complement repeats

Quality scores options:
 -qt <assignmentType>      quality scores distribution: 1=uniform, 2=gaussian
 -qf <statsFile>           load file: mean, standard deviation (when: -qt 2)
 -qc <template>            custom template ascii alphabet

Filtering options:
 -eh                       excludes the use of headers from output
 -eo                       excludes the use of optional headers (+) from output
 -ed                       excludes the use of DNA bases from output
 -edb                      excludes '\n' when DNA bases line size is reached
 -es                       excludes the use of quality scores from output

Stochastic options:
 -s  <seed>                generation seed

<genFile>                  simulated output file

Common usage:
 ./XS -v -t 1 -i n=MySeq -ld 30:80 -n 20000 -qt=1 -qc 33,36,39:43 File
 ./XS -v -ls 100 -n 10000 -eh -eo -es -edb -f 0.3,0.2,0.2,0.3,0.0 -rn 50 -ri 300 -ra 3000 -rm 0.1 File
```

An example of such an input file is:

```
@SRR001666.1 071112_SLXA-EAS1_s_7:5:1:817:345 length=60
GGGTGATGGCCGCTGCCGATGGCGTCAAATCCCACCAAGTTACCCTTAACAACTTAAGGG
+
```

```
IIIIIIIIIIIIIIIIIIIIIIIIIIIIII9IG9ICIIIIIIIIIIIIIIIIIIIIDIII
@SRR001666.2 071112_SLXA-EAS1_s_7:5:1:801:338 length=72
GTTCAGGGATACGACGTTTGTATTTTAAGAATCTGAAGCAGAAGTCGATGATAATACGCGTCGTTTTATCAT
+
IIIIIIIIIIIIIIIIIIIIIIIIIIIIIIII6IBIIIIIIIIIIIIIIIIIIIIIIIGII>IIIII-I)8I
```

##### Output

The output of the `gto_fastq_xs` program is a FASTQ file
Using the input above using the common usage with 5 reads (-n 5), an output example for this is the following:

```
@output.fastq.598 LQGQLWH01D5WVZ length=62
TTCNTNCCAGGTAAAGAGAACATNCCGNCGCACTACTCGTAAGACTTGCTGGNCGAGAAAGG
+
)(+!*!$')($(()+'))$$()'!)!$!!$*+)+''('!)))!+!)(!+!*$!'$*)**++!
@output.fastq.1510 LQGQLWH01A7LJI length=57
CTAGACTACTCGAGCACTAGGCTCGCGTNTACCANGGGGNCTGCGNGTTGGCNCGGT
+
)+(*(+$*)+!*)!'!!(!(!!(*'$!+!(()$'!!+*+!!))!*!')***+!$+''
@output.fastq.2153 LQGQLWH01CHBQJ length=33
ACTTTTTGCTCAAGCAGGGTTGCCTAGCAANAC
+
*)++!+$''')*)**!+)$(*((*)$!'!+!!*
@output.fastq.3251 LQGQLWH01C8OY4 length=75
TCTTTCCTTCNCGNCCNAATTCCCCATAANAACTTAAAATCNCNNGCTGCGCGTGATCAACAATATTAATACTCC
+
!*''+*'!''!+!!!*!'!+(++)*(*($!!*((')$*!$(!'!!'+)$+*!$*!**!'()$!*'+'*'+!!+'(
@output.fastq.3934 LQGQLWH01AQDXM length=36
GGTAACNNGGAATTCTTCCAATTANCCNTGTCCGGC
+
$+)'!'!!)+)+!''**$$*!!')!+)!)*()!))$
```

#### 2.22 Program gto_fastq_clust_reads

The `gto_fastq_clust_reads` agroups reads and creates an index file. It cluster reads in therms of Seq k-mer Lexicographical order.
For help type:

```
./gto_fastq_clust_reads -h
```

In the following subsections, we explain the input and output paramters.

##### Input parameters

The `gto_fastq_clust_reads` program needs two streams for the computation, namely the input and output standard. The input stream is a FASTQ file. The program sorts the FASTQ reads accoring to the lexicographic order of the genomic sequences.
The attribution is given according to:

```
Usage: ./gto_fastq_clust_reads [options] [[--] args]
   or: ./gto_fastq_clust_reads [options]

It agroups reads and creates an index file.
It cluster reads in therms of Seq k-mer Lexicographical order

    -h, --help            Show this help message and exit

Basic options
    -c, --ctx=<int>
    < input.fastq         Input FASTQ file format (stdin)
    > output.fastq        Output FASTQ file format (stdout)

Example: ./gto_fastq_clust_reads -c <ctx> < input.fastq > output.fastq
```

An example of such an input file is:

```
@SRR001661.1 071112_SLXA-EAS1_s_7:5:1:817:345
GGGTGATGGCCGCTGCCGATGGCGTCAAATCCCACCAAGTTACCCTTAACAACTTAAGGG
+
IIIIIIIIIIIIIIIIIIIIIIIIIIIIII9IG9ICIIIIIIIIIIIIIIIIIIIIDIII
@SRR001661.2 071112_SLXA-EAS1_s_7:5:1:801:338
GTTCAGGGATACGACGTTTGTATTTTAAGAATCTGAAGCAGAAGTCGATGATAATACGCG
+
IIIIIIIIIIIIIIIIIIIIIIIIIIIIIIII6IBIIIIIIIIIIIIIIIIIIIIIIIGI
@SRR001661.3 071112_SLXA-EAS1_s_7:5:1:821:328
AACGCGTATTCGGAGCTTCTTCGTTGGGTACGTGCGCCTATTATGCGGCGCGATTGCTAT
+
IIIIII6BBB6BBBBBBBBBBBBBBBBBBDDDDDDDDDDDDDDDDDDDDDDDDDDDDDDD
@SRR001661.4 071112_SLXA-EAS1_s_7:5:1:943:128
ATCGCGCATTCGACTGGTACGTGTACGTGTAGTCGTAGCGTATGTTCGGTCGTATGCGTG
+
II77777LPMMMPPMMMMIIIIIIIIIIIII777777777BBBBBBBBDDDDDIIIIII
```

#### Output

The output of the `gto_fastq_clust_reads` program is a FASTQ file with clustered reads in therms of the genomic sequence k-mer Lexicographical order. An example, for the output, is:

```
@SRR001661.3 071112_SLXA-EAS1_s_7:5:1:821:328
AACGCGTATTCGGAGCTTCTTCGTTGGGTACGTGCGCCTATTATGCGGCGCGATTGCTAT
+
IIIIII6BBB6BBBBBBBBBBBBBBBBBBDDDDDDDDDDDDDDDDDDDDDDDDDDDDDDD
@SRR001661.4 071112_SLXA-EAS1_s_7:5:1:943:128
ATCGCGCATTCGACTGGTACGTGTACGTGTAGTCGTAGCGTATGTTCGGTCGTATGCGTG
+
II77777LPMMMPPMMMMIIIIIIIIIIIII777777777BBBBBBBBDDDDDIIIIII
@SRR001661.1 071112_SLXA-EAS1_s_7:5:1:817:345
GGGTGATGGCCGCTGCCGATGGCGTCAAATCCCACCAAGTTACCCTTAACAACTTAAGGG
+
IIIIIIIIIIIIIIIIIIIIIIIIIIIIII9IG9ICIIIIIIIIIIIIIIIIIIIIDIII
```

```
@SRR001661.2 071112_SLXA-EAS1_s_7:5:1:801:338
GTTCAGGGATACGACGTTTGTATTTTAAGAATCTGAAGCAGAAGTCGATGATAATACGCG
+
IIIIIIIIIIIIIIIIIIIIIIIIIIIIII6IBIIIIIIIIIIIIIIIIIIIIIIIIGI
```

#### 2.23 Program gto_fastq_complement

The `gto_fastq_complement` replaces the ACGT bases with their complements in a FASTQ file format. For help type:

```
./gto_fastq_complement -h
```

In the following subsections, we explain the input and output paramters.

##### Input parameters

The `gto_fastq_complement` program needs two streams for the computation, namely the input and output standard. The input stream is a FASTQ file.
The attribution is given according to:

```
Usage: ./gto_fastq_complement [options] [[--] args]
   or: ./gto_fastq_complement [options]

It replaces the ACGT bases with their complements in a FASTQ file format.

    -h, --help            Show this help message and exit

Basic options
    < input.fastq         Input FASTQ file (stdin)
    > output.fastq        Output FASTQ file (stdout)

Example: ./gto_fastq_complement < input.fastq > output.fastq
```

An example of such an input file is:

```
@SRR001666.1 071112_SLXA-EAS1_s_7:5:1:817:345 length=72
GGGTGATGGCCGCTGCCGATGGCGTCAAATCCCACCAAGTTACCCTTAACAACTTAAGGGTTTTCAAATAGA
+SRR001666.1 071112_SLXA-EAS1_s_7:5:1:817:345 length=72
IIIIIIIIIIIIIIIIIIIIIIIIIIIIII9IG9ICIIIIIIIIIIIIIIIIIIIIDIIIIIII>IIIIII/
@SRR001666.2 071112_SLXA-EAS1_s_7:5:1:801:338 length=72
GTTCAGGGATACGACGTTTGTATTTTAAGAATCTGAAGCAGAAGTCGATGATAATACGCGTCGTTTTATCAT
+SRR001666.2 071112_SLXA-EAS1_s_7:5:1:801:338 length=72
IIIIIIIIIIIIIIIIIIIIIIIIIIIIIIII6IBIIIIIIIIIIIIIIIIIIIIIIIIGII>IIIII-I)8I
```

##### Output

The output of the `gto_fastq_complement` program is the FASTQ file with the ACGT base complements. Using the input above, an output example for this is the following:

```
@SRR001666.1 071112_SLXA-EAS1_s_7:5:1:817:345 length=72
CCCACTACCGGCGACGGCTACCGCAGTTTAGGGTGGTTCAATGGGAATTGTTGAATTCCCAAAAGTTTATCT
+SRR001666.1 071112_SLXA-EAS1_s_7:5:1:817:345 length=72
IIIIIIIIIIIIIIIIIIIIIIIIIIIIII9IG9ICIIIIIIIIIIIIIIIIIIIIDIIIIIII>IIIIII/
@SRR001666.2 071112_SLXA-EAS1_s_7:5:1:801:338 length=72
CAAGTCCCTATGCTGCAAACATAAAATTCTTAGACTTCGTCTTCAGCTACTATTATGCGCAGCAAAATAGTA
+SRR001666.2 071112_SLXA-EAS1_s_7:5:1:801:338 length=72
IIIIIIIIIIIIIIIIIIIIIIIIIIIIIIII6IBIIIIIIIIIIIIIIIIIIIIIIIGII>IIIII-I)8I
```

#### 2.24 Program gto_fastq_reverse

The `gto_fastq_reverse` reverses the ACGT bases order for each read in a FASTQ file format.
For help type:

```
./gto_fastq_reverse -h
```

In the following subsections, we explain the input and output paramters.

##### Input parameters

The `gto_fastq_reverse` program needs two streams for the computation, namely the input and output standard. The input stream is a FASTQ file.
The attribution is given according to:

```
Usage: ./gto_fastq_reverse [options] [[--] args]
   or: ./gto_fastq_reverse [options]

It reverses the ACGT bases order for each read in a FASTQ file.

    -h, --help            Show this help message and exit

Basic options
    < input.fastq         Input FASTQ file (stdin)
    > output.fastq        Output FASTQ file (stdout)

Example: ./gto_fastq_reverse < input.fastq > output.fastq
```

An example of such an input file is:

```
@SRR001666.1 071112_SLXA-EAS1_s_7:5:1:817:345 length=72
GGGTGATGGCCGCTGCCGATGGCGTCAAATCCCACCAAGTTACCCTTAACAACTTAAGGGTTTTCAAATAGA
+SRR001666.1 071112_SLXA-EAS1_s_7:5:1:817:345 length=72
IIIIIIIIIIIIIIIIIIIIIIIIIIIIII9IG9ICIIIIIIIIIIIIIIIIIIIIDIIIIIII>IIIIII/
@SRR001666.2 071112_SLXA-EAS1_s_7:5:1:801:338 length=72
GTTCAGGGATACGACGTTTGTATTTTAAGAATCTGAAGCAGAAGTCGATGATAATACGCGTCGTTTTATCAT
+SRR001666.2 071112_SLXA-EAS1_s_7:5:1:801:338 length=72
IIIIIIIIIIIIIIIIIIIIIIIIIIIIIIII6IBIIIIIIIIIIIIIIIIIIIIIIIGII>IIIII-I)8I
```

##### Output

The output of the gto_fastq_reverse program is the FASTQ file complement with the flag "(Reversed)" added in the header.
Using the input above, an output example for this is the following:

```
@SRR001666.1 071112_SLXA-EAS1_s_7:5:1:817:345 length=72 (Reversed)
AGATAAACTTTTGGGAATTCAACAATTCCCATTGAACCACCCTAAACTGCGGTAGCCGTCGCCGGTAGTGGG
+
/IIIIII>IIIIIIDIIIIIIIIIIIIIIIIIIIICI9GI9IIIIIIIIIIIIIIIIIIIIIIIIIIIII
@SRR001666.2 071112_SLXA-EAS1_s_7:5:1:801:338 length=72 (Reversed)
TACTATTTTGCTGCGCATAATAGTAGCTGAAGACGAAGTCTAAGAATTTTATGTTTGCAGCATAGGGACTTG
+
I8)I-IIIII>IIGIIIIIIIIIIIIIIIIIIIIIIIIBI6IIIIIIIIIIIIIIIIIIIIIIIIIIIIIII
```

#### 2.25 Program gto_fastq_variation_map

The gto_fastq_variation_map identifies the variation that occours in the sequences relative to the reads or a set of reads.
For help type:

```
./gto_fastq_variation_map -h
```

In the following subsections, we explain the input and output paramters.

##### Input parameters

The gto_fastq_variation_map program needs FASTQ, FASTA or SEQ files to be used as reference and target files.
The attribution is given according to:

```
Usage: ./gto_fastq_variation_map <OPTIONS>... [FILE]:<...> [FILE]:<...>
./gto_fastq_variation_map: a tool to map relative singularity regions
The (probabilistic) Bloom filter is automatically set.

  -v                         verbose mode,
  -a                         about CHESTER,
  -s <value>                 bloom size,
  -i                         use inversions,
  -p                         show positions/words,
  -k <value>                 k-mer size (up to 30),

  [rFile1]:<rFile2>:<...>    reference file(s),
  [tFile1]:<tFile2>:<...>    target file(s).

The reference files may be FASTA, FASTQ or DNA SEQ,
while the target files may be FASTA OR DNA SEQ.
Report bugs to <{pratas,raquelsilva,ap,pjf}@ua.pt>.
```

An example of a reference file (Multi-FASTA format) is:

```
>AB000264 |acc=AB000264|descr=Homo sapiens mRNA
ACAAGACGGCCTCCTGCTGCTGCTGCTCTCCGGGGCCACGGCCCTGGAGGGTCCACCGCTGCCCTGCTGCCATTGTCCC
CGGCCCCACCTAAGGAAAAGCAGCCTCCTGACTTTCCTCGCTTGGGCCGAGACAGCGAGCATATGCAGGAAGCGGCAGG
AAGTGGTTTGAGTGGACCTCCGGGCCCCTCATAGGAGAGGAAGCTCGGGAGGTGGCCAGGCGGCAGGAAGCAGGCCAGT
GCCGCGAATCCGCGCGCCGGGACAGAATCTCCTGCAAAGCCCTGCAGGAACTTCTTCTGGAAGACCTTCTCCACCCCCC
CAGCTAAAACCTCACCCATGAATGCTCACGCAAGTTTAATTACAGACCTGAA
>AB000263 |acc=AB000263|descr=Homo sapiens mRNA
ACAAGATGCCATTGTCCCCCGGCCTCCTGCTGCTGCTGCTCTCCGGGGCCACGGCCACCGCTGCCCTGCCCCTGGAGGG
TGGCCCCACCGGCCGAGACAGCGAGCATATGCAGGAAGCGGCAGGAATAAGGAAAAGCAGCCTCCTGACTTTCCTCGCT
TGGTGGTTTGAGTGGACCTCCCAGGCCAGTGCCGGGCCCCTCATAGGAGAGGAAGCTCGGGAGGTGGCCAGGCGGCAGG
AAGGCGCACCCCCCCAGCAATCCGCGCGCCGGGACAGAATGCCCTGCAGGAACTTCTTCTGGAAGACCTTCTCCTCCTG
CAAATAAAACCTCACCCATGAATGCTCACGCAAGTTTAATTACAGACCTGAA
```

An example for the target file (FASTQ format) is:

```
@SRR001666.1 071112_SLXA-EAS1_s_7:5:1:817:345 length=60
GGGTGATGGCCGCTGCCGATGGCGTCAAATCCCACCAAGTTACCCTTAACAACTTAAGGG
+
IIIIIIIIIIIIIIIIIIIIIIIIIIIIII9IG9ICIIIIIIIIIIIIIIIIIIIIDIII
@SRR001666.2 071112_SLXA-EAS1_s_7:5:1:801:338 length=72
GTTCAGGGATACGACGTTTGTATTTTAAGAATCTGAAGCAGAAGTCGATGATAATACGCGTCGTTTTATCAT
+
IIIIIIIIIIIIIIIIIIIIIIIIIIIIIIII6IBIIIIIIIIIIIIIIIIIIIIIIIGII>IIIII-I)8I
```

##### Output

The output of the gto_fastq_variation_map program is a text file identifying the relative regions. Using the inputs above, an output example for this is the following:

```
11111111111111111111111111111000000000000000000000000000000001111111111111111111
1111111111000000000000000000000000000000000000000000
```

#### 2.26 Program gto_fastq_variation_filter

The gto_fastq_variation_filter filters and segments the regions of singularity from the output of gto_fastq_variation_map.
For help type:

```
./gto_fastq_variation_filter -h
```

In the following subsections, we explain the input and output paramters.

##### Input parameters

The gto_fastq_variation_filter program needs a the output of gto_fastq_variation_map to compute. The attribution is given according to:

```
Usage: ./gto_fastq_variation_filter <OPTIONS>... [FILE]:<...>
./gto_fastq_variation_filter: a tool to filter maps (gto_fastq_variation_map)

  -v                          verbose mode,
  -a                          about CHESTER,
  -t <value>                  threshold [0.0;1.0],
  -w <value>                  window size,
  -u <value>                  sub-sampling,

  [tFile1]:<tFile2>:<...>  target file(s).

The target files may be generated by gto_fastq_variation_map.
Report bugs to <{pratas,raquelsilva,ap,pjf}@ua.pt>.
```

An example of such an input file is:

```
1111111111111111111111111111100000000000000000000000000000001111111111111111111
11111111110000000000000000000000000000000000000000000
```

##### Output

The output of the `gto_fastq_variation_filter` program is a text file with the coordenates of the segmented regions.
Using the inputs above, an output example for this is the following:

```
#132#132
30:60
90:130
```

#### 2.27 Program gto_fastq_variation_visual

The `gto_fastq_variation_visual` depites the regions of singularity using the output from `gto_fastq_variation_filt` into an SVG image.
For help type:

```
./gto_fastq_variation_visual -h
```

In the following subsections, we explain the input and output paramters.

##### Input parameters

The `gto_fastq_variation_visual` program needs a the output of `gto_fastq_variation_filter` to compute.
The attribution is given according to:

```
Usage: ./gto_fastq_variation_visual <OPTIONS>... [FILE]:<...>
./gto_fastq_variation_visual: visualize relative singularity regions.

  -v                         verbose mode,
  -a                         about CHESTER,
  -e <value>                 enlarge painted regions,

  [tFile1]:<tFile2>:<...>    target file(s).

Report bugs to <{pratas,raquelsilva,ap,pjf}@ua.pt>.
```

An example of such an input file is:

```
#132#132
30:60
90:130
```

##### Output

The output of the `gto_fastq_variation_visual` program is a SVG plot with the maps.
In the Figure 2.1 is represented the plot using the input above.

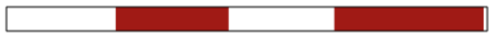

Figure 2.1: `gto_fastq_variation_visual` execution plot.

#### 2.28 Program gto_fastq_metagenomics

The `gto_fastq_metagenomics` is an ultra-fast method to infer metagenomic composition of sequenced reads relative to a database. gto_fastq_metagenomics measures similarity between any FASTQ file (or FASTA), independently from the size, against any multi-FASTA database, such as the entire set of complete genomes from the NCBI. gto_fastq_metagenomics supports single reads, paired-end reads, and compositions of both. It has been tested in many plataforms, such as Illumina MySeq, HiSeq, Novaseq, IonTorrent.
gto_fastq_metagenomics is efficient to detect the presence and authenticate a given species in the FASTQ reads. The core of the method is based on relative data compression. gto_fastq_metagenomics uses variable multi-threading, without multiplying the memory for each thread, being able to run efficiently in a common laptop.
For help type:

```
./gto_fastq_metagenomics -h
```

In the following subsections, we explain the input and output paramters.

##### Input parameters

The `gto_fastq_metagenomics` program needs a FASTQ file to compute.

The attribution is given according to:

```
NAME
      gto_fastq_metagenomics v3.1: a tool to infer metagenomic composition.

SYNOPSIS
      gto_fastq_metagenomics [OPTION]... [FILE1]:[FILE2]:... [FILE]

SAMPLE
      gto_fastq_metagenomics -v -F -l 47 -Z -y pro.com reads1.fq:reads2.fq DB.fa

DESCRIPTION
      It infers metagenomic sample composition of sequenced reads.
      The core of the method uses a cooperation between multiple
      context and tolerant context models with several depths.
      The reference sequences must be in a multi-FASTA format.
      The sequenced reads must be trimmed and in FASTQ format.

      Non-mandatory arguments:

      -h                   give this help,
      -F                   force mode (overwrites top file),
      -V                   display version number,
      -v                   verbose mode (more information),
      -Z                   database local similarity,
      -s                   show compression levels,

      -l <level>           compression level [1;47],
      -p <sample>          subsampling (default: 1),
      -t <top>             top of similarity (default: 20),
      -n <nThreads>        number of threads (default: 2),

      -x <FILE>            similarity top filename,
      -y <FILE>            profile filename (-Z must be on).

      Mandatory arguments:

      [FILE1]:[FILE2]:...  metagenomic filename (FASTQ),
                           Use ":" for splitting files.

      [FILE]               database filename (Multi-FASTA).

COPYRIGHT
      Copyright (C) 2014-2019, IEETA, University of Aveiro.
      This is a Free software, under GPLv3. You may redistribute
      copies of it under the terms of the GNU - General Public
      License v3 <http://www.gnu.org/licenses/gpl.html>.
```

An example of such an input file is:

```
@SRR001666.1 071112_SLXA-EAS1_s_7:5:1:817:345 length=72
GGGTGATGGCCGCTGCCGATGGCGTCAAATCCCACCAAGTTACCCTTAACAACTTAAGGGTTTTCAAATAGA
+SRR001666.1 071112_SLXA-EAS1_s_7:5:1:817:345 length=72
IIIIIIIIIIIIIIIIIIIIIIIIIIIIII9IG9ICIIIIIIIIIIIIIIIIIIIIDIIIIIII>IIIIII/
@SRR001666.2 071112_SLXA-EAS1_s_7:5:1:801:338 length=72
GTTCAGGGATACGACGTTTGTATTTTAAGAATCTGAAGCAGAAGTCGATGATAATACGCGTCGTTTTATCAT
+SRR001666.2 071112_SLXA-EAS1_s_7:5:1:801:338 length=72
IIIIIIIIIIIIIIIIIIIIIIIIIIIIIIII6IBIIIIIIIIIIIIIIIIIIIIIIIGII>IIIII-I)8I
```

#### Output

The output of the `gto_fastq_metagenomics` program is a CSV file (top.csv) with the highest probability of being contained in the samples. An example for this CSV file is the following:

```
1  66725   12.263   NC_037703.1_Saccharomycodes_ludwigii_strain_Y-8871_mitochondrion
2  66725   12.263   NC_037703.1_Saccharomycodes_ludwigii_strain_Y-8871_mitochondrion
3  107123  11.492   NC_012621.1_Nakaseomyces_bacillisporus_mitochondrion
4  107123  11.492   NC_012621.1_Nakaseomyces_bacillisporus_mitochondrion
5  16592   11.153   NC_024030.1_Equus_przewalskii_mitochondrial_DNA
6  14583   10.851   NC_021120.1_Bursaphelenchus_mucronatus_mitochondrion
7  162504  10.607   NC_018415.1_Candidatus_Carsonella_ruddii_CS_isolate_Thao2000
8  10315   10.586   NC_016117.1_Mnemiopsis_leidyi_mitochondrion
9  162589  10.550   NC_018414.1_Candidatus_Carsonella_ruddii_CE_isolate_Thao2000
10 166163  10.476   NC_018416.1_Candidatus_Carsonella_ruddii_HC_isolate_Thao2000
```

### Chapter 3

### FASTA tools

The FASTA tool subset has similar goals to the FASTQ tools. With these tools, it is possible convert data from different formats to the FASTA and multi-FASTA files, or the opposite. In these tools, there are also features to extract and filter reads based on patterns, which can solve specific problems in genomic analytic workflows. The currently available FASTA tools, for analysis and manipulation, are:

1. `gto_fasta_to_seq`: it converts a FASTA or Multi-FASTA file format to a seq.
2. `gto_fasta_from_seq`: it converts a genomic sequence to pseudo FASTA file format.
3. `gto_fasta_extract`: it extracts sequences from a FASTA file, which the range is defined by the user in the parameters.
4. `gto_fasta_extract_by_read`: it extracts sequences from each read in a Multi-FASTA file (splited by \n), which the range is defined by the user in the parameters.
5. `gto_fasta_info`: it shows the readed information of a FASTA or Multi-FASTA file format.
6. `gto_fasta_mutate`: it reates a synthetic mutation of a fasta file given specific rates of editions, deletions and additions.
7. `gto_fasta_rand_extra_chars`: it substitues in the DNA sequence the outside ACGT chars by random ACGT symbols.
8. `gto_fasta_extract_read_by_pattern`: it extracts reads from a Multi-FASTA file format given a pattern in the header.
9. `gto_fasta_find_n_pos`: it reports the "N" regions in a sequence or FASTA (seq) file.
10. `gto_fasta_split_reads`: it splits a Multi-FASTA file to multiple FASTA files.
11. `gto_fasta_rename_human_headers`: it changes the headers of FASTA or Multi-FASTA file to simple chrX by order, where X is the number.

12. `gto_fasta_extract_pattern_coords`: it extracts the header and coordinates from a Multi-FASTA file format given a pattern/motif in the sequence.

13. `gto_fasta_complement`: it replaces the ACGT bases with their complements in FASTA or Multi-FASTA file format.

14. `gto_fasta_reverse`: it reverses the order of a FASTA or Multi-FASTA file format.

15. `gto_fasta_variation_map`: this tool is an alias to gto_fastq_variation_map tool. Please check the documentation of this tool in the in the section of FASTQ tools.

16. `gto_fasta_variation_filter`: this tool is an alias to gto_fastq_variation_filter tool. Please check the documentation of this tool in the in the section of FASTQ tools.

17. `gto_fasta_variation_visual`: this tool is an alias to gto_fastq_variation_visual tool. Please check the documentation of this tool in the in the section of FASTQ tools.

#### 3.1 Program gto_fasta_to_seq

The `gto_fasta_to_seq` converts a FASTA or Multi-FASTA file format to a sequence.
For help type:

```
./gto_fasta_to_seq -h
```

In the following subsections, we explain the input and output paramters.

##### Input parameters

The `gto_fasta_to_seq` program needs two streams for the computation, namely the input and output standard. The input stream is a FASTA or Multi-FASTA file.
The attribution is given according to:

```
Usage: ./gto_fasta_to_seq [options] [[--] args]
   or: ./gto_fasta_to_seq [options]

It converts a FASTA or Multi-FASTA file format to a seq.

    -h, --help            show this help message and exit

Basic options
    < input.fasta         Input FASTA or Multi-FASTA file format (stdin)
    > output.seq          Output sequence file (stdout)

Example: ./gto_fasta_to_seq < input.mfasta > output.seq
```

An example of such an input file is:

```
>AB000264 |acc=AB000264|descr=Homo sapiens mRNA
ACAAGACGGCCTCCTGCTGCTGCTGCTCTCCGGGGCCACGGCCCTGGAGGGTCCACCGCTGCCCTGCTGCCATTGTCCCC
GGCCCCACCTAAGGAAAAGCAGCCTCCTGACTTTCCTCGCTTGGGCCGAGACAGCGAGCATATGCAGGAAGCGGCAGGAA
GTGGTTTGAGTGGACCTCCGGGCCCCTCATAGGAGAGGAAGCTCGGGAGGTGGCCAGGCGGCAGGAAGCAGGCCAGTGCC
GCGAATCCGCGCGCCGGGACAGAATCTCCTGCAAAGCCCTGCAGGAACTTCTTCTGGAAGACCTTCTCCACCCCCCCAGC
TAAAACCTCACCCATGAATGCTCACGCAAGTTTAATTACAGACCTGAA
>AB000263 |acc=AB000263|descr=Homo sapiens mRNA
ACAAGATGCCATTGTCCCCCGGCCTCCTGCTGCTGCTGCTCTCCGGGGCCACGGCCACCGCTGCCCTGCCCCTGGAGGGT
GGCCCCACCGGCCGAGACAGCGAGCATATGCAGGAAGCGGCAGGAATAAGGAAAAGCAGCCTCCTGACTTTCCTCGCTTG
GTGGTTTGAGTGGACCTCCCAGGCCAGTGCCGGGCCCCTCATAGGAGAGGAAGCTCGGGAGGTGGCCAGGCGGCAGGAAG
GCGCACCCCCCCAGCAATCCGCGCGCCGGGACAGAATGCCCTGCAGGAACTTCTTCTGGAAGACCTTCTCCTCCTGCAAA
TAAAACCTCACCCATGAATGCTCACGCAAGTTTAATTACAGACCTGAA
```

##### Output

The output of the `gto_fasta_to_seq` program is a group sequence.
Using the input above, an output example for this is the following:

```
ACAAGACGGCCTCCTGCTGCTGCTGCTCTCCGGGGCCACGGCCCTGGAGGGTCCACCGCTGCCCTGCTGCCATTGTCCCC
GGCCCCACCTAAGGAAAAGCAGCCTCCTGACTTTCCTCGCTTGGGCCGAGACAGCGAGCATATGCAGGAAGCGGCAGGAA
GTGGTTTGAGTGGACCTCCGGGCCCCTCATAGGAGAGGAAGCTCGGGAGGTGGCCAGGCGGCAGGAAGCAGGCCAGTGCC
GCGAATCCGCGCGCCGGGACAGAATCTCCTGCAAAGCCCTGCAGGAACTTCTTCTGGAAGACCTTCTCCACCCCCCCAGC
TAAAACCTCACCCATGAATGCTCACGCAAGTTTAATTACAGACCTGAAACAAGATGCCATTGTCCCCCGGCCTCCTGCTG
CTGCTGCTCTCCGGGGCCACGGCCACCGCTGCCCTGCCCCTGGAGGGTGGCCCCACCGGCCGAGACAGCGAGCATATGCA
GGAAGCGGCAGGAATAAGGAAAAGCAGCCTCCTGACTTTCCTCGCTTGGTGGTTTGAGTGGACCTCCCAGGCCAGTGCCG
GGCCCCTCATAGGAGAGGAAGCTCGGGAGGTGGCCAGGCGGCAGGAAGGCGCACCCCCCCAGCAATCCGCGCGCCGGGAC
AGAATGCCCTGCAGGAACTTCTTCTGGAAGACCTTCTCCTCCTGCAAATAAAACCTCACCCATGAATGCTCACGCAAGTT
TAATTACAGACCTGAA
```

#### 3.2 Program gto_fasta_from_seq

The `gto_fasta_from_seq` converts a genomic sequence to pseudo FASTA file format.
For help type:

```
./gto_fasta_from_seq -h
```

In the following subsections, we explain the input and output paramters.

##### Input parameters

The `gto_fasta_from_seq` program needs two streams for the computation, namely the input and output standard. The input stream is a sequence group file.
The attribution is given according to:

```
Usage: ./gto_fasta_from_seq [options] [[--] args]
   or: ./gto_fasta_from_seq [options]

It converts a genomic sequence to pseudo FASTA file format.
```

```
    -h, --help             show this help message and exit

Basic options
    < input.seq            Input sequence file (stdin)
    > output.fasta         Output FASTA file format (stdout)

Optional options
    -n, --name=<str>       The read's header
    -l, --lineSize=<int>   The maximum of chars for line

Example: ./gto_fasta_from_seq -l <lineSize> -n <name> < input.seq > output.fasta
```

An example of such an input file is:

```
ACAAGACGGCCTCCTGCTGCTGCTGCTCTCCGGGGCCACGGCCCTGGAGGGTCCACCGCTGCCCTGCTGCCATTGTCCCC
GGCCCCACCTAAGGAAAAGCAGCCTCCTGACTTTCCTCGCTTGGGCCGAGACAGCGAGCATATGCAGGAAGCGGCAGGAA
GTGGTTTGAGTGGACCTCCGGGCCCCTCATAGGAGAGGAAGCTCGGGAGGTGGCCAGGCGGCAGGAAGCAGGCCAGTGCC
GCGAATCCGCGCGCCGGGACAGAATCTCCTGCAAAGCCCTGCAGGAACTTCTTCTGGAAGACCTTCTCCACCCCCCCAGC
TAAAACCTCACCCATGAATGCTCACGCAAGTTTAATTACAGACCTGAAACAAGATGCCATTGTCCCCCGGCCTCCTGCTG
CTGCTGCTCTCCGGGGCCACGGCCACCGCTGCCCTGCCCCTGGAGGGTGGCCCCACCGGCCGAGACAGCGAGCATATGCA
GGAAGCGGCAGGAATAAGGAAAAGCAGCCTCCTGACTTTCCTCGCTTGGTGGTTTGAGTGGACCTCCCAGGCCAGTGCCG
GGCCCCTCATAGGAGAGGAAGCTCGGGAGGTGGCCAGGCGGCAGGAAGGCGCACCCCCCCAGCAATCCGCGCGCCGGGAC
AGAATGCCCTGCAGGAACTTCTTCTGGAAGACCTTCTCCTCCTGCAAATAAAACCTCACCCATGAATGCTCACGCAAGTT
TAATTACAGACCTGAA
```

##### Output

The output of the `gto_fasta_from_seq` program is a pseudo FASTA file.
Using the input above with the size line as 80 and the read's header as ”SeqToFasta”, an output example for this is the following:

```
>SeqToFasta
ACAAGACGGCCTCCTGCTGCTGCTGCTCTCCGGGGCCACGGCCCTGGAGGGTCCACCGCTGCCCTGCTGCCATTGTCCCC
GGCCCCACCTAAGGAAAAGCAGCCTCCTGACTTTCCTCGCTTGGGCCGAGACAGCGAGCATATGCAGGAAGCGGCAGGAA
GTGGTTTGAGTGGACCTCCGGGCCCCTCATAGGAGAGGAAGCTCGGGAGGTGGCCAGGCGGCAGGAAGCAGGCCAGTGCC
GCGAATCCGCGCGCCGGGACAGAATCTCCTGCAAAGCCCTGCAGGAACTTCTTCTGGAAGACCTTCTCCACCCCCCCAGC
TAAAACCTCACCCATGAATGCTCACGCAAGTTTAATTACAGACCTGAAACAAGATGCCATTGTCCCCCGGCCTCCTGCTG
CTGCTGCTCTCCGGGGCCACGGCCACCGCTGCCCTGCCCCTGGAGGGTGGCCCCACCGGCCGAGACAGCGAGCATATGCA
GGAAGCGGCAGGAATAAGGAAAAGCAGCCTCCTGACTTTCCTCGCTTGGTGGTTTGAGTGGACCTCCCAGGCCAGTGCCG
GGCCCCTCATAGGAGAGGAAGCTCGGGAGGTGGCCAGGCGGCAGGAAGGCGCACCCCCCCAGCAATCCGCGCGCCGGGAC
AGAATGCCCTGCAGGAACTTCTTCTGGAAGACCTTCTCCTCCTGCAAATAAAACCTCACCCATGAATGCTCACGCAAGTT
TAATTACAGACCTGAA
```

#### 3.3 Program gto_fasta_extract

The `gto_fasta_extract` extracts sequences from a FASTA file, which the range is defined by the user in the parameters.
For help type:

```
./gto_fasta_extract -h
```

In the following subsections, we explain the input and output paramters.

##### Input parameters

The `gto_fasta_extract` program needs two paramenters, which defines the begin and the end of the extraction, and two streams for the computation, namely the input and output standard. The input stream is a FASTA file.
The attribution is given according to:

```
Usage: ./gto_fasta_extract [options] [[--] args]
   or: ./gto_fasta_extract [options]

It extracts sequences from a FASTA file.

    -h, --help             show this help message and exit

Basic options
    -i, --init=<int>       The first position to start the extraction (default 0)
    -e, --end=<int>        The last extract position (default 100)
    < input.fasta          Input FASTA or Multi-FASTA file format (stdin)
    > output.seq           Output sequence file (stdout)

Example: ./gto_fasta_extract -i <init> -e <end> < input.fasta > output.seq
```

An example of such an input file is:

```
>AB000264 |acc=AB000264|descr=Homo sapiens mRNA
ACAAGACGGCCTCCTGCTGCTGCTGCTCTCCGGGGCCACGGCCCTGGAGGGTCCACCGCTGCCCTGCTGCCATTGTCCCC
GGCCCCACCTAAGGAAAAGCAGCCTCCTGACTTTCCTCGCTTGGGCCGAGACAGCGAGCATATGCAGGAAGCGGCAGGAA
GTGGTTTGAGTGGACCTCCGGGCCCCTCATAGGAGAGGAAGCTCGGGAGGTGGCCAGGCGGCAGGAAGCAGGCCAGTGCC
GCGAATCCGCGCGCCGGGACAGAATCTCCTGCAAAGCCCTGCAGGAACTTCTTCTGGAAGACCTTCTCCACCCCCCCAGC
TAAAACCTCACCCATGAATGCTCACGCAAGTTTAATTACAGACCTGAA
```

##### Output

The output of the `gto_fasta_extract` program is a group sequence.
Using the input above with the value 0 as the extraction starting point and the 50 as the ending, an output example for this is the following:

```
ACAAGACGGCCTCCTGCTGCTGCTGCTCTCCGGGGCCACGGCCCTGGAGG
```

#### 3.4 Program gto_fasta_extract_by_read

The `gto_fasta_extract_by_read` extracts sequences from a FASTA or Multi-FASTA file, which the range is defined by the user in the parameters.

For help type:

```
./gto_fasta_extract_by_read -h
```

In the following subsections, we explain the input and output paramters.

##### Input parameters

The `gto_fasta_extract_by_read` program needs two paramenters, which defines the begin and the end of the extraction, and two streams for the computation, namely the input and output standard. The input stream is a FASTA or Multi-FASTA file.
The attribution is given according to:

```
Usage: ./gto_fasta_extract_by_read [options] [[--] args]
   or: ./gto_fasta_extract_by_read [options]

It extracts sequences from each read in a Multi-FASTA file (splited by \n)

    -h, --help              show this help message and exit

Basic options
    -i, --init=<int>        The first position to start the extraction (default 0)
    -e, --end=<int>         The last extract position (default 100)
    < input.fasta           Input FASTA or Multi-FASTA file format (stdin)
    > output.fasta          Output FASTA or Multi-FASTA file format (stdout)

Example: ./gto_fasta_extract_by_read -i <init> -e <end> < input.mfasta > output.mfasta
```

An example of such an input file is:

```
>AB000264 |acc=AB000264|descr=Homo sapiens mRNA
ACAAGACGGCCTCCTGCTGCTGCTGCTCTCCGGGGCCACGGCCCTGGAGGGTCCACCGCTGCCCTGCTGCCATTGTCCCC
GGCCCCACCTAAGGAAAAGCAGCCTCCTGACTTTCCTCGCTTGGGCCGAGACAGCGAGCATATGCAGGAAGCGGCAGGAA
GTGGTTTGAGTGGACCTCCGGGCCCCTCATAGGAGAGGAAGCTCGGGAGGTGGCCAGGCGGCAGGAAGCAGGCCAGTGCC
GCGAATCCGCGCGCCGGGACAGAATCTCCTGCAAAGCCCTGCAGGAACTTCTTCTGGAAGACCTTCTCCACCCCCCCAGC
TAAAACCTCACCCATGAATGCTCACGCAAGTTTAATTACAGACCTGAA
>AB000263 |acc=AB000263|descr=Homo sapiens mRNA
ACAAGATGCCATTGTCCCCCGGCCTCCTGCTGCTGCTGCTCTCCGGGGCCACGGCCACCGCTGCCCTGCCCCTGGAGGGT
GGCCCCACCGGCCGAGACAGCGAGCATATGCAGGAAGCGGCAGGAATAAGGAAAAGCAGCCTCCTGACTTTCCTCGCTTG
GTGGTTTGAGTGGACCTCCCAGGCCAGTGCCGGGCCCCTCATAGGAGAGGAAGCTCGGGAGGTGGCCAGGCGGCAGGAAG
GCGCACCCCCCCAGCAATCCGCGCGCCGGGACAGAATGCCCTGCAGGAACTTCTTCTGGAAGACCTTCTCCTCCTGCAAA
TAAAACCTCACCCATGAATGCTCACGCAAGTTTAATTACAGACCTGAA
```

##### Output

The output of the `gto_fasta_extract_by_read` program is FASTA or Multi-FASTA file wiht the extracted sequences.
Using the input above with the value 0 as the extraction starting point and the 50 as the ending, an output example for this is the following:

```
>AB000264 |acc=AB000264|descr=Homo sapiens mRNA
ACAAGACGGCCTCCTGCTGCTGCTGCTCTCCGGGGCCACGGCCCTGGAGG
>AB000263 |acc=AB000263|descr=Homo sapiens mRNA
ACAAGATGCCATTGTCCCCCGGCCTCCTGCTGCTGCTGCTCTCCGGGGCC
```

#### 3.5 Program gto_fasta_info

The `gto_fasta_info` shows the readed information of a FASTA or Multi-FASTA file format.
For help type:

```
./gto_fasta_info -h
```

In the following subsections, we explain the input and output paramters.

##### Input parameters

The `gto_fasta_info` program needs two streams for the computation, namely the input and output standard. The input stream is a FASTA or Multi-FASTA file.
The attribution is given according to:

```
Usage: ./gto_fasta_info [options] [[--] args]
   or: ./gto_fasta_info [options]

It shows read information of a FASTA or Multi-FASTA file format.

    -h, --help             show this help message and exit

Basic options
    < input.fasta          Input FASTA or Multi-FASTA file format (stdin)
    > output               Output read information (stdout)

Example: ./gto_fasta_info < input.mfasta > output

Output example :
Number of reads      : value
Number of bases      : value
MIN of bases in read : value
MAX of bases in read : value
AVG of bases in read : value
```

An example of such an input file is:

```
>AB000264 |acc=AB000264|descr=Homo sapiens mRNA
ACAAGACGGCCTCCTGCTGCTGCTGCTCTCCGGGGCCACGGCCCTGGAGGGTCCACCGCTGCCCTGCTGCCATTGTCCCC
GGCCCCACCTAAGGAAAAGCAGCCTCCTGACTTTCCTCGCTTGGGCCGAGACAGCGAGCATATGCAGGAAGCGGCAGGAA
GTGGTTTGAGTGGACCTCCGGGCCCCTCATAGGAGAGGAAGCTCGGGAGGTGGCCAGGCGGCAGGAAGCAGGCCAGTGCC
GCGAATCCGCGCGCCGGGACAGAATCTCCTGCAAAGCCCTGCAGGAACTTCTTCTGGAAGACCTTCTCCACCCCCCCAGC
TAAAACCTCACCCATGAATGCTCACGCAAGTTTAATTACAGACCTGAA
>AB000263 |acc=AB000263|descr=Homo sapiens mRNA
```

```
ACAAGATGCCATTGTCCCCCGGCCTCCTGCTGCTGCTGCTCTCCGGGGCCACGGCCACCGCTGCCCTGCCCCTGGAGGGT
GGCCCCACCGGCCGAGACAGCGAGCATATGCAGGAAGCGGCAGGAATAAGGAAAAGCAGCCTCCTGACTTTCCTCGCTTG
GTGGTTTGAGTGGACCTCCCAGGCCAGTGCCGGGCCCCTCATAGGAGAGGAAGCTCGGGAGGTGGCCAGGCGGCAGGAAG
GCGCACCCCCCCAGCAATCCGCGCGCCGGGACAGAATGCCCTGCAGGAACTTCTTCTGGAAGACCTTCTCCTCCTGCAAA
TAAAACCTCACCCATGAATGCTCACGCAAGTTTAATTACAGACCTGAA
```

##### Output

The output of the `gto_fasta_info` program is a set of information related to the file read.
Using the input above, an output example for this is the following:

```
Number of reads      : 2
Number of bases      : 736
MIN of bases in read : 368
MAX of bases in read : 368
AVG of bases in read : 368.0000
```

#### 3.6 Program gto_fasta_mutate

The `gto_fasta_mutate` creates a synthetic mutation of a FASTA file given specific rates of editions, deletions and additions. All these paramenters are defined by the user, and their are optional.
For help type:

```
./gto_fasta_mutate -h
```

In the following subsections, we explain the input and output paramters.

##### Input parameters

The `gto_fasta_mutate` program needs two streams for the computation, namely the input and output standard. However, optional settings can be supplied too, such as the starting point to the random generator, and the edition, deletion and insertion rates. Also, the user can choose to use the ACGTN alphabet in the synthetic mutation. The input stream is a FASTA or Multi-FASTA File.
The attribution is given according to:

```
Usage: ./gto_fasta_mutate [options] [[--] args]
   or: ./gto_fasta_mutate [options]

Creates a synthetic mutation of a fasta file given specific rates of editions,
deletions and additions

    -h, --help                    show this help message and exit

Basic options
    < input.fasta                 Input FASTA or Multi-FASTA file format (stdin)
    > output.fasta                Output FASTA or Multi-FASTA file format (stdout)

Optional
```

```
    -s, --seed=<int>              Starting point to the random generator
    -e, --edit-rate=<dbl>         Defines the edition rate (default 0.0)
    -d, --deletion-rate=<dbl>     Defines the deletion rate (default 0.0)
    -i, --insertion-rate=<dbl>    Defines the insertion rate (default 0.0)
    -a, --ACGTN-alphabet          When active, the application uses the ACGTN alphabet

Example: ./gto_fasta_mutate -s <seed> -e <edit rate> -d <deletion rate> -i
<insertion rate> -a < input.mfasta > output.fasta
```

An example of such an input file is:

```
>AB000264 |acc=AB000264|descr=Homo sapiens mRNA
ACAAGACGGCCTCCTGCTGCTGCTGCTCTCCGGGGCCACGGCCCTGGAGGGTCCACCGCTGCCCTGCTGCCATTGTCCCC
GGCCCCACCTAAGGAAAAGCAGCCTCCTGACTTTCCTCGCTTGGGCCGAGACAGCGAGCATATGCAGGAAGCGGCAGGAA
GTGGTTTGAGTGGACCTCCGGGCCCCTCATAGGAGAGGAAGCTCGGGAGGTGGCCAGGCGGCAGGAAGCAGGCCAGTGCC
GCGAATCCGCGCGCCGGGACAGAATCTCCTGCAAAGCCCTGCAGGAACTTCTTCTGGAAGACCTTCTCCACCCCCCCAGC
TAAAACCTCACCCATGAATGCTCACGCAAGTTTAATTACAGACCTGAA
>AB000263 |acc=AB000263|descr=Homo sapiens mRNA
ACAAGATGCCATTGTCCCCCGGCCTCCTGCTGCTGCTGCTCTCCGGGGCCACGGCCACCGCTGCCCTGCCCCTGGAGGGT
GGCCCCACCGGCCGAGACAGCGAGCATATGCAGGAAGCGGCAGGAATAAGGAAAAGCAGCCTCCTGACTTTCCTCGCTTG
GTGGTTTGAGTGGACCTCCCAGGCCAGTGCCGGGCCCCTCATAGGAGAGGAAGCTCGGGAGGTGGCCAGGCGGCAGGAAG
GCGCACCCCCCCAGCAATCCGCGCGCCGGGACAGAATGCCCTGCAGGAACTTCTTCTGGAAGACCTTCTCCTCCTGCAAA
TAAAACCTCACCCATGAATGCTCACGCAAGTTTAATTACAGACCTGAA
```

##### Output

The output of the `gto_fasta_mutate` program is a FASTA or Multi-FASTA file whith the synthetic mutation of input file.
Using the input above with the seed value as 1 and the edition rate as 0.5, an output example for this is the following:

```
>AB000264 |acc=AB000264|descr=Homo sapiens mRNA
ACGCAACGNATTCCTGCTGATCATANTGTNCCGCNCCCCNGCGACGGGGNCTCNCNNGCACACATNGTACCATTGTCCAC
NCTTNCANGTNANCGCTAGCAGGCTACNGTTTNTCCTCNCCTANNCCAANCNGGCGTNNNTACACTGGCACGTGCAGGCA
TNGGTCGGCNGGNNCCTCCGGNAACGGCACCGGAGACGAAGCTCGGNGGNTATACAGGTGTCANGAAACATCCCCGCGNC
GNGTGNCCNNGAANCCANAGAGTATCTCACTCACAACCCTGCGTGCACNTCTAGAGNANGACCTTACNCACCNTCCCNTT
NNGTACCACACCAATGAACGCTGCAGAAAGTCTGTTTNNAGGNGNGCA
>AB000263 |acc=AB000263|descr=Homo sapiens mRNA
ATTTGAAGGCAANCGGNCCAGNAATNCGGNGGGTGCNGCTCNTGTNGGCTACGGNCATCGCGGCCCTGCTNTANTAAGCN
TGAACCACCGNTCGNNGCACTTAGCAATNGCGNAANCCGTCGGCACGGCGGAGACNAANCCGCTANTNNTTTCCCGCTNA
ATGGNTGTACAAGACCNACTANACCANCCTCCGTCACCACACTGGAGCGCANGATGGNNCGCTGNCTAGNAGNCNNTGAG
GCGCTCCNTCCTANAAANCCGTGGNCGAGCNCCCTATGGNAGNGTGGGGGTTTTACCGGAAGACCNTCGNGCCCTATGGG
AGCAATCANAANCTAGAAAGCTTACNGATGGTGANGAANTAGACTANG
```

#### 3.7 Program gto_fasta_rand_extra_chars

The `gto_fasta_rand_extra_chars` substitues in the DNA sequence the outside ACGT chars by random ACGT symbols. It works both in FASTA and Multi-FASTA file formats.
For help type:

```
./gto_fasta_rand_extra_chars -h
```

In the following subsections, we explain the input and output paramters.

##### Input parameters

The `gto_fasta_rand_extra_chars` program needs two streams for the computation, namely the input and output standard. The input stream is a FASTA or Multi-FASTA file.
The attribution is given according to:

```
Usage: ./gto_fasta_rand_extra_chars [options] [[--] args]
   or: ./gto_fasta_rand_extra_chars [options]

It substitues in the DNA sequence the outside ACGT chars by random ACGT symbols.
It works both in FASTA and Multi-FASTA file formats

    -h, --help            show this help message and exit

Basic options
    < input.fasta         Input FASTA or Multi-FASTA file format (stdin)
    > output.fasta        Output FASTA or Multi-FASTA file format (stdout)

Example: ./gto_fasta_rand_extra_chars < input.mfasta > output.mfasta
```

An example of such an input file is:

```
>AB000264 |acc=AB000264|descr=Homo sapiens mRNA
ANAAGACGGCCTCCTGCTGCTGCTGCTCTCCGGGGCCACGNCCCTGGAGGGTCCNCCGCTGCCCTGCTGCCATTGNCNCC
NGCCCCACCTAAGGAAAAGCAGCCTCCTGACTTTCCTCGCTTGGGCCGAGACAGCGAGCATATGCNGGAAGCGGCAGGAA
GNGGTTTGAGTGGACCTCCNGGCCCCTCATAGGAGAGGAAGCNNGGGAGGTGGCCAGGCGGCAGGAAGCAGGCCAGTGNC
GCGAATCCGNGCGCCGGGACAGAATCTCCTGCAAAGCCCTGCAGGAACTTCTTCTGGAAGACCTTCTCCACCCCCCCNNN
TAAANNNTCACCCATGAATGCTCACGCAANTTTAATTACAGACCTGAA
>AB000263 |acc=AB000263|descr=Homo sapiens mRNA
GCGAATCCGNGCGCCGGGACAGAATCTCCTTCTCCACCCCCCCNNNTGCAAAGCCCTGCAGGAACTTCTTCTGGAAGACC
NGCCCCACCTAAGGAAAAGCAGCCTCCAGGAACTGACTTTCCTCGCTTGGGCCGAGACAGCGAGCATATGCNGGAAGCGG
ANAAGACGGCCTCCTGCTGCTGCTGCTCTCCGGGGCCACGNCCCTGGCNCCAGGGTCCNCCGCTGCCCTGCTGCCATTGN
GAGGAAGCNNGGGAGGTGGCCAGGCGGCAGGAAGCAGGCCAGTGNCGNGGTTTGAGTGGACCTCCNGGCCCCTCATAGGA
TCACGCAANTTTAATTACAGACCTGAATAAANNNTCACCCATGAATGC
```

##### Output

The output of the `gto_fasta_rand_extra_chars` program is a FASTA or Multi-FASTA file.
Using the input above, an output example for this is the following:

```
>AB000264 |acc=AB000264|descr=Homo sapiens mRNA
ATAAGACGGCCTCCTGCTGCTGCTGCTCTCCGGGGCCACGGCCCTGGAGGGTCCCCCGCTGCCCTGCTGCCATTGTCCCC
TGCCCCACCTAAGGAAAAGCAGCCTCCTGACTTTCCTCGCTTGGGCCGAGACAGCGAGCATATGCGGGAAGCGGCAGGAA
GAGGTTTGAGTGGACCTCCCGGCCCCTCATAGGAGAGGAAGCCGGGGAGGTGGCCAGGCGGCAGGAAGCAGGCCAGTGTC
GCGAATCCGGGCGCCGGGACAGAATCTCCTGCAAAGCCCTGCAGGAACTTCTTCTGGAAGACCTTCTCCACCCCCCCTTG
```

```
TAAAAGATCACCCATGAATGCTCACGCAAATTTAATTACAGACCTGAA
>AB000263 |acc=AB000263|descr=Homo sapiens mRNA
GCGAATCCGTGCGCCGGGACAGAATCTCCTTCTCCACCCCCCCATCTGCAAAGCCCTGCAGGAACTTCTTCTGGAAGACC
GGCCCCACCTAAGGAAAAGCAGCCTCCAGGAACTGACTTTCCTCGCTTGGGCCGAGACAGCGAGCATATGCGGGAAGCGG
AGAAGACGGCCTCCTGCTGCTGCTGCTCTCCGGGGCCACGTCCCTGGCTCCAGGGTCCTCCGCTGCCCTGCTGCCATTGC
GAGGAAGCGGGGGAGGTGGCCAGGCGGCAGGAAGCAGGCCAGTGGCGCGGTTTGAGTGGACCTCCTGGCCCCTCATAGGA
TCACGCAACTTTAATTACAGACCTGAATAAAATGTCACCCATGAATGC
```

#### 3.8 Program gto_fasta_extract_read_by_pattern

The `gto_fasta_extract_read_by_pattern` extracts reads from a Multi-FASTA file format given a pattern in the header. Also, this pattern is case insensitive.
For help type:

```
./gto_fasta_extract_read_by_pattern -h
```

In the following subsections, we explain the input and output paramters.

##### Input parameters

The `gto_fasta_extract_read_by_pattern` program needs two streams for the computation, namely the input and output standard. The input stream is a Multi-FASTA file.
The attribution is given according to:

```
Usage: ./gto_fasta_extract_read_by_pattern [options] [[--] args]
   or: ./gto_fasta_extract_read_by_pattern [options]

It extracts reads from a Multi-FASTA file format given a pattern in the header (ID).

    -h, --help             show this help message and exit

Basic options
    -p, --pattern=<str>    Pattern to search in the file header
    < input.fasta             Input Multi-FASTA file format (stdin)
    > output.fasta            Output Multi-FASTA file format (stdout)

Example: ./gto_fasta_extract_read_by_pattern -p <pattern> < input.mfasta > output.fasta
```

An example of such an input file is:

```
>AB000264 |acc=AB000264|descr=Homo sapiens mRNA
ACAAGACGGCCTCCTGCTGCTGCTGCTCTCCGGGGCCACGGCCCTGGAGGGTCCACCGCTGCCCTGCTGCCATTGTCCCC
GGCCCCACCTAAGGAAAAGCAGCCTCCTGACTTTCCTCGCTTGGGCCGAGACAGCGAGCATATGCAGGAAGCGGCAGGAA
GTGGTTTGAGTGGACCTCCGGGCCCCTCATAGGAGAGGAAGCTCGGGAGGTGGCCAGGCGGCAGGAAGCAGGCCAGTGCC
GCGAATCCGCGCGCCGGGACAGAATCTCCTGCAAAGCCCTGCAGGAACTTCTTCTGGAAGACCTTCTCCACCCCCCCAGC
TAAAACCTCACCCATGAATGCTCACGCAAGTTTAATTACAGACCTGAA
>AB000263 |acc=AB000263|descr=Homo sapiens mRNA
ACAAGATGCCATTGTCCCCCGGCCTCCTGCTGCTGCTGCTCTCCGGGGCCACGGCCACCGCTGCCCTGCCCCTGGAGGGT
GGCCCCACCGGCCGAGACAGCGAGCATATGCAGGAAGCGGCAGGAATAAGGAAAAGCAGCCTCCTGACTTTCCTCGCTTG
GTGGTTTGAGTGGACCTCCCAGGCCAGTGCCGGGCCCCTCATAGGAGAGGAAGCTCGGGAGGTGGCCAGGCGGCAGGAAG
```

```
GCGCACCCCCCCAGCAATCCGCGCGCCGGGACAGAATGCCCTGCAGGAACTTCTTCTGGAAGACCTTCTCCTCCTGCAAA
TAAAACCTCACCCATGAATGCTCACGCAAGTTTAATTACAGACCTGAA
```

##### Output

The output of the `gto_fasta_extract_read_by_pattern` program is a Multi-FASTA file.
Using the input above with the pattern value as "264", an output example for this is the following:

```
>AB000264 |acc=AB000264|descr=Homo sapiens mRNA
ACAAGACGGCCTCCTGCTGCTGCTGCTCTCCGGGGCCACGGCCCTGGAGGGTCCACCGCTGCCCTGCTGCCATTGTCCCC
GGCCCCACCTAAGGAAAAGCAGCCTCCTGACTTTCCTCGCTTGGGCCGAGACAGCGAGCATATGCAGGAAGCGGCAGGAA
GTGGTTTGAGTGGACCTCCGGGCCCCTCATAGGAGAGGAAGCTCGGGAGGTGGCCAGGCGGCAGGAAGCAGGCCAGTGCC
GCGAATCCGCGCGCCGGGACAGAATCTCCTGCAAAGCCCTGCAGGAACTTCTTCTGGAAGACCTTCTCCACCCCCCCAGC
TAAAACCTCACCCATGAATGCTCACGCAAGTTTAATTACAGACCTGAA
```

#### 3.9 Program gto_fasta_find_n_pos

The `gto_fasta_find_n_pos` reports the "N" regions in a sequence or FASTA (seq) file.
For help type:

```
./gto_fasta_find_n_pos -h
```

In the following subsections, we explain the input and output paramters.

##### Input parameters

The `gto_fasta_find_n_pos` program needs two streams for the computation, namely the input and output standard. The input stream is a FASTA file or a sequence.
The attribution is given according to:

```
Usage: ./gto_fasta_find_n_pos [options] [[--] args]
   or: ./gto_fasta_find_n_pos [options]

It reports the 'N' regions in a sequence or FASTA (seq) file.

    -h, --help            show this help message and exit

Basic options
    < input.fasta         Input FASTQ file format or a sequence (stdin)
    > output              Output report of 'N' positions (stdout)

Example: ./gto_fasta_find_n_pos < input.fasta > output

The output obeys the following structure:
Begin   End Positions
<value> <value> <value>
```

An example of such an input file is:

```
>AB000264 |acc=AB000264|descr=Homo sapiens mRNA
NCNNNACGGCCTCCTGCTGCTGCTGCTCTCCGGGGCCACGGCCCTGGAGGGTCCACCGCTGCCCTGCTGCCATTGTCCCC
GNCCCCACCTAAGGAAAAGCAGCCTCCTGACTTTCCTCGCTTGGGCCGAGACAGCGAGCATATGCAGGAAGCGGCAGGAA
GTNGTTTGAGTGGACCTCCGGGCCCCTCATAGGAGAGGAAGCTCGGGAGGTGGCCAGGCGGCAGGAAGCAGGCCAGTGCC
GCGAATCCGCGCGCCGGGACAGAATCTCCTGCAAAGCCCTGCAGGAACNTCTTCTGGAAGACCTTCTCCACCCCCCCAGC
TAAAACCTCACCCATGAATGCTCACGCAAGTTTAATTACAGACCTGAN
```

##### Output

The output of the `gto_fasta_find_n_pos` program is a structured report of ”N” appearances in the sequence or FASTA file. The first column is the first position of the ”N” appearance, the second is the position of the last ”N” in the interval found, and the last column is the count of ”N” in this interval.
Using the input above, an output example for this is the following:

```
1   1   1
3   5   3
82  82  1
163 163 1
289 289 1
```

#### 3.10 Program gto_fasta_split_reads

The `gto_fasta_split_reads` splits a Multi-FASTA file to multiple FASTA files.
For help type:

```
./gto_fasta_split_reads -h
```

In the following subsections, we explain the input and output paramters.

##### Input parameters

The `gto_fasta_split_reads` program needs one stream for the computation, namely the input standard. This input stream is a Multi-FASTA file.
The attribution is given according to:

```
Usage: ./gto_fasta_split_reads [options] [[--] args]
   or: ./gto_fasta_split_reads [options]

It splits a Multi-FASTA file to multiple FASTA files.

    -h, --help            show this help message and exit

Basic options
    < input.fasta         Input Multi-FASTA file format (stdin)

Optional options
    -l, --location=<str>  Location to store the files
```

```
Example: ./gto_fasta_split_reads < input.mfasta
```

An example of such an input file is:

```
>AB000264 |acc=AB000264|descr=Homo sapiens mRNA
ACAAGACGGCCTCCTGCTGCTGCTGCTCTCCGGGGCCACGGCCCTGGAGGGTCCACCGCTGCCCTGCTGCCATTGTCCCC
GGCCCCACCTAAGGAAAAGCAGCCTCCTGACTTTCCTCGCTTGGGCCGAGACAGCGAGCATATGCAGGAAGCGGCAGGAA
GTGGTTTGAGTGGACCTCCGGGCCCCTCATAGGAGAGGAAGCTCGGGAGGTGGCCAGGCGGCAGGAAGCAGGCCAGTGCC
GCGAATCCGCGCGCCGGGACAGAATCTCCTGCAAAGCCCTGCAGGAACTTCTTCTGGAAGACCTTCTCCACCCCCCCAGC
TAAAACCTCACCCATGAATGCTCACGCAAGTTTAATTACAGACCTGAA
>AB000263 |acc=AB000263|descr=Homo sapiens mRNA
ACAAGATGCCATTGTCCCCCGGCCTCCTGCTGCTGCTGCTCTCCGGGGCCACGGCCACCGCTGCCCTGCCCCTGGAGGGT
GGCCCCACCGGCCGAGACAGCGAGCATATGCAGGAAGCGGCAGGAATAAGGAAAAGCAGCCTCCTGACTTTCCTCGCTTG
GTGGTTTGAGTGGACCTCCCAGGCCAGTGCCGGGCCCCTCATAGGAGAGGAAGCTCGGGAGGTGGCCAGGCGGCAGGAAG
GCGCACCCCCCCAGCAATCCGCGCGCCGGGACAGAATGCCCTGCAGGAACTTCTTCTGGAAGACCTTCTCCTCCTGCAAA
TAAAACCTCACCCATGAATGCTCACGCAAGTTTAATTACAGACCTGAA
```

##### Output

The output of the `gto_fasta_split_reads` program is a report summary of the execution, and the files created in the defined location.
Using the input above, an output example for this is the following:

```
1 : Splitting to file:./out1.fasta
2 : Splitting to file:./out2.fasta
```

#### 3.11 Program gto_fasta_rename_human_headers

The `gto_fasta_rename_human_headers` changes the headers of FASTA or Multi-FASTA file to simple chrX by order, where X is the number.
For help type:

```
./gto_fasta_rename_human_headers -h
```

In the following subsections, we explain the input and output paramters.

##### Input parameters

The `gto_fasta_rename_human_headers` program needs two streams for the computation, namely the input and output standard. The input stream is a FASTA or Multi-FASTA file.
The attribution is given according to:

```
Usage: ./gto_fasta_rename_human_headers [options] [[--] args]
   or: ./gto_fasta_rename_human_headers [options]

It changes the headers of FASTA or Multi-FASTA file to simple chr$1 by order.
```

```
    -h, --help            show this help message and exit

Basic options
    < input.fasta         Input FASTA or Multi-FASTA file format (stdin)
    > output.fasta        Output FASTA or Multi-FASTA file format (stdout)

Example: ./gto_fasta_rename_human_headers < input.mfasta > output.mfasta
```

An example of such an input file is:

```
> AB000264 | acc = AB000264 | descr = Homo sapiens mRNA
ACAAGACGGCCTCCTGCTGCTGCTGCTCTCCGGGGCCACGGCCCTGGAGGGTCCACCGCTGCCCTGCTGCCATTGTCCCC
GGCCCCACCTAAGGAAAAGCAGCCTCCTGACTTTCCTCGCTTGGGCCGAGACAGCGAGCATATGCAGGAAGCGGCAGGAA
GTGGTTTGAGTGGACCTCCGGGCCCCTCATAGGAGAGGAAGCTCGGGAGGTGGCCAGGCGGCAGGAAGCAGGCCAGTGCC
GCGAATCCGCGCGCCGGGACAGAATCTCCTGCAAAGCCCTGCAGGAACTTCTTCTGGAAGACCTTCTCCACCCCCCCAGC
TAAAACCTCACCCATGAATGCTCACGCAAGTTTAATTACAGACCTGAA
> AB000263 | acc = AB000263 | descr = Homo sapiens mRNA
ACAAGATGCCATTGTCCCCCGGCCTCCTGCTGCTGCTGCTCTCCGGGGCCACGGCCACCGCTGCCCTGCCCCTGGAGGGT
GGCCCCACCGGCCGAGACAGCGAGCATATGCAGGAAGCGGCAGGAATAAGGAAAAGCAGCCTCCTGACTTTCCTCGCTTG
GTGGTTTGAGTGGACCTCCCAGGCCAGTGCCGGGCCCCTCATAGGAGAGGAAGCTCGGGAGGTGGCCAGGCGGCAGGAAG
GCGCACCCCCCCAGCAATCCGCGCGCCGGGACAGAATGCCCTGCAGGAACTTCTTCTGGAAGACCTTCTCCTCCTGCAAA
TAAAACCTCACCCATGAATGCTCACGCAAGTTTAATTACAGACCTGAA
```

##### Output

The output of the `gto_fasta_rename_human_headers` program is a FASTA or Multi-FASTA file.
Using the input above, an output example for this is the following:

```
>chr1
ACAAGACGGCCTCCTGCTGCTGCTGCTCTCCGGGGCCACGGCCCTGGAGGGTCCACCGCTGCCCTGCTGCCATTGTCCCC
GGCCCCACCTAAGGAAAAGCAGCCTCCTGACTTTCCTCGCTTGGGCCGAGACAGCGAGCATATGCAGGAAGCGGCAGGAA
GTGGTTTGAGTGGACCTCCGGGCCCCTCATAGGAGAGGAAGCTCGGGAGGTGGCCAGGCGGCAGGAAGCAGGCCAGTGCC
GCGAATCCGCGCGCCGGGACAGAATCTCCTGCAAAGCCCTGCAGGAACTTCTTCTGGAAGACCTTCTCCACCCCCCCAGC
TAAAACCTCACCCATGAATGCTCACGCAAGTTTAATTACAGACCTGAA
>chr2
ACAAGATGCCATTGTCCCCCGGCCTCCTGCTGCTGCTGCTCTCCGGGGCCACGGCCACCGCTGCCCTGCCCCTGGAGGGT
GGCCCCACCGGCCGAGACAGCGAGCATATGCAGGAAGCGGCAGGAATAAGGAAAAGCAGCCTCCTGACTTTCCTCGCTTG
GTGGTTTGAGTGGACCTCCCAGGCCAGTGCCGGGCCCCTCATAGGAGAGGAAGCTCGGGAGGTGGCCAGGCGGCAGGAAG
GCGCACCCCCCCAGCAATCCGCGCGCCGGGACAGAATGCCCTGCAGGAACTTCTTCTGGAAGACCTTCTCCTCCTGCAAA
TAAAACCTCACCCATGAATGCTCACGCAAGTTTAATTACAGACCTGAA
```

#### 3.12 Program gto_fasta_extract_pattern_coords

The `gto_fasta_extract_pattern_coords` extracts the header and coordinates from a Multi-FASTA file format given a pattern/motif in the sequence.
For help type:

```
./gto_fasta_extract_pattern_coords -h
```

In the following subsections, we explain the input and output paramters.

##### Input parameters

The `gto_fasta_extract_pattern_coords` program needs two streams for the computation, namely the input and output standard. The input stream is a Multi-FASTA file.
The attribution is given according to:

```
Usage: ./gto_fasta_extract_pattern_coords [options] [[--] args]
   or: ./gto_fasta_extract_pattern_coords [options]

It extracts the header and coordinates from a Multi-FASTA file format given a
pattern/motif in the sequence.

    -h, --help             show this help message and exit

Basic options
    -p, --pattern=<str>    Pattern to search in the file header
    < input.fasta          Input Multi-FASTA file format (stdin)
    > output.coords        Output coordinates (stdout)

Example: ./gto_fasta_extract_pattern_coords -p <pattern> < input.fasta > output.coords
```

An example of such an input file is:

```
>AB000264 |acc=AB000264|descr=Homo sapiens mRNA
ACAAGACGGCCTCCTGCTGCTGCTGCTCTCCGGGGCCACGGCCCTGGAGGGTCCACCGCTGCCCTGCTGCCATTGTCCCC
GGCCCCACCTAAGGAAAAGCAGCCTCCTGACTTTCCTCGCTTGGGCCGAGACAGCGAGCATATGCAGGAAGCGGCAGGAA
GTGGTTTGAGTGGACCTCCGGGCCCCTCATAGGAGAGGAAGCTCGGGAGGTGGCCAGGCGGCAGGAAGCAGGCCAGTGCC
GCGAATCCGCGCGCCGGGACAGAATCTCCTGCAAAGCCCTGCAGGAACTTCTTCTGGAAGACCTTCTCCACCCCCCCAGC
TAAAACCTCACCCATGAATGCTCGCAACACGCAAGTTTAATTCGCAAGTTAGACCTGAACGGGAGGTGGCCACGCAAGTT
```

##### Output

The output of the `gto_fasta_extract_pattern_coords` program is a Multi-FASTA file.
Using the input above, with the pattern ACA, an output example for this is the following:

```
1   3   >AB000264 |acc=AB000264|descr=Homo sapiens mRNA
131 133 >AB000264 |acc=AB000264|descr=Homo sapiens mRNA
259 261 >AB000264 |acc=AB000264|descr=Homo sapiens mRNA
347 349 >AB000264 |acc=AB000264|descr=Homo sapiens mRNA
```

#### 3.13 Program gto_fasta_complement

The `gto_fasta_complement` replaces the ACGT bases with their complements in FASTA or Multi-FASTA file format.
For help type:

```
./gto_fasta_complement -h
```

In the following subsections, we explain the input and output paramters.

##### Input parameters

The `gto_fasta_complement` program needs two streams for the computation, namely the input and output standard. The input stream is a FASTA or Multi-FASTA file.
The attribution is given according to:

```
Usage: ./gto_fasta_complement [options] [[--] args]
   or: ./gto_fasta_complement [options]

It replaces the ACGT bases with their complements in FASTA or Multi-FASTA file format.

    -h, --help              Show this help message and exit

Basic options
    < input.fasta           Input FASTA or Multi-FASTA file format (stdin)
    > output.fasta          Output FASTA or Multi-FASTA file format (stdout)

Example: ./gto_fasta_complement < input.mfasta > output.mfasta
```

An example of such an input file is:

```
>AB000264 |acc=AB000264|descr=Homo sapiens mRNA
ACAAGACGGCCTCCTGCTGCTGCTGCTCTCCGGGGCCACGGCCCTGGAGGGTCCACCGCTGCCCTGCTGCCATTGTCCC
CGGCCCCACCTAAGGAAAAGCAGCCTCCTGACTTTCCTCGCTTGGGCCGAGACAGCGAGCATATGCAGGAAGCGGCAGG
AAGTGGTTTGAGTGGACCTCCGGGCCCCTCATAGGAGAGGAAGCTCGGGAGGTGGCCAGGCGGCAGGAAGCAGGCCAGT
GCCGCGAATCCGCGCGCCGGGACAGAATCTCCTGCAAAGCCCTGCAGGAACTTCTTCTGGAAGACCTTCTCCACCCCCC
CAGCTAAAACCTCACCCATGAATGCTCACGCAAGTTTAATTACAGACCTGAA
>AB000263 |acc=AB000263|descr=Homo sapiens mRNA
ACAAGATGCCATTGTCCCCCGGCCTCCTGCTGCTGCTGCTCTCCGGGGCCACGGCCACCGCTGCCCTGCCCCTGGAGGG
TGGCCCCACCGGCCGAGACAGCGAGCATATGCAGGAAGCGGCAGGAATAAGGAAAAGCAGCCTCCTGACTTTCCTCGCT
TGGTGGTTTGAGTGGACCTCCCAGGCCAGTGCCGGGCCCCTCATAGGAGAGGAAGCTCGGGAGGTGGCCAGGCGGCAGG
AAGGCGCACCCCCCCAGCAATCCGCGCGCCGGGACAGAATGCCCTGCAGGAACTTCTTCTGGAAGACCTTCTCCTCCTG
CAAATAAAACCTCACCCATGAATGCTCACGCAAGTTTAATTACAGACCTGAA
```

##### Output

The output of the `gto_fasta_complement` program is FASTA or Multi-FASTA file with the ACGT base complements.
Using the input above, an output example for this is the following:

```
>AB000264 |acc=AB000264|descr=Homo sapiens mRNA
TGTTCTGCCGGAGGACGACGACGACGAGAGGCCCCGGTGCCGGGACCTCCCAGGTGGCGACGGGACGACGGTAACAGGG
GCCGGGGTGGATTCCTTTTCGTCGGAGGACTGAAAGGAGCGAACCCGGCTCTGTCGCTCGTATACGTCCTTCGCCGTCC
TTCACCAAACTCACCTGGAGGCCCGGGGAGTATCCTCTCCTTCGAGCCCTCCACCGGTCCGCCGTCCTTCGTCCGGTCA
```

```
CGGCGCTTAGGCGCGCGGCCCTGTCTTAGAGGACGTTTCGGGACGTCCTTGAAGAAGACCTTCTGGAAGAGGTGGGGGG
GTCGATTTTGGAGTGGGTACTTACGAGTGCGTTCAAATTAATGTCTGGACTT
>AB000263 |acc=AB000263|descr=Homo sapiens mRNA
TGTTCTACGGTAACAGGGGGCCGGAGGACGACGACGACGAGAGGCCCCGGTGCCGGTGGCGACGGGACGGGGACCTCCC
ACCGGGGTGGCCGGCTCTGTCGCTCGTATACGTCCTTCGCCGTCCTTATTCCTTTTCGTCGGAGGACTGAAAGGAGCGA
ACCACCAAACTCACCTGGAGGGTCCGGTCACGGCCCGGGGAGTATCCTCTCCTTCGAGCCCTCCACCGGTCCGCCGTCC
TTCCGCGTGGGGGGGTCGTTAGGCGCGCGGCCCTGTCTTACGGGACGTCCTTGAAGAAGACCTTCTGGAAGAGGAGGAC
GTTTATTTTGGAGTGGGTACTTACGAGTGCGTTCAAATTAATGTCTGGACTT
```

#### 3.14 Program gto_fasta_reverse

The `gto_fasta_reverse` reverses the ACGT bases order for each read in a FASTA or Multi-FASTA file format.
For help type:

```
./gto_fasta_reverse -h
```

In the following subsections, we explain the input and output paramters.

##### Input parameters

The `gto_fasta_reverse` program needs two streams for the computation, namely the input and output standard. The input stream is a FASTA or Multi-FASTA file.
The attribution is given according to:

```
Usage: ./gto_fasta_reverse [options] [[--] args]
   or: ./gto_fasta_reverse [options]

It reverses the ACGT bases order for each read in a FASTA or Multi-FASTA file.

    -h, --help            Show this help message and exit

Basic options
    < input.fasta         Input FASTA or Multi-FASTA file format (stdin)
    > output.fasta        Output FASTA or Multi-FASTA file format (stdout)

Example: ./gto_fasta_reverse < input.mfasta > output.mfasta
```

An example of such an input file is:

```
>AB000264 |acc=AB000264|descr=Homo sapiens mRNA
ACAAGACGGCCTCCTGCTGCTGCTGCTCTCCGGGGCCACGGCCCTGGAGGGTCCACCGCTGCCCTGCTGCCATTGTCCC
CGGCCCCACCTAAGGAAAAGCAGCCTCCTGACTTTCCTCGCTTGGGCCGAGACAGCGAGCATATGCAGGAAGCGGCAGG
AAGTGGTTTGAGTGGACCTCCGGGCCCCTCATAGGAGAGGAAGCTCGGGAGGTGGCCAGGCGGCAGGAAGCAGGCCAGT
GCCGCGAATCCGCGCGCCGGGACAGAATCTCCTGCAAAGCCCTGCAGGAACTTCTTCTGGAAGACCTTCTCCACCCCCC
CAGCTAAAACCTCACCCATGAATGCTCACGCAAGTTTAATTACAGACCTGAA
>AB000263 |acc=AB000263|descr=Homo sapiens mRNA
ACAAGATGCCATTGTCCCCCGGCCTCCTGCTGCTGCTGCTCTCCGGGGCCACGGCCACCGCTGCCCTGCCCCTGGAGGG
TGGCCCCACCGGCCGAGACAGCGAGCATATGCAGGAAGCGGCAGGAATAAGGAAAAGCAGCCTCCTGACTTTCCTCGCT
TGGTGGTTTGAGTGGACCTCCCAGGCCAGTGCCGGGCCCCTCATAGGAGAGGAAGCTCGGGAGGTGGCCAGGCGGCAGG
```

```
AAGGCGCACCCCCCCAGCAATCCGCGCGCCGGGACAGAATGCCCTGCAGGAACTTCTTCTGGAAGACCTTCTCCTCCTG
CAAATAAAACCTCACCCATGAATGCTCACGCAAGTTTAATTACAGACCTGAA
```

##### Output

The output of the `gto_fasta_reverse` program is FASTA or Multi-FASTA file with the bases reversed and the flag "(Reversed)" added in the header.
Using the input above, an output example for this is the following:

```
>AB000264 |acc=AB000264|descr=Homo sapiens mRNA  (Reversed)
AAGTCCAGACATTAATTTGAACGCACTCGTAAGTACCCACTCCAAAATCGACCCCCCCACCTCTTCCAGAAGGTCTTCT
TCAAGGACGTCCCGAAACGTCCTCTAAGACAGGGCCGCGCGCCTAAGCGCCGTGACCGGACGAAGGACGGCGGACCGGT
GGAGGGCTCGAAGGAGAGGATACTCCCCGGGCCTCCAGGTGAGTTTGGTGAAGGACGGCGAAGGACGTATACGAGCGAC
AGAGCCGGGTTCGCTCCTTTCAGTCCTCCGACGAAAAGGAATCCACCCCGGCCCCTGTTACCGTCGTCCCGTCGCCACC
TGGGAGGTCCCGGCACCGGGGCCTCTCGTCGTCGTCGTCCTCCGGCAGAACA
>AB000263 |acc=AB000263|descr=Homo sapiens mRNA  (Reversed)
AAAGTCCAGACATTAATTTGAACGCACTCGTAAGTACCCACTCCAAAATAAACGTCCTCCTCTTCCAGAAGGTCTTCTT
CAAGGACGTCCCGTAAGACAGGGCCGCGCGCCTAACGACCCCCCCACGCGGAAGGACGGCGGACCGGTGGAGGGCTCGA
AGGAGAGGATACTCCCCGGGCCGTGACCGGACCCTCCAGGTGAGTTTGGTGGTTCGCTCCTTTCAGTCCTCCGACGAAA
AGGAATAAGGACGGCGAAGGACGTATACGAGCGACAGAGCCGGCCACCCCGGTGGGAGGTCCCCGTCCCGTCGCCACCG
GCACCGGGGCCTCTCGTCGTCGTCGTCCTCCGGCCCCCTGTTACCGTAGAACA
```

### Chapter 4

### Genomic sequence tools

The Genomic Sequence subset works directly with the DNA sequences, without any standard format. These tools allow the data extraction, summarising and some mathematical operations over those files. Usually, these are used in the pipeline as a complementary tool. The current available genomic sequence tools, for analysis and manipulation, are:

1. `gto_genomic_gen_random_dna`: it generates a synthetic DNA.
2. `gto_genomic_rand_seq_extra_chars`: it substitues in the DNA sequence the outside ACGT chars by random ACGT symbols.
3. `gto_genomic_dna_mutate`: it creates a synthetic mutation of a sequence file given specific rates of mutations, deletions and additions.
4. `gto_genomic_extract`: it extracts sequences from a sequence file, which the range is defined by the user in the parameters.
5. `gto_genomic_period`: it calculates the best order depth of a sequence, using FCMs.
6. `gto_genomic_count_bases`: it counts the number of bases in sequence, FASTA or FASTQ files.
7. `gto_genomic_compressor`: it compress and decompress genomic sequences for storage purposes (also under the alias gto_geco).
8. `gto_genomic_complement`: it replaces the ACGT bases with their complements in a DNA sequence.
9. `gto_genomic_reverse`: it reverses the ACGT bases order for each read in a sequence file (also under the alias gto_reverse).
10. `gto_genomic_variation_map`: this tool is an alias to gto_fastq_variation_map tool. Please check the documentation of this tool in the in the section of FASTQ tools.
11. `gto_genomic_variation_filter`: this tool is an alias to gto_fastq_variation_filter tool. Please check the documentation of this tool in the in the section of FASTQ tools.

12. `gto_genomic_variation_visual`: this tool is an alias to gto_fastq_variation_visual tool. Please check the documentation of this tool in the in the section of FASTQ tools.

#### 4.1 Program gto_genomic_gen_random_dna

The `gto_genomic_gen_random_dna` generates a synthetic DNA.
For help type:

```
./gto_genomic_gen_random_dna -h
```

In the following subsections, we explain the input and output paramters.

##### Input parameters

The `gto_genomic_gen_random_dna` program needs one stream for the computation, namely the output standard.
The attribution is given according to:

```
Usage: ./gto_genomic_gen_random_dna [options] [[--] args]
   or: ./gto_genomic_gen_random_dna [options]

It generates a synthetic DNA.

    -h, --help                show this help message and exit

Basic options
    > output.seq              Output synthetic DNA sequence (stdout)

Optional
    -s, --seed=<int>          Starting point to the random generator (Default 0)
    -n, --nSymbols=<int>      Number of symbols generated (Default 100)
    -f, --frequency=<str>     The frequency of each base. It should be represented
                              in the following format: <fa,fc,fg,ft>.

Example: ./gto_genomic_gen_random_dna -s <seed> -n <nsyboml s> -f <fa,fc,fg,ft> > output.seq
```

##### Output

The output of the `gto_genomic_gen_random_dna` program is a sequence group file whith the synthetic DNA.
Using the input above with the seed value as 1 and the number of symbols as 400, an output example for this is the following:

```
TCTTTACTCGCGCGTTGGAGAAATACAATAGTGCGGCTCTGTCTCCTTATGAAGTCAACAATTTCGCTGGGACTTGCGGC
TCTTTACTCGCGCGTTGGAGAAATACAATAGTGCGGCTCTGTCTCCTTATGAAGTCAACAATTTCGCTGGGACTTGCGGC
GACTTCATCGTGGTCTCTGTCATTATGCGCTCCAACGCATAACTTTGCGCCAGAAGATAGATAGAATGGTGTAAGAAACT
GTAATATATATAATGAACTTCGGCGAGTCTGTGGAGTTTTTGTTGCATTAGAGAGCCAAGAGGTCGGACGTCCTCACGTA
GCCCGAGACGGGCAGGGCGATGGCGACTGAACGGGCTCCATATCACTTTGAGCTTTTATGCTTTCGACTCCTCCAGGAGC
```

```
TGAACAACCTTGTTCCCGGCAAAGCCCACTGCGTCATGGAGCTCACGGTCTACATTCATGACTGACTAACCGTAAACTGC
```

#### 4.2 Program gto_genomic_rand_seq_extra_chars

The `gto_genomic_rand_seq_extra_chars` substitues in the DNA sequence the outside ACGT chars by random ACGT symbols. It works in sequence file formats.
For help type:

```
./gto_genomic_rand_seq_extra_chars -h
```

In the following subsections, we explain the input and output paramters.

##### Input parameters

The `gto_genomic_rand_seq_extra_chars` program needs two streams for the computation, namely the input and output standard. The input stream is a sequence file.
The attribution is given according to:

```
Usage: ./gto_genomic_rand_seq_extra_chars [options] [[--] args]
   or: ./gto_genomic_rand_seq_extra_chars [options]

It substitues in the DNA sequence the outside ACGT chars by random ACGT symbols.
It works in sequence file formats

    -h, --help        show this help message and exit

Basic options
    < input.seq       Input sequence file (stdin)
    > output.seq      Output sequence file (stdout)

Example: ./gto_genomic_rand_seq_extra_chars < input.seq > output.seq
```

An example of such an input file is:

```
ANAAGACGNNNTCCTGCTGCTGCTGCTCTCCGGGGCCACGGCCCTGGAGGGTCCACCGCTGCCCTGCTGCCATTGTCCCC
NNCCCCACCTAAGGAAAAGCAGCCTCCTGACTTTCCTCGCTTGGGCCGAGACAGCGAGCATATGCAGGAAGCGGCAGGAA
GTGGTTTGAGTGGACCTCCGGGCCCCNNNNNGGAGAGGAAGCTCGGGAGNGTNNNGGCCAGGCGGCAGNNNNCCAGTGCC
GCGAATCCGCGCGCCGGGACAGAATCTCCTGCAAAGCCCTGCAGGAACTTCTTCTGGAAGACCTTCTCCACCCCCCCAGC
TANNNNCTCACCCATGAATGCTCACGCAAGTTTAATTACAGACCTGAAACAAGATGCCATTGTCCCCCGGCCTCCTGCTG
CTGCTGCTCTCCGGGGCCACGGCCACCGCTGCCCTGCCCCTGGAGGGTGGCCCCACCGGCCGAGACAGCGAGCATATGCA
GGAAGCGGCAGGAATAAGNNNAAGCAGCCTCCTGACTTTCCTCGCTTGNNNNTTTGAGTGGACCTCCCAGGCCAGTGCCG
GGCCCCTCATAGGAGAGGAAGCTCGGGAGGTGGCCAGGCGGCAGGAAGGCGCACCCCCCCAGCAATCCGCGCGCCGGGAC
AGAATGCCCTGCAGGAACTTCTTCTGGAAGACCTTCTCCTCCTGCAAATAAAACCTCACCCATGAATGCTCACGCAAGTT
NNATTACNNNCCTGNN
```

##### Output

The output of the `gto_genomic_rand_seq_extra_chars` program is a sequence file.
Using the input above, an output example for this is the following:

```
ATAAGACGGCTTCCTGCTGCTGCTGCTCTCCGGGGCCACGGCCCTGGAGGGTCCACCGCTGCCCTGCTGCCATTGTCCCC
CTCCCCACCTAAGGAAAAGCAGCCTCCTGACTTTCCTCGCTTGGGCCGAGACAGCGAGCATATGCAGGAAGCGGCAGGAA
GTGGTTTGAGTGGACCTCCGGGCCCCGACCGGGAGAGGAAGCTCGGGAGTGTGTTGGCCAGGCGGCAGGAGACCAGTGCC
GCGAATCCGCGCGCCGGGACAGAATCTCCTGCAAAGCCCTGCAGGAACTTCTTCTGGAAGACCTTCTCCACCCCCCCAGC
TAATATCTCACCCATGAATGCTCACGCAAGTTTAATTACAGACCTGAAACAAGATGCCATTGTCCCCCGGCCTCCTGCTG
CTGCTGCTCTCCGGGGCCACGGCCACCGCTGCCCTGCCCCTGGAGGGTGGCCCCACCGGCCGAGACAGCGAGCATATGCA
GGAAGCGGCAGGAATAAGCGGAAGCAGCCTCCTGACTTTCCTCGCTTGGTTTTTTGAGTGGACCTCCCAGGCCAGTGCCG
GGCCCCTCATAGGAGAGGAAGCTCGGGAGGTGGCCAGGCGGCAGGAAGGCGCACCCCCCCAGCAATCCGCGCGCCGGGAC
AGAATGCCCTGCAGGAACTTCTTCTGGAAGACCTTCTCCTCCTGCAAATAAAACCTCACCCATGAATGCTCACGCAAGTT
CGATTACGGCCCTGTC
```

#### 4.3 Program gto_genomic_dna_mutate

The `gto_genomic_dna_mutate` creates a synthetic mutation of a sequence file given specific rates of mutations, deletions and additions. All these paramenters are defined by the user, and their are optional.
For help type:

```
./gto_genomic_dna_mutate -h
```

In the following subsections, we explain the input and output paramters.

##### Input parameters

The `gto_genomic_dna_mutate` program needs two streams for the computation, namely the input and output standard. However, optional settings can be supplied too, such as the starting point to the random generator, and the edition, deletion and insertion rates. Also, the user can choose to use the ACGTN alphabet in the synthetic mutation. The input stream is a sequence File.
The attribution is given according to:

```
Usage: ./gto_genomic_dna_mutate [options] [[--] args]
   or: ./gto_genomic_dna_mutate [options]

Creates a synthetic mutation of a sequence file given specific rates of mutations,
deletions and additions

    -h, --help                    show this help message and exit

Basic options
    < input.seq                   Input sequence file (stdin)
    > output.seq                  Output sequence file (stdout)

Optional
    -s, --seed=<int>              Starting point to the random generator
    -m, --mutation-rate=<dbl>     Defines the mutation rate (default 0.0)
    -d, --deletion-rate=<dbl>     Defines the deletion rate (default 0.0)
    -i, --insertion-rate=<dbl>    Defines the insertion rate (default 0.0)
    -a, --ACGTN-alphabet          When active, the application uses the ACGTN alphabet
```

```
Example: ./gto_genomic_dna_mutate -s <seed> -m <mutation rate> -d <deletion rate> -i
<insertion rate> -a < input.seq > output.seq
```

An example of such an input file is:

```
TCTTTACTCGCGCGTTGGAGAAATACAATAGTGCGGCTCTGTCTCCTTATGAAGTCAACAATTTCGCTGGGACTTGCGGC
TCTTTACTCGCGCGTTGGAGAAATACAATAGTGCGGCTCTGTCTCCTTATGAAGTCAACAATTTCGCTGGGACTTGCGGC
GACTTCATCGTGGTCTCTGTCATTATGCGCTCCAACGCATAACTTTGCGCCAGAAGATAGATAGAATGGTGTAAGAAACT
GTAATATATATAATGAACTTCGGCGAGTCTGTGGAGTTTTTGTTGCATTAGAGAGCCAAGAGGTCGGACGTCCTCACGTA
GCCCGAGACGGGCAGGGCGATGGCGACTGAACGGGCTCCATATCACTTTGAGCTTTTATGCTTTCGACTCCTCCAGGAGC
TGAACAACCTTGTTCCCGGCAAAGCCCACTGCGTCATGGAGCTCACGGTCTACATTCATGACTGACTAACCGTAAACTGC
```

##### Output

The output of the `gto_genomic_dna_mutate` program is a sequence file whith the synthetic mutation of input file.
Using the input above with the seed value as 1 and the mutation rate as 0.5, an output example for this is the following:

```
TCACGACTGTCGCGTTGGCACACCAGATAGGTGCTTCTACGTTTTGTATCTAATTTACAATTCTCGCTGGGAGTTCATTC
GCTATTGATGGGACTAGAAACCCATCCGTAGCTTGCCGCCGTTTAAGAATAAACACTCCACTTGCACCGAGACGTAGCGC
AACCAAGGCTATGTTCTTTGACCTTATGCGGTCCAACGCAGGAGTAGACCCCCGTAGTTAGGTACTATCGCAGAATAGGC
TTAAGCAGCCGTGCTGAACGCTGGAGGGTCTGTTTAATTACTGAGTGAATGGAGAGCTAAGAGTTCGGAGCACCGCACGA
GGCTCAAGAGCGGAAGGGCGTCAGCCTGGCGACCACCTGCCTACCGCTCGAGTCTGTCTTCACTACAGTCCGTGGAGGAC
CCCCAACGACCTAGTATCCTACAAAGCCGCATACGACTTACAGAACAGGCTGTATCGTCAGGAGTGTGTACACGAAGAGT
A
```

#### 4.4 Program gto_genomic_extract

The `gto_genomic_extract` extracts sequences from a sequence file, which the range is defined by the user in the parameters.
For help type:

```
./gto_genomic_extract -h
```

In the following subsections, we explain the input and output paramters.

##### Input parameters

The `gto_genomic_extract` program needs two paramenters, which defines the begin and the end of the extraction, and two streams for the computation, namely the input and output standard. The input stream is a sequence file.
The attribution is given according to:

```
Usage: ./gto_genomic_extract [options] [[--] args]
   or: ./gto_genomic_extract [options]

It extracts sequences from a sequence file.

    -h, --help        show this help message and exit

Basic options
    -i, --init=<int>  The first position to start the extraction (default 0)
    -e, --end=<int>   The last extract position (default 100)
    < input.seq       Input sequence file (stdin)
    > output.seq      Output sequence file (stdout)

Example: ./gto_genomic_extract -i <init> -e <end> < input.seq > output.seq
```

An example of such an input file is:

```
TCTTTACTCGCGCGTTGGAGAAATACAATAGTGCGGCTCTGTCTCCTTATGAAGTCAACAATTTCGCTGGGACTTGCGGC
TCTTTACTCGCGCGTTGGAGAAATACAATAGTGCGGCTCTGTCTCCTTATGAAGTCAACAATTTCGCTGGGACTTGCGGC
GACTTCATCGTGGTCTCTGTCATTATGCGCTCCAACGCATAACTTTGCGCCAGAAGATAGATAGAATGGTGTAAGAAACT
GTAATATATATAATGAACTTCGGCGAGTCTGTGGAGTTTTTGTTGCATTAGAGAGCCAAGAGGTCGGACGTCCTCACGTA
GCCCGAGACGGGCAGGGCGATGGCGACTGAACGGGCTCCATATCACTTTGAGCTTTTATGCTTTCGACTCCTCCAGGAGC
TGAACAACCTTGTTCCCGGCAAAGCCCACTGCGTCATGGAGCTCACGGTCTACATTCATGACTGACTAACCGTAAACTGC
```

##### Output

The output of the `gto_genomic_extract` program is a group sequence.
Using the input above with the value 0 as the extraction starting point and the 50 as the ending, an output example for this is the following:

```
TCTTTACTCGCGCGTTGGAGAAATACAATAGTGCGGCTCTGTCTCCTTAT
```

#### 4.5 Program gto_genomic_period

The `gto_genomic_period` calculates the best order depth of a sequence, using FCMs. It only works "ACGT", while the rest will be discarded.
This application has a dependency to represent the results. It requires the Gnuplot to show the execution result.
For help type:

```
./gto_genomic_period -h
```

In the following subsections, we explain the input and output paramters.

##### Input parameters

The `gto_genomic_period` program needs two streams for the computation, namely the input and output standard. The input stream is a sequence file.
The attribution is given according to:

```
Usage: ./gto_genomic_period [options] [[--] args]
   or: ./gto_genomic_period [options]

It calculates the best order depth of a sequence, using FCMs.It only works "ACGT",
while the rest will be discarded.

    -h, --help        show this help message and exit

Basic options
    < input.seq       Input sequence file format (stdin)
    > output          Output is given by log_2(4)*K(x)/|x|) (stdout)

Example: ./gto_genomic_period < input.seq > output
```

An example of such an input file is:

```
TCTTTACTCGCGCGTTGGAGAAATACAATAGTGCGGCTCTGTCTCCTTATGAAGTCAACAATTTCGCTGGGACTTGCGGC
TCTTTACTCGCGCGTTGGAGAAATACAATAGTGCGGCTCTGTCTCCTTATGAAGTCAACAATTTCGCTGGGACTTGCGGC
GACTTCATCGTGGTCTCTGTCATTATGCGCTCCAACGCATAACTTTGCGCCAGAAGATAGATAGAATGGTGTAAGAAACT
GTAATATATATAATGAACTTCGGCGAGTCTGTGGAGTTTTTGTTGCATTAGAGAGCCAAGAGGTCGGACGTCCTCACGTA
GCCCGAGACGGGCAGGGCGATGGCGACTGAACGGGCTCCATATCACTTTGAGCTTTTATGCTTTCGACTCCTCCAGGAGC
TGAACAACCTTGTTCCCGGCAAAGCCCACTGCGTCATGGAGCTCACGGTCTACATTCATGACTGACTAACCGTAAACTGC
```

##### Output

The output of the `gto_genomic_period` program is a execution report, followed by the plot with this information.
Using the input above, an report example for this is the following:

```
Running order: 1 ... Done!
Running order: 2 ... Done!
Running order: 3 ... Done!
Running order: 4 ... Done!
Running order: 5 ... Done!
Running order: 6 ... Done!
Running order: 7 ... Done!
Running order: 8 ... Done!
Running order: 9 ... Done!
Running order: 10 ... Done!
Running order: 11 ... Done!
Running order: 12 ... Done!
Running order: 13 ... Done!
Running order: 14 ... Done!
Running order: 15 ... Done!
Running order: 16 ... Done!
```

```
Running order: 17 ... Done!
Running order: 18 ... Done!
Running order: 19 ... Done!
Running order: 20 ... Done!
 1  2.246
 2  2.225
 3  2.237
 4  2.079
 5  1.821
 6  1.733
 7  1.717
 8  1.708
 9  1.717
10  1.712
11  1.717
12  1.721
13  1.725
14  1.729
15  1.733
16  1.738
17  1.742
18  1.746
19  1.75
20  1.754
```

In the Figure 4.1 is represented the plot for the execution above.

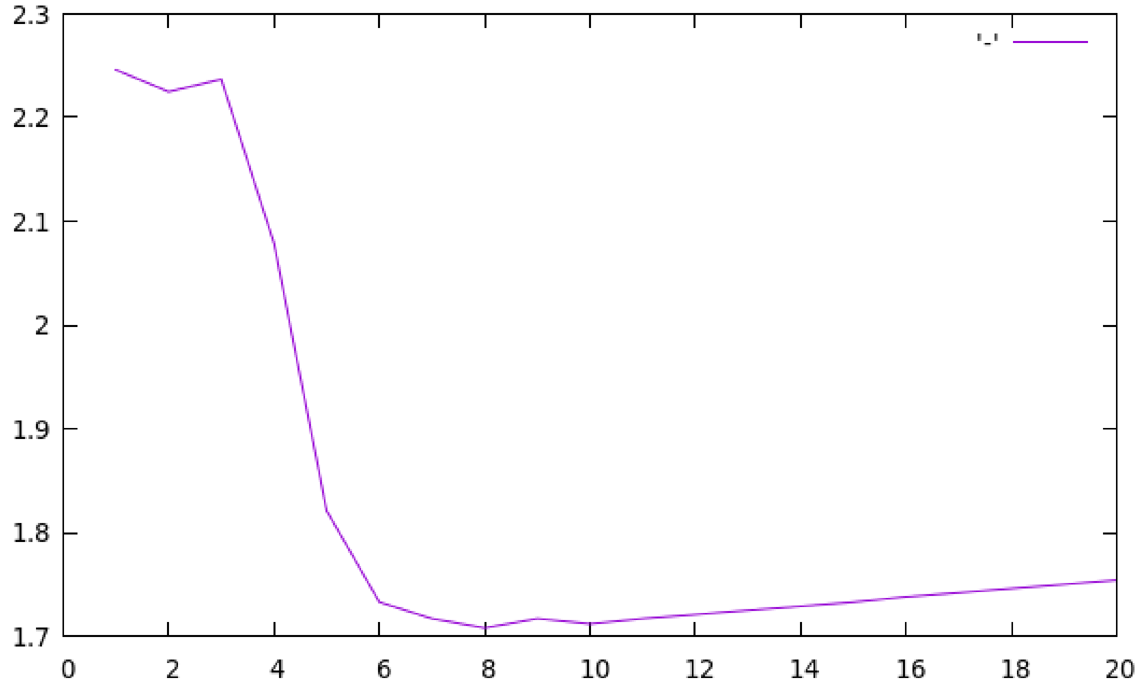

Figure 4.1: `gto_genomic_period` execution plot.

#### 4.6 Program gto_genomic_count_bases

The `gto_genomic_count_bases` counts the number of bases in sequence, FASTA or FASTQ files.
For help type:

```
./gto_genomic_count_bases -h
```

In the following subsections, we explain the input and output paramters.

##### Input parameters

The `gto_genomic_count_bases` program needs two streams for the computation, namely the input and output standard. The input stream is a sequence, FASTA or FASTQ file.
The attribution is given according to:

```
Usage: ./gto_genomic_count_bases [options] [[--] args]
   or: ./gto_genomic_count_bases [options]

It counts the number of bases in sequence, FASTA or FASTQ files.

    -h, --help      Show this help message and exit

Basic options
    < input         Input sequence, FASTA or FASTQ file format (stdin)
    > output        Output read information (stdout)

Example: ./gto_genomic_count_bases < input.seq > output

Output example :
File type        : value
Number of bases  : value
Number of a/A    : value
Number of c/C    : value
Number of g/G    : value
Number of t/T    : value
Number of n/N    : value
Number of others : value
```

An example of such an input file is:

```
TCTTTACTCGCGCGTTGGAGAAATACAATAGTGCGGCTCTGTCTCCTTATGAAGTCAACAATTTCGCTGGGACTTGCGGC
TCTTTACTCGCGCGTTGGAGAAATACAATAGTGCGGCTCTGTCTCCTTATGAAGTCAACAATTTCGCTGGGACTTGCGGC
GACTTCATCGTGGTCTCTGTCATTATGCGCTCCAACGCATAACTTTGCGCCAGAAGATAGATAGAATGGTGTAAGAAACT
GTAATATATATAATGAACTTCGGCGAGTCTGTGGAGTTTTTGTTGCATTAGAGAGCCAAGAGGTCGGACGTCCTCACGTA
GCCCGAGACGGGCAGGGCGATGGCGACTGAACGGGCTCCATATCACTTTGAGCTTTTATGCTTTCGACTCCTCCAGGAGC
TGAACAACCTTGTTCCCGGCAAAGCCCACTGCGTCATGGAGCTCACGGTCTACATTCATGACTGACTAACCGTAAACTGC
```

##### Output

The output of the `gto_genomic_count_bases` program is report which describes the number of each base in the file, and the file type.

Using the input above, an output example for this is the following:

```
File type        : DNA
Number of bases  : 480
Number of a/A    : 114
Number of c/C    : 116
Number of g/G    : 120
Number of t/T    : 130
Number of n/N    : 0
Number of others : 0
```

#### 4.7 Program gto_genomic_compressor

The `gto_genomic_compressor` is able to provide additional compression gains over several top specific tools, while as an analysis tool, it is able to determine absolute measures, namely for many distance computations, and local measures, such as the information content contained in each element, providing a way to quantify and locate specific genomic events.
For help type:

```
./gto_genomic_compressor -h
```

In the following subsections, we explain the input and output paramters.

##### Input parameters

The `gto_genomic_compressor` program needs a sequence to compress.
The attribution is given according to:

```
SYNOPSIS
      ./gto_genomic_compressor [OPTION]... -r [FILE] [FILE]:[FILE]:[FILE]:[...]

SAMPLE
      Run Compression         :  ./gto_genomic_compressor -v -l 3 sequence.txt
      Run Decompression       :  ./gto_genomic_decompressor -v sequence.txt.co
      Run Information Profile :  ./gto_genomic_compressor -v -l 3 -e sequence.txt

DESCRIPTION
      Compress and decompress genomic sequences for storage purposes.
      Measure an upper bound of the sequences entropy.
      Compute information profiles of genomic sequences.

      -h,  --help
           usage guide (help menu).

      -V,  --version
           Display program and version information.

      -F,  --force
           force mode. Overwrites old files.
```

```
-v,  --verbose
     verbose mode (more information).

-x,  --examples
     show several running examples (parameter examples).

-s,  --show-levels
     show pre-computed compression levels (configured parameters).

-e,  --estimate
     it creates a file with the extension ".iae" with the
     respective information content. If the file is FASTA or
     FASTQ it will only use the "ACGT" (genomic) sequence.

-l [NUMBER],  --level [NUMBER]
     Compression level (integer).
     Default level: 5.
     It defines compressibility in balance with computational
     resources (RAM & time). Use -s for levels perception.

-tm [NB_C]:[NB_D]:[NB_I]:[NB_H]:[NB_G]/[NB_S]:[NB_E]:[NB_A]
     Template of a target context model.
     Parameters:
     [NB_C]: (integer [1;20]) order size of the regular context
             model. Higher values use more RAM but, usually, are
             related to a better compression score.
     [NB_D]: (integer [1;5000]) denominator to build alpha, which
             is a parameter estimator. Alpha is given by 1/[NB_D].
             Higher values are usually used with higher [NB_C],
             and related to confiant bets. When [NB_D] is one,
             the probabilities assume a Laplacian distribution.
     [NB_I]: (integer {0,1,2}) number to define if a sub-program
             which addresses the specific properties of DNA
             sequences (Inverted repeats) is used or not. The
             number 2 turns ON this sub-program without the
             regular context model (only inverted repeats). The
             number 1 turns ON the sub-program using at the same
             time the regular context model. The number 0 does
             not contemple its use (Inverted repeats OFF). The
             use of this sub-program increases the necessary time
             to compress but it does not affect the RAM.
     [NB_H]: (integer [1;254]) size of the cache-hash for deeper
             context models, namely for [NB_C] > 14. When the
             [NB_C] <= 14 use, for example, 1 as a default. The
             RAM is highly dependent of this value (higher value
             stand for higher RAM).
     [NB_G]: (real [0;1)) real number to define gamma. This value
             represents the decayment forgetting factor of the
             regular context model in definition.
     [NB_S]: (integer [0;20]) maximum number of editions allowed
             to use a substitutional tolerant model with the same
             memory model of the regular context model with
             order size equal to [NB_C]. The value 0 stands for
```

```
                turning the tolerant context model off. When the
                model is on, it pauses when the number of editions
                is higher that [NB_C], while it is turned on when
                a complete match of size [NB_C] is seen again. This
                is probabilistic-algorithmic model very usefull to
                handle the high substitutional nature of genomic
                sequences. When [NB_S] > 0, the compressor used more
                processing time, but uses the same RAM and, usually,
                achieves a substantial higher compression ratio. The
                impact of this model is usually only noticed for
                [NB_C] >= 14.
        [NB_E]: (integer [1;5000]) denominator to build alpha for
                substitutional tolerant context model. It is
                analogous to [NB_D], however to be only used in the
                probabilistic model for computing the statistics of
                the substitutional tolerant context model.
        [NB_A]: (real [0;1)) real number to define gamma. This value
                represents the decayment forgetting factor of the
                substitutional tolerant context model in definition.
                Its definition and use is analogus to [NB_G].

   ... (you may use several target models with custom parameters)

   -rm [NB_C]:[NB_D]:[NB_I]:[NB_H]:[NB_G]/[NB_S]:[NB_E]:[NB_A]
        Template of a reference context model.
        Use only when -r [FILE] is set (referential compression).
        Parameters: the same as in -tm.

   ... (you may use several reference models with custom parameters)

   -r [FILE], --reference [FILE]
        Reference sequence filename ("-rm" are trainned here).
        Example: -r file1.txt.

   [FILE]
        Input sequence filename (to compress) -- MANDATORY.
        File(s) to compress (last argument).
        For more files use splitting ":" characters.
        Example: file1.txt:file2.txt:file3.txt.
```

In the following example, it will be downloaded seventeen DNA sequences, and compress and decompress one of the smallest (BuEb). Finally, it compares if the uncompressed sequence is equal to the original.

```
wget http://sweet.ua.pt/pratas/datasets/DNACorpus.zip
unzip DNACorpus.zip
cp DNACorpus/BuEb .
../../bin/gto_genomic_compressor -v -l 2 BuEb
../../bin/gto_genomic_decompressor -v BuEb.co
cmp BuEb BuEb.de -l
```

#### 4.8 Program gto_genomic_complement

The `gto_genomic_complement` replaces the ACGT bases with their complements in a DNA sequence. It works in sequence file formats.
For help type:

```
./gto_genomic_complement -h
```

In the following subsections, we explain the input and output paramters.

##### Input parameters

The `gto_genomic_complement` program needs two paramenters, namely the input and output standard. The input stream is a sequence file.
The attribution is given according to:

```
Usage: ./gto_genomic_complement [options] [[--] args]
   or: ./gto_genomic_complement [options]

It replaces the ACGT bases with their complements in a DNA sequence.
It works in sequence file formats

    -h, --help          Show this help message and exit

Basic options
    < input.seq         Input sequence file (stdin)
    > output.seq        Output sequence file (stdout)

Example: ./gto_genomic_complement < input.seq > output.seq
```

An example of such an input file is:

```
TCTTTACTCGCGCGTTGGAGAAATACAATAGTGCGGCTCTGTCTCCTTATGAAGTCAACAATTTCGCTGGGACTTGCGG
CTCTTTACTCGCGCGTTGGAGAAATACAATAGTGCGGCTCTGTCTCCTTATGAAGTCAACAATTTCGCTGGGACTTGCG
GCGACTTCATCGTGGTCTCTGTCATTATGCGCTCCAACGCATAACTTTGCGCCAGAAGATAGATAGAATGGTGTAAGAA
ACTGTAATATATATAATGAACTTCGGCGAGTCTGTGGAGTTTTTGTTGCATTAGAGAGCCAAGAGGTCGGACGTCCTCA
CGTAGCCCGAGACGGGCAGGGCGATGGCGACTGAACGGGCTCCATATCACTTTGAGCTTTTATGCTTTCGACTCCTCCA
GGAGCTGAACAACCTTGTTCCCGGCAAAGCCCACTGCGTCATGGAGCTCACGGTCTACATTCATGACTGACTAACCGTA
AACTGC
```

##### Output

The output of the `gto_genomic_complement` program is a group sequence with the ACGT base complements.
Using the input above, an output example for this is the following:

```
AGAAATGAGCGCGCAACCTCTTTATGTTATCACGCCGAGACAGAGGAATACTTCAGTTGTTAAAGCGACCCTGAACGCC
GAGAAATGAGCGCGCAACCTCTTTATGTTATCACGCCGAGACAGAGGAATACTTCAGTTGTTAAAGCGACCCTGAACGC
CGCTGAAGTAGCACCAGAGACAGTAATACGCGAGGTTGCGTATTGAAACGCGGTCTTCTATCTATCTTACCACATTCTT
TGACATTATATATATTACTTGAAGCCGCTCAGACACCTCAAAAACAACGTAATCTCTCGGTTCTCCAGCCTGCAGGAGT
GCATCGGGCTCTGCCCGTCCCGCTACCGCTGACTTGCCCGAGGTATAGTGAAACTCGAAAATACGAAAGCTGAGGAGGT
CCTCGACTTGTTGGAACAAGGGCCGTTTCGGGTGACGCAGTACCTCGAGTGCCAGATGTAAGTACTGACTGATTGGCAT
TTGACG
```

#### 4.9 Program gto_genomic_reverse

The `gto_genomic_reverse` reverses the ACGT bases order for each read in a sequence file.
For help type:

```
./gto_genomic_reverse -h
```

In the following subsections, we explain the input and output paramters.

##### Input parameters

The `gto_genomic_reverse` program needs two streams for the computation, namely the input and output standard. The input stream is a sequence file.
The attribution is given according to:

```
Usage: ./gto_genomic_reverse [options] [[--] args]
   or: ./gto_genomic_reverse [options]

It reverses the ACGT bases order for each read in a sequence file.

    -h, --help        show this help message and exit

Basic options
    < input.seq       Input sequence file (stdin)
    > output.seq      Output sequence file (stdout)

Example: ./gto_genomic_reverse < input.seq > output.seq
```

An example of such an input file is:

```
ACAAGACGGCCTCCTGCTGCTGCTGCTCTCCGGGGCCACGGCCCTGGAGGGTCCACCGCTGCCCTGCTGCCATTGTCCCC
GGCCCCACCTAAGGAAAAGCAGCCTCCTGACTTTCCTCGCTTGGGCCGAGACAGCGAGCATATGCAGGAAGCGGCAGGAA
GTGGTTTGAGTGGACCTCCGGGCCCCTCATAGGAGAGGAAGCTCGGGAGGTGGCCAGGCGGCAGGAAGCAGGCCAGTGCC
GCGAATCCGCGCGCCGGGACAGAATCTCCTGCAAAGCCCTGCAGGAACTTCTTCTGGAAGACCTTCTCCACCCCCCCAGC
TAAAACCTCACCCATGAATGCTCACGCAAGTTTAATTACAGACCTGAAACAAGATGCCATTGTCCCCCGGCCTCCTGCTG
CTGCTGCTCTCCGGGGCCACGGCCACCGCTGCCCTGCCCCTGGAGGGTGGCCCCACCGGCCGAGACAGCGAGCATATGCA
GGAAGCGGCAGGAATAAGGAAAAGCAGCCTCCTGACTTTCCTCGCTTGGTGGTTTGAGTGGACCTCCCAGGCCAGTGCCG
GGCCCCTCATAGGAGAGGAAGCTCGGGAGGTGGCCAGGCGGCAGGAAGGCGCACCCCCCCAGCAATCCGCGCGCCGGGAC
AGAATGCCCTGCAGGAACTTCTTCTGGAAGACCTTCTCCTCCTGCAAATAAAACCTCACCCATGAATGCTCACGCAAGTT
TAATTACAGACCTGAA
```

#### Output

The output of the `gto_genomic_reverse` program is a group sequence.

Using the input above, an output example for this is the following:

```
AAGTCCAGACATTAATTTGAACGCACTCGTAAGTACCCACTCCAAAATAAACGTCCTCCTCTTCCAGAAGGTCTTCTTCA
AGGACGTCCCGTAAGACAGGGCCGCGCGCCTAACGACCCCCCCACGCGGAAGGACGGCGGACCGGTGGAGGGCTCGAAGG
AGAGGATACTCCCCGGGCCGTGACCGGACCCTCCAGGTGAGTTTGGTGGTTCGCTCCTTTCAGTCCTCCGACGAAAAGGA
ATAAGGACGGCGAAGGACGTATACGAGCGACAGAGCCGGCCACCCCGGTGGGAGGTCCCCGTCCCGTCGCCACCGGCACC
GGGGCCTCTCGTCGTCGTCGTCCTCCGGCCCCCTGTTACCGTAGAACAAAGTCCAGACATTAATTTGAACGCACTCGTAA
GTACCCACTCCAAAATCGACCCCCCCACCTCTTCCAGAAGGTCTTCTTCAAGGACGTCCCGAAACGTCCTCTAAGACAGG
GCCGCGCGCCTAAGCGCCGTGACCGGACGAAGGACGGCGGACCGGTGGAGGGCTCGAAGGAGAGGATACTCCCCGGGCCT
CCAGGTGAGTTTGGTGAAGGACGGCGAAGGACGTATACGAGCGACAGAGCCGGGTTCGCTCCTTTCAGTCCTCCGACGAA
AAGGAATCCACCCCGGCCCCTGTTACCGTCGTCCCGTCGCCACCTGGGAGGTCCCGGCACCGGGGCCTCTCGTCGTCGTC
GTCCTCCGGCAGAACA
```

### Chapter 5

### Amino acid sequence tools

A more specific subset of tools is the Amino Acid Sequence tools, designed to manipulate amino acid sequences. The main features of those tools are grouping sequences, for instance by their properties, such as electric charge (positive and negative), uncharged side chains, hydrophobic side chains and special cases. It is also possible generating pseudo-DNA with characteristics passed by amino acid sequences, or for data compression, using cooperation between multiple contexts and substitutional tolerant context models. The current available amino acid sequence tools, for analysis and manipulation, are:

1. `gto_amino_acid_to_group`: it converts an amino acid sequence to a group sequence.
2. `gto_amino_acid_to_pseudo_dna`: it converts an amino acid (protein) sequence to a pseudo DNA sequence.
3. `gto_amino_acid_compressor`: it is a new lossless compressor to compress efficiently amino acid sequences (proteins).
4. `gto_amino_acid_from_fasta`: it converts DNA sequences in FASTA or Multi-FASTA file format to an amino acid sequence.
5. `gto_amino_acid_from_fastq`: it converts DNA sequences in the FASTQ file format to an amino acid sequence.
6. `gto_amino_acid_from_seq`: it converts DNA sequences to an amino acid sequence.

#### 5.1 Program gto_amino_acid_to_group

The `gto_amino_acid_to_group` converts an amino acid sequence to a group sequence.
For help type:

```
./gto_amino_acid_to_group -h
```

In the following subsections, we explain the input and output paramters.

#### Input parameters

The `gto_amino_acid_to_group` program needs two streams for the computation, namely the input and output standard. The input stream is an amino acid sequence. The attribution is given according to:

```
Usage: ./gto_amino_acid_to_group [options] [[--] args]
   or: ./gto_amino_acid_to_group [options]

It converts a amino acid sequence to a group sequence.

    -h, --help             show this help message and exit

Basic options
    < input.prot           Input amino acid sequence file (stdin)
    > output.group         Output group sequence file (stdout)

Example: ./gto_amino_acid_to_group < input.prot > output.group
Table:
Prot    Group
R       P
H       P  Amino acids with electric charged side chains: POSITIVE
K       P
-       -
D       N
E       N  Amino acids with electric charged side chains: NEGATIVE
-       -
S       U
T       U
N       U  Amino acids with electric UNCHARGED side chains
Q       U
-       -
C       S
U       S
G       S  Special cases
P       S
-       -
A       H
V       H
I       H
L       H
M       H  Amino acids with hydrophobic side chains
F       H
Y       H
W       H
-       -
*       *  Others
X       X  Unknown
```

It can be used to group amino acids by properties, such as electric charge (positive and negative), uncharged side chains, hydrophobic side chains and special cases. An example of such an input file is:

```
IPFLLKKQFALADKLVLSKLRQLLGGRIKMMPCGGAKLEPAIGLFFHAIGINIKLGYGMTETTATVSCWHDFQFNPNSIG
TLMPKAEVKIGENNEILVRGGMVMKGYYKKPEETAQAFTEDGFLKTGDAGEFDEQGNLFITDRIKELMKTSNGKYIAPQY
```

```
IESKIGKDKFIEQIAIIADAKKYVSALIVPCFDSLEEYAKQLNIKYHDRLELLKNSDILKMFE
```

##### Output

The output of the `gto_amino_acid_to_group` program is a group sequence.
Using the input above, an output example for this is the following:

```
HSHHHPPUHHHHNPHHHUPHPUHHSSPHPHHSSSSHPHNSHHSHHHPHHSHUHPHSHSHUNUUHUHUSHPNHUHUSUUHS
UHHSPHNHPHSNUUNHHHPSSHHHPSHHPPSNNUHUHHUNNSHHPUSNHSNHNNUSUHHHUNPHPNHHPUUUSPHHHSUH
HNUPHSPNPHHNUHHHHHNHPPHHUHHHHSSHNUHNNHHPUHUHPHPNPHNHHPUUNHHPHHN
```

#### 5.2 Program gto_amino_acid_to_pseudo_dna

The `gto_amino_acid_to_pseudo_dna` converts an amino acid (protein) sequence to a pseudo DNA sequence.
For help type:

```
./gto_amino_acid_to_pseudo_dna -h
```

In the following subsections, we explain the input and output paramters.

##### Input parameters

The `gto_amino_acid_to_pseudo_dna` program needs two streams for the computation, namely the input and output standard. The input stream is an amino acid sequence. The attribution is given according to:

```
Usage: ./gto_amino_acid_to_pseudo_dna [options] [[--] args]
   or: ./gto_amino_acid_to_pseudo_dna [options]

It converts a protein sequence to a pseudo DNA sequence.

    -h, --help        show this help message and exit

Basic options
    < input.prot      Input amino acid sequence file (stdin)
    > output.dna      Output DNA sequence file (stdout)

Example: ./gto_amino_acid_to_pseudo_dna < input.prot > output.dna
Table:
Prot    DNA
A       GCA
C       TGC
D       GAC
E       GAG
F       TTT
G       GGC
H       CAT
I       ATC
```

```
K       AAA
L       CTG
M       ATG
N       AAC
P       CCG
Q       CAG
R       CGT
S       TCT
T       ACG
V       GTA
W       TGG
Y       TAC
*       TAG
X       GGG
```

It can be used to generate pseudo-DNA with characteristics passed by amino acid (protein) sequences. An example of such an input file is:

```
IPFLLKKQFALADKLVLSKLRQLLGGRIKMMPCGGAKLEPAIGLFFHAIGINIKLGYGMTETTATVSCWHDFQFNPNSIG
TLMPKAEVKIGENNEILVRGGMVMKGYYKKPEETAQAFTEDGFLKTGDAGEFDEQGNLFITDRIKELMKTSNGKYIAPQY
IESKIGKDKFIEQIAIIADAKKYVSALIVPCFDSLEEYAKQLNIKYHDRLELLKNSDILKMFE
```

##### Output

The output of the `gto_amino_acid_to_pseudo_dna` program is a DNA sequence.
Using the input above, an output example for this is the following:

```
ATCCCGTTTCTGCTGAAAAAACAGTTTGCACTGGCAGACAAACTGGTACTGTCTAAACTGCGTCAGCTGCTGGGCGGCCG
TATCAAAATGATGCCGTGCGGCGGCGCAAAACTGGAGCCGGCAATCGGCCTGTTTTTTCATGCAATCGGCATCAACATCA
AACTGGGCTACGGCATGACGGAGACGACGGCAACGGTATCTTGCTGGCATGACTTTCAGTTTAACCCGAACTCTATCGGC
ACGCTGATGCCGAAAGCAGAGGTAAAAATCGGCGAGAACAACGAGATCCTGGTACGTGGCGGCATGGTAATGAAAGGCTA
CTACAAAAAACCGGAGGAGACGGCACAGGCATTTACGGAGGACGGCTTTCTGAAAACGGGCGACGCAGGCGAGTTTGACG
AGCAGGGCAACCTGTTTATCACGGACCGTATCAAAGAGCTGATGAAAACGTCTAACGGCAAATACATCGCACCGCAGTAC
ATCGAGTCTAAAATCGGCAAAGACAAATTTATCGAGCAGATCGCAATCATCGCAGACGCAAAAAAATACGTATCTGCACT
GATCGTACCGTGCTTTGACTCTCTGGAGGAGTACGCAAAACAGCTGAACATCAAATACCATGACCGTCTGGAGCTGCTGA
AAAACTCTGACATCCTGAAAATGTTTGAG
```

#### 5.3 Program gto_amino_acid_compressor

The `gto_amino_acid_compressor` is a new lossless compressor to compress efficiently amino acid sequences (proteins). It uses a cooperation between multiple context and substitutional tolerant context models. The cooperation between models is balanced with weights that benefit the models with better performance according to a forgetting function specific for each model.
For help type:

```
./gto_amino_acid_compressor -h
```

In the following subsections, we explain the input and output paramters.

##### Input parameters

The `gto_amino_acid_compressor` program needs a file with amino acid sequences to compress. The attribution is given according to:

```
Usage: ./gto_amino_acid_compressor [OPTION]... -r [FILE]  [FILE]:[...]
Compression of amino acid sequences.

Non-mandatory arguments:

  -h                     give this help,
  -s                     show AC compression levels,
  -v                     verbose mode (more information),
  -V                     display version number,
  -f                     force overwrite of output,
  -l <level>             level of compression [1;7] (lazy -tm setup),
  -t <threshold>         threshold frequency to discard from alphabet,
  -e                     it creates a file with the extension ".iae"
                         with the respective information content.

  -rm <c>:<d>:<g>/<m>:<e>:<a>  reference model (-rm 1:10:0.9/0:0:0),
  -rm <c>:<d>:<g>/<m>:<e>:<a>  reference model (-rm 5:90:0.9/1:50:0.8),
  ...
  -tm <c>:<d>:<g>/<m>:<e>:<a>  target model (-tm 1:1:0.8/0:0:0),
  -tm <c>:<d>:<g>/<m>:<e>:<a>  target model (-tm 7:100:0.9/2:10:0.85),
  ...
                         target and reference templates use <c> for
                         context-order size, <d> for alpha (1/<d>), <g>
                         for gamma (decayment forgetting factor) [0;1),
                         <m> to the maximum sets the allowed mutations,
                         on the context without being discarded (for
                         deep contexts), under the estimator <e>, using
                         <a> for gamma (decayment forgetting factor)
                         [0;1) (tolerant model),

  -r <FILE>              reference file ("-rm" are loaded here),

Mandatory arguments:

  <FILE>:<...>:<...>     file to compress (last argument). For more
                         files use splitting ":" characters.

Example:

  [Compress]   ./gto_amino_acid_compressor -v -tm 1:1:0.8/0:0:0 -tm 5:20:0.9/3:20:0.9 seq.txt
  [Decompress] ./gto_amino_acid_decompressor -v seq.txt.co
```

In the following example, it will be downloaded nine amino acid sequences and compress and decompress one of the smallest (HI). Finally, it compares if the uncompressed sequence is equal to the original.

```
wget http://sweet.ua.pt/pratas/datasets/AminoAcidsCorpus.zip
unzip AminoAcidsCorpus.zip
cp AminoAcidsCorpus/HI .
```

```
./gto_amino_acid_compressor -v -l 2 HI
./gto_amino_acid_decompressor -v HI.co
cmp HI HI.de
```

#### 5.4 Program gto_amino_acid_from_fasta

The `gto_amino_acid_from_fasta` converts DNA sequences in FASTA or Multi-FASTA file format to an amino acid sequence.
For help type:

```
./gto_amino_acid_from_fasta -h
```

In the following subsections, we explain the input and output paramters.

##### Input parameters

The `gto_amino_acid_from_fasta` program needs two streams for the computation, namely the input and output standard. The input stream is a FASTA or Multi-FASTA file.
The attribution is given according to:

```
Usage: ../../bin/gto_amino_acid_from_fasta [options] [[--] args]
   or: ../../bin/gto_amino_acid_from_fasta [options]

It converts FASTA or Multi-FASTA file format to an amino acid sequence (translation).

    -h, --help              Show this help message and exit

Basic options
    < input.mfasta          Input FASTA or Multi-FASTA file format (stdin)
    > output.prot           Output amino acid sequence file (stdout)

Optional
    -f, --frame=<int>       Translation codon frame (1, 2 or 3)

Example: ../../bin/gto_amino_acid_from_fasta < input.mfasta > output.prot
```

An example of such an input file is:

```
>AB000264 |acc=AB000264|descr=Homo sapiens mRNA
ACAAGACGGCCTCCTGCTGCTGCTGCTCTCCGGGGCCACGGCCCTGGAGGGTCCACCGCTGCCCTGCTGCCATTGTCCC
CGGCCCCACCTAAGGAAAAGCAGCCTCCTGACTTTCCTCGCTTGGGCCGAGACAGCGAGCATATGCAGGAAGCGGCAGG
AAGTGGTTTGAGTGGACCTCCGGGCCCCTCATAGGAGAGGAAGCTCGGGAGGTGGCCAGGCGGCAGGAAGCAGGCCAGT
GCCGCGAATCCGCGCGCCGGGACAGAATCTCCTGCAAAGCCCTGCAGGAACTTCTTCTGGAAGACCTTCTCCACCCCCC
CAGCTAAAACCTCACCCATGAATGCTCACGCAAGTTTAATTACAGACCTGAA
>AB000263 |acc=AB000263|descr=Homo sapiens mRNA
ACAAGATGCCATTGTCCCCCGGCCTCCTGCTGCTGCTGCTCTCCGGGGCCACGGCCACCGCTGCCCTGCCCCTGGAGGG
TGGCCCCACCGGCCGAGACAGCGAGCATATGCAGGAAGCGGCAGGAATAAGGAAAAGCAGCCTCCTGACTTTCCTCGCT
TGGTGGTTTGAGTGGACCTCCCAGGCCAGTGCCGGGCCCCTCATAGGAGAGGAAGCTCGGGAGGTGGCCAGGCGGCAGG
```

```
AAGGCGCACCCCCCCCAGCAATCCGCGCGCCGGGACAGAATGCCCTGCAGGAACTTCTTCTGGAAGACCTTCTCCTCCTG
CAAATAAAACCTCACCCATGAATGCTCACGCAAGTTTAATTACAGACCTGAA
```

##### Output

The output of the `gto_amino_acid_from_fasta` program is an amino acid sequence.
Using the input above, an output example for this is the following:

```
TRRPPAAAALRGHGPGGSTAALLPLSPAPPKEKQPPDFPRLGRDSEHMQEAAGSGLSGPPGPS-ERKLGRWPGGRKQAS
AANPRAGTESPAKPCRNFFWKTFSTPPAKTSPMNAHASLITDLTRCHCPPASCCCCSPGPRPPLPCPWRVAPPAETASI
CRKRQE-GKAAS-LSSLGGLSGPPRPVPGPS-ERKLGRWPGGRKAHPPSNPRAGTECPAGTSSGRPSPPANKTSPMNAH
ASLITDL
```

#### 5.5 Program gto_amino_acid_from_fastq

The `gto_amino_acid_from_fastq` converts DNA sequences in the FASTQ file format to an amino acid sequence.
For help type:

```
./gto_amino_acid_from_fastq -h
```

In the following subsections, we explain the input and output paramters.

##### Input parameters

The `gto_amino_acid_from_fastq` program needs two streams for the computation, namely the input and output standard. The input stream is a FASTQ file.
The attribution is given according to:

```
Usage: ../../bin/gto_amino_acid_from_fastq [options] [[--] args]
   or: ../../bin/gto_amino_acid_from_fastq [options]

It converts FASTQ file format to an amino acid sequence (translation).

    -h, --help            Show this help message and exit

Basic options
    < input.fastq         Input FASTQ file format (stdin)
    > output.prot         Output amino acid sequence file (stdout)

Optional
    -f, --frame=<int>     Translation codon frame (1, 2 or 3)

Example: ../../bin/gto_amino_acid_from_fastq < input.fastq > output.prot
```

An example of such an input file is:

```
@SRR001666.1 071112_SLXA-EAS1_s_7:5:1:817:345 length=72
GGGTGATGGCCGCTGCCGATGGCGTCAAATCCCACCAAGTTACCCTTAACAACTTAAGGGTTTTCAAATAGA
+SRR001666.1 071112_SLXA-EAS1_s_7:5:1:817:345 length=72
IIIIIIIIIIIIIIIIIIIIIIIIIIIIII9IG9ICIIIIIIIIIIIIIIIIIIIIDIIIIIII>IIIIII/
@SRR001666.2 071112_SLXA-EAS1_s_7:5:1:801:338 length=72
GTTCAGGGATACGACGTTTGTATTTTAAGAATCTGAAGCAGAAGTCGATGATAATACGCGTCGTTTTATCAT
+SRR001666.2 071112_SLXA-EAS1_s_7:5:1:801:338 length=72
IIIIIIIIIIIIIIIIIIIIIIIIIIIIIIII6IBIIIIIIIIIIIIIIIIIIIIIIIGII>IIIII-I)8I
```

##### Output

The output of the `gto_amino_acid_from_fastq` program is an amino acid sequence.
Using the input above, an output example for this is the following:

```
G-WPLPMASNPTKLPLTT-GFSNRVQGYDVCILRI-SRSR--YASFYH
```

#### 5.6 Program gto_amino_acid_from_seq

The `gto_amino_acid_from_seq` converts DNA sequence to an amino acid sequence.
For help type:

```
./gto_amino_acid_from_seq -h
```

In the following subsections, we explain the input and output paramters.

##### Input parameters

The `gto_amino_acid_from_seq` program needs two streams for the computation, namely the input and output standard. The input stream is a DNA sequence.
The attribution is given according to:

```
Usage: ../../bin/gto_amino_acid_from_seq [options] [[--] args]
   or: ../../bin/gto_amino_acid_from_seq [options]

It converts DNA sequence to an amino acid sequence (translation).

    -h, --help            Show this help message and exit

Basic options
    < input.seq           Input sequence file (stdin)
    > output.prot         Output amino acid sequence file (stdout)

Optional
    -f, --frame=<int>     Translation codon frame (1, 2 or 3)

Example: ../../bin/gto_amino_acid_from_seq < input.seq > output.prot
```

An example of such an input file is:

```
ACAAGACGGCCTCCTGCTGCTGCTGCTCTCCGGGGCCACGGCCCTGGAGGGTCCACCGCTGCCCTGCTGCCATTGTCCC
CGGCCCCACCTAAGGAAAAGCAGCCTCCTGACTTTCCTCGCTTGGGCCGAGACAGCGAGCATATGCAGGAAGCGGCAGG
AAGTGGTTTGAGTGGACCTCCGGGCCCCTCATAGGAGAGGAAGCTCGGGAGGTGGCCAGGCGGCAGGAAGCAGGCCAGT
GCCGCGAATCCGCGCGCCGGGACAGAATCTCCTGCAAAGCCCTGCAGGAACTTCTTCTGGAAGACCTTCTCCACCCCCC
CAGCTAAAACCTCACCCATGAATGCTCACGCAAGTTTAATTACAGACCTGAAACAAGATGCCATTGTCCCCCGGCCTCC
TGCTGCTGCTGCTCTCCGGGGCCACGGCCACCGCTGCCCTGCCCCTGGAGGGTGGCCCCACCGGCCGAGACAGCGAGCA
TATGCAGGAAGCGGCAGGAATAAGGAAAAGCAGCCTCCTGACTTTCCTCGCTTGGTGGTTTGAGTGGACCTCCCAGGCC
AGTGCCGGGCCCCTCATAGGAGAGGAAGCTCGGGAGGTGGCCAGGCGGCAGGAAGGCGCACCCCCCCAGCAATCCGCGC
GCCGGGACAGAATGCCCTGCAGGAACTTCTTCTGGAAGACCTTCTCCTCCTGCAAATAAAACCTCACCCATGAATGCTC
ACGCAAGTTTAATTACAGACCTGAA
```

##### Output

The output of the `gto_amino_acid_from_seq` program is an amino acid sequence.
Using the input above, an output example for this is the following:

```
TRRPPAAAALRGHGPGGSTAALLPLSPAPPKEKQPPDFPRLGRDSEHMQEAAGSGLSGPPGPS-ERKLGRWPGGRKQAS
AANPRAGTESPAKPCRNFFWKTFSTPPAKTSPMNAHASLITDLKQDAIVPRPPAAAALRGHGHRCPAPGGWPHRPRQRA
YAGSGRNKEKQPPDFPRLVV-VDLPGQCRAPHRRGSSGGGQAAGRRTPPAIRAPGQNALQELLLEDLLLLQIKPHP-ML
TQV-LQT-
```

### Chapter 6

### General purpose tools

The toolkit also has a set of tools with a more general-purpose, which were not designed to work with a specific data format. Instead, it was developed as an auxiliary component to help the construction of pipelines combining all the described subsets. This contains tools for char manipulations, such as reversing, segmentation and permutation, for manipulating numerical scores, such sum, filter, calculate the min and the max of a numeric matrix mainly originated from the tools' outputs. The current available tools for general purposes are:

1. `gto_char_to_line`: it splits a sequence into lines, creating an output sequence which has a char for each line.
2. `gto_new_line_on_new_x`: it splits different rows with a new empty row.
3. `gto_upper_bound`: it sets an upper bound in a file with a value per line.
4. `gto_lower_bound`: it sets an lower bound in a file with a value per line.
5. `gto_brute_force_string`: it generates all combinations, line by line, for an inputted alphabet and specific size.
6. `gto_real_to_binary_with_threshold`: it converts a sequence of real numbers into a binary sequence, given a threshold.
7. `gto_sum`: it adds decimal values in file, line by line, splitted by spaces or tabs.
8. `gto_filter`: it filters numerical sequences.
9. `gto_word_search`: it search for a word in a file.
10. `gto_permute_by_blocks`: it permutates by block sequence, FASTA and Multi-FASTA files.
11. `gto_info`: it gives the basic properties of the file, namely size, cardinality, distribution percentage of the symbols, among others.
12. `gto_segment`: it segments a filtered sequence.

13. `gto_comparative_map`: it creates a visualization for comparative maps.

14. `gto_max`: it computes the maximum value in each row between two files.

15. `gto_min`: it computes the minimum value in each row between two files.

#### 6.1 Program gto_char_to_line

The `gto_char_to_line` splits a sequence into lines, creating an output sequence which has a char for each line.
For help type:

```
./gto_char_to_line -h
```

In the following subsections, we explain the input and output paramters.

##### Input parameters

The `gto_char_to_line` program needs two streams for the computation, namely the input and output standard. The input stream is a sequence file.
The attribution is given according to:

```
Usage: ./gto_char_to_line [options] [[--] args]
   or: ./gto_char_to_line [options]

It splits a sequence into lines, creating an output sequence which has a char for each line.

    -h, --help         show this help message and exit

Basic options
    < input.seq        Input sequence file (stdin)
    > output.seq       Output sequence file (stdout)

Example: ./gto_char_to_line < input.seq > output.seq
```

An example of such an input file is:

```
ACAAGACGGCCTCCTGCTGCTGCTGCTCTCCGGGGCCACGGCCCTGGAGGGTCCACCGCTGCCCTGCTGCCATTGTCCCC
GGCCCCACCTAAGGAAAAGCAGCCTCCTGACTTTCCTCGCTTGGGCCGAGACAGCGAGCATATGCAGGAAGCGGCAGGAA
GTGGTTTGAGTGGACCTCCGGGCCCCTCATAGGAGAGGAAGCTCGGGAGGTGGCCAGGCGGCAGGAAGCAGGCCAGTGCC
GCGAATCCGCGCGCCGGGACAGAATCTCCTGCAAAGCCCTGCAGGAACTTCTTCTGGAAGACCTTCTCCACCCCCCCAGC
TAAAACCTCACCCATGAATGCTCACGCAAGTTTAATTACAGACCTGAAACAAGATGCCATTGTCCCCCGGCCTCCTGCTG
CTGCTGCTCTCCGGGGCCACGGCCACCGCTGCCCTGCCCCTGGAGGGTGGCCCCACCGGCCGAGACAGCGAGCATATGCA
GGAAGCGGCAGGAATAAGGAAAAGCAGCCTCCTGACTTTCCTCGCTTGGTGGTTTGAGTGGACCTCCCAGGCCAGTGCCG
GGCCCCTCATAGGAGAGGAAGCTCGGGAGGTGGCCAGGCGGCAGGAAGGCGCACCCCCCCAGCAATCCGCGCGCCGGGAC
AGAATGCCCTGCAGGAACTTCTTCTGGAAGACCTTCTCCTCCTGCAAATAAAACCTCACCCATGAATGCTCACGCAAGTT
TAATTACAGACCTGAA
```

##### Output

The output of the `gto_char_to_line` program is a group sequence splited by \n foreach character.
Using the input above, an output example for this is the following:

```
A
C
A
A
G
A
C
G
G
C
C
T
C
C
T
G
C
T
G
C
T
...
```

#### 6.2 Program gto_new_line_on_new_x

The `gto_new_line_on_new_x` splits different rows with a new empty row.
For help type:

```
./gto_new_line_on_new_x -h
```

In the following subsections, we explain the input and output paramters.

##### Input parameters

The `gto_new_line_on_new_x` program needs two streams for the computation, namely the input and output standard. The input stream is a matrix file format with 3 columns.
The attribution is given according to:

```
Usage: ./gto_new_line_on_new_x [options] [[--] args]
   or: ./gto_new_line_on_new_x [options]

It splits different rows with a new empty row.

    -h, --help    show this help message and exit

```

```
Basic options
    < input        Input file with 3 column matrix format (stdin)
    > output       Output file with 3 column matrix format (stdout)

Example: ./gto_new_line_on_new_x < input > output
```

An example of such an input file is:

```
1   2   2
1   2   2
4   4   1
10  12  2
15  15  1
45  47  3
45  47  3
45  47  3
45  47  3
55  55  1
```

##### Output

The output of the `gto_new_line_on_new_x` program is a 3 column matrix, with an empty line between different rows.

Using the input above, an output example for this is the following:

```
1.000000    2.000000    2.000000
1.000000    2.000000    2.000000

4.000000    4.000000    1.000000

10.000000   12.000000   2.000000

15.000000   15.000000   1.000000

45.000000   47.000000   3.000000
45.000000   47.000000   3.000000
45.000000   47.000000   3.000000
45.000000   47.000000   3.000000

55.000000   55.000000   1.000000
```

#### 6.3 Program gto_upper_bound

The `gto_upper_bound` sets an upper bound in a file with a value per line.

For help type:

```
./gto_upper_bound -h
```

In the following subsections, we explain the input and output paramters.

##### Input parameters

The `gto_upper_bound` program needs two streams for the computation, namely the input and output standard. The input stream is a numeric file.
The attribution is given according to:

```
Usage: ./gto_upper_bound [options] [[--] args]
   or: ./gto_upper_bound [options]

It sets an upper bound in a file with a value per line.

    -h, --help                show this help message and exit

Basic options
    -u, --upperbound=<int>    The upper bound value
    < input.num               Input numeric file (stdin)
    > output.num              Output numeric file (stdout)

Example: ./gto_upper_bound -u <upperbound> < input.num > output.num
```

An example of such an input file is:

```
0.123
3.432
2.341
1.323
7.538
4.122
0.242
0.654
5.633
```

##### Output

The output of the `gto_upper_bound` program is a set of numbers truncated at the a defined upper bound. Using the input above, an output example for this is the following:

```
Using upper bound: 4
0.123000
3.432000
2.341000
1.323000
4.000000
4.000000
0.242000
0.654000
4.000000
```

#### 6.4 Program gto_lower_bound

The `gto_lower_bound` sets an lower bound in a file with a value per line.
For help type:

```
./gto_lower_bound -h
```

In the following subsections, we explain the input and output paramters.

##### Input parameters

The `gto_lower_bound` program needs two streams for the computation, namely the input and output standard. The input stream is a numeric file.
The attribution is given according to:

```
Usage: ./gto_lower_bound [options] [[--] args]
   or: ./gto_lower_bound [options]

It sets an lower bound in a file with a value per line.

    -h, --help                  show this help message and exit

Basic options
    -l, --lowerbound=<int>      The lower bound value
    < input.num                 Input numeric file (stdin)
    > output.num                Output numeric file (stdout)

Example: ./gto_lower_bound -l <lowerbound> < input.num > output.num
```

An example of such an input file is:

```
0.123
3.432
2.341
1.323
7.538
4.122
0.242
0.654
5.633
```

##### Output

The output of the `gto_lower_bound` program is a set of numbers truncated at the a defined lower bound.
Using the input above, an output example for this is the following:

```
Using lower bound: 2
2.000000
3.432000
2.341000
```

```
2.000000
7.538000
4.122000
2.000000
2.000000
5.633000
```

#### 6.5 Program gto_brute_force_string

The `gto_brute_force_string` generates all combinations, line by line, for an inputted alphabet and specific size.
For help type:

```
./gto_brute_force_string -h
```

In the following subsections, we explain the input and output paramters.

##### Input parameters

The `gto_brute_force_string` program needs some paramenters for the computation, namely the alphabet and the key size.
The attribution is given according to:

```
Usage: ./gto_brute_force_string [options] [[--] args]
   or: ./gto_brute_force_string [options]

It generates all combinations, line by line, for an inputted alphabet and specific size.

    -h, --help             show this help message and exit

Basic options
    -a, --alphabet=<str>  The input alphabet
    -s, --size=<int>      The combinations size
    > output              Output all the combinations (stdout)

Example: ./gto_brute_force_string -a <alphabet> -s <size> > output
```

##### Output

The output of the `gto_brute_force_string` program is a set of all possible word combinations with a defined size, using the input alphabet.
Using the input above with the alphabet ”abAB” with the word size of 3, an output example for this is the following:

```
aaa
aab
aaA
aaB
```

```
aba
...
BBb
BBA
BBB
```

#### 6.6 Program gto_real_to_binary_with_threshold

The **gto_real_to_binary_with_threshold** converts a sequence of real numbers into a binary sequence, given a threshold. The numbers below to the threshold will be 0.
For help type:

```
./gto_real_to_binary_with_threshold -h
```

In the following subsections, we explain the input and output paramters.

##### Input parameters

The **gto_real_to_binary_with_threshold** program needs two streams for the computation, namely the real sequence as input. These numbers should be splitted by lines.
The attribution is given according to:

```
Usage: ./gto_real_to_binary_with_threshold [options] [[--] args]
   or: ./gto_real_to_binary_with_threshold [options]

It converts a sequence of real numbers into a binary sequence given a threshold.

    -h, --help                show this help message and exit

Basic options
    -t, --threshold=<dbl>     The threshold in real format
    < input.num               Input numeric file (stdin)
    > output.bin              Output binary file (stdout)

Example: ./gto_real_to_binary_with_threshold -t <threshold> < input.num > output.bin
```

An example of such an input file is:

```
12.25
1.2
5.44
5.51
7.97
2.34
8.123
```

##### Output

The output of the **gto_real_to_binary_with_threshold** program is a binary sequence.
Using the input above with the threshold of 5.5, an output example for this is the following:

```
1
0
0
1
1
0
1
```

#### 6.7 Program gto_sum

The `gto_sum` adds decimal values in file, line by line, splitted by spaces or tabs.
For help type:

```
./gto_sum -h
```

In the following subsections, we explain the input and output paramters.

##### Input parameters

The `gto_sum` program needs two streams for the computation, namely the input, which is a decimal file. The attribution is given according to:

```
Usage: ./gto_sum [options] [[--] args]
   or: ./gto_sum [options]

It adds decimal values in file, line by line, splitted by spaces or tabs.

    -h, --help         show this help message and exit

Basic options
    < input.num        Input numeric file (stdin)
    > output.num       Output numeric file (stdout)

Optional
    -r, --sumrows      When active, the application adds all the values line by line
    -a, --sumall       When active, the application adds all values

Example: ./gto_sum -a < input.num > output.num
```

An example of such an input file is:

```
0.123   5   5
3.432
2.341   3   2
1.323
7.538   5
4.122
0.242
0.654
```

```
5.633   10
```

##### Output

The output of the gto_sum program is a sum of the elements in the input file.
Executing the application with the provided input and with the flag to add only the elements in each row, the output of this execution is:

```
10.123000
3.432000
7.341000
1.323000
12.538000
4.122000
0.242000
0.654000
15.633000
```

#### 6.8 Program gto_filter

The gto_filter filters numerical sequences using a low-pass filter.
For help type:

```
./gto_filter -h
```

In the following subsections, we explain the input and output paramters.

##### Input parameters

The gto_filter program needs two streams for the computation, namely the input and output standard. The input stream is a numeric file.
The attribution is given according to:

```
Usage: ./gto_filter [options] [[--] args]
   or: ./gto_filter [options]

It filters numerical sequences using a low-pass filter.

    -h, --help                show this help message and exit

Basic options
    < input.num               Input numeric file (stdin)
    > output.num              Output numeric file (stdout)

Optional
    -w, --windowsize=<int>    Window size (defaut 0)
    -d, --drop=<int>          Discard elements (default 0.0)
    -t, --windowtype=<int>    Window type (0=Hamm, 1=Hann, 2=Black, 3=rec) (default 0 (Hamm))
    -c, --onecolumn           Read from one column
```

```
    -p, --printone            Print one column
    -r, --reverse             Reverse mode

Example: ./gto_filter -w <windowsize> -d <drop> -t <windowtype> -c -p -r < input.num > output.num
```

An example of such an input file is:

```
1   1.77
5   2.18
10  2.32
15  3.15
20  2.52
25  4.43
30  1.23
```

##### Output

The output of the `gto_filter` program is a numeric file, identical of the input.
Using the input above with the window size of 3, an output example for this is the following:

```
Got 7 entries from file
1   2.085
5   2.256
10  2.507
15  2.757
20  2.905
25  2.860
30  2.674
```

#### 6.9 Program gto_word_search

The `gto_word_search` search for a word in a file. It is case sensitive.
For help type:

```
./gto_word_search -h
```

In the following subsections, we explain the input and output paramters.

##### Input parameters

The `gto_word_search` program needs two streams for the computation, namely the input and output standard. The input stream is a text file.
The attribution is given according to:

```
Usage: ./gto_word_search [options] [[--] args]
   or: ./gto_word_search [options]

Searching for a word in a text file. It is case sensitive.
```

```
    -h, --help          show this help message and exit

Basic options
    -w, --word=<str>  Word to search in the file
    < input.txt       Input text file (stdin)
    > output.txt      Output text file (stdout)

Example: ./gto_word_search -w <word> < input.txt > output.txt
```

An example of such an input file is:

```
No guts, no story. Chris Brady
My life is my message. Mahatma Gandhi
Screw it, letâĂŹs do it. Richard Branson
Boldness be my friend. William Shakespeare
Keep going. Be all in. Bryan Hutchinson
My life is my argument. Albert Schweitzer
Fight till the last gasp. William Shakespeare
Leave no stone unturned. Euripides
```

##### Output

The output of the `gto_word_search` program is a text file with the matching paragraphs and the location of the word found.
Using the input above with the word "Shakespeare", an output example for this is the following:

```
Found match in range [ 1536 : 2048 ]
Boldness be my friend. William Shakespeare

Found match in range [ 3072 : 3584 ]
Fight till the last gasp. William Shakespeare
```

#### 6.10 Program gto_permute_by_blocks

The `gto_permute_by_blocks` permutates by block sequence, FASTA and Multi-FASTA files.
For help type:

```
./gto_ -h
```

In the following subsections, we explain the input and output paramters.

##### Input parameters

The `gto_permute_by_blocks` program needs two streams for the computation, namely the input and output standard. The input stream is a sequence, FASTA or Multi-FASTA file.
The attribution is given according to:

```
Usage: ./gto_permute_by_blocks [options] [[--] args]
   or: ./gto_permute_by_blocks [options]

It permutates by block sequence, FASTA and Multi-FASTA files.

    -h, --help             show this help message and exit

Basic options
    -b, --numbases=<int>  The number of bases in each block
    -s, --seed=<int>      Starting point to the random generator
    < input               Input sequence, FASTA or Multi-FASTA file format (stdin)
    > output              Output sequence, FASTA or Multi-FASTA file format (stdout)

Example: ./gto_permute_by_blocks -b <numbases> -s <seed> < input.fasta > output.fasta
```

An example of such an input file is:

```
>AB000264 |acc=AB000264|descr=Homo sapiens mRNA
ACAAGACGGCCTCCTGCTGCTGCTGCTCTCCGGGGCCACGGCCCTGGAGGGTCCACCGCTGCCCTGCTGCCATTGTCCCC
GGCCCCACCTAAGGAAAAGCAGCCTCCTGACTTTCCTCGCTTGGGCCGAGACAGCGAGCATATGCAGGAAGCGGCAGGAA
GTGGTTTGAGTGGACCTCCGGGCCCCTCATAGGAGAGGAAGCTCGGGAGGTGGCCAGGCGGCAGGAAGCAGGCCAGTGCC
GCGAATCCGCGCGCCGGGACAGAATCTCCTGCAAAGCCCTGCAGGAACTTCTTCTGGAAGACCTTCTCCACCCCCCCAGC
TAAAACCTCACCCATGAATGCTCGCAACACGCAAGTTTAATTCGCAAGTTAGACCTGAACGGGAGGTGGCCACGCAAGTT
```

##### Output

The output of the `gto_permute_by_blocks` program is a sequence, FASTA or Multi-FASTA file permuted following some parameters.
Using the input above with the base number as 80, an output example for this is the following:

```
GGCCCCACCTAAGGAAAAGCAGCCTCCTGACTTTCCTCGCTTGGGCCGAGACAGCGAGCATATGCAGGAAGCGGCAGGAA
GTGGTTTGAGTGGACCTCCGGGCCCCTCATAGGAGAGGAAGCTCGGGAGGTGGCCAGGCGGCAGGAAGCAGGCCAGTGCC
GCGAATCCGCGCGCCGGGACAGAATCTCCTGCAAAGCCCTGCAGGAACTTCTTCTGGAAGACCTTCTCCACCCCCCCAGC
ACAAGACGGCCTCCTGCTGCTGCTGCTCTCCGGGGCCACGGCCCTGGAGGGTCCACCGCTGCCCTGCTGCCATTGTCCCC
TAAAACCTCACCCATGAATGCTCGCAACACGCAAGTTTAATTCGCAAGTTAGACCTGAACGGGAGGTGGCCACGCAAGTT
```

#### 6.11 Program gto_info

The `gto_info` gives the basic properties of the file, namely size, cardinality, distribution percentage of the symbols, among others.
For help type:

```
./gto_info -h
```

In the following subsections, we explain the input and output paramters.

#### Input parameters

The `gto_info` program needs two streams for the computation, namely the input and output standard. The input stream is a file withou any specific format.

The attribution is given according to:

```
Usage: ./gto_info [options] [[--] args]
   or: ./gto_info [options]

It gives the basic properties of the file, namely size, cardinality, distribution
percentage of the symbols, among others.

    -h, --help    show this help message and exit

Basic options
    < input       Input file (stdin)
    > output      Output read information (stdout)

Optional
    -a, --ascii   When active, the application shows the ASCII codes

Example: ./gto_info < input > output

Output example :
Number of symbols  : value
Alphabet size      : value
Alphabet           : value
Symbol distribution:
<Symbol/Code ASCII>  <Symbol count>  <Distribution percentage>
```

An example of such an input file is:

```
>AB000264 |acc=AB000264|descr=Homo sapiens mRNA
ACAAGACGGCCTCCTGCTGCTGCTGCTCTCCGGGGCCACGGCCCTGGAGGGTCCACCGCTGCCCTGCTGCCATTGTCCCC
GGCCCCACCTAAGGAAAAGCAGCCTCCTGACTTTCCTCGCTTGGGCCGAGACAGCGAGCATATGCAGGAAGCGGCAGGAA
GTGGTTTGAGTGGACCTCCGGGCCCCTCATAGGAGAGGAAGCTCGGGAGGTGGCCAGGCGGCAGGAAGCAGGCCAGTGCC
GCGAATCCGCGCGCCGGGACAGAATCTCCTGCAAAGCCCTGCAGGAACTTCTTCTGGAAGACCTTCTCCACCCCCCCAGC
TAAAACCTCACCCATGAATGCTCGCAACACGCAAGTTTAATTCGCAAGTTAGACCTGAACGGGAGGTGGCCACGCAAGTT
```

#### Output

The output of the `gto_info` program is a set of information related to the file read.

Using the input above, an output example for this is the following:

```
Number of symbols  : 453
Alphabet size      : 28
Alphabet           :|srponmiedcaTRNHGCBA>=6420 \n
Symbol distribution:
|  :    2      0.4415011
s  :    3      0.66225166
r  :    1      0.22075055
```

```
p  :    1       0.22075055
o  :    2       0.4415011
n  :    1       0.22075055
m  :    2       0.4415011
i  :    1       0.22075055
e  :    2       0.4415011
d  :    1       0.22075055
c  :    3       0.66225166
a  :    2       0.4415011
T  :    66      14.569536
R  :    1       0.22075055
N  :    1       0.22075055
H  :    1       0.22075055
G  :    117     25.827815
C  :    131     28.918322
B  :    2       0.4415011
A  :    89      19.646799
>  :    1       0.22075055
=  :    2       0.4415011
6  :    2       0.4415011
4  :    2       0.4415011
2  :    2       0.4415011
0  :    6       1.3245033
   :    4       0.88300221
\n :    5       1.1037528
```

#### 6.12 Program gto_segment

The `gto_segment` segments a filtered sequence.

For help type:

```
./gto_segment -h
```

In the following subsections, we explain the input and output paramters.

##### Input parameters

The `gto_segment` program needs two streams for the computation, namely the input and output standard. The input stream is a numeric file.

The attribution is given according to:

```
Usage: ./gto_segment [options] [[--] args]
   or: ./gto_segment [options]

It segments a filtered sequence.

    -h, --help                show this help message and exit

Basic options
    -t, --threshold=<dbl>     The segment threshold
    < input.num               Input numeric file (stdin)
```

```
    > output                    Output the segment file (stdout)

Example: ./gto_segment -t <threshold> < input.num > output
```

An example of such an input file is:

```
1   1.77
5   2.18
10  2.32
15  3.15
20  2.52
25  4.43
30  1.23
```

##### Output

The output of the gto_segment program is the interval of values âĂŃâĂŃbelow the threshold.
Using the input above with a threshold of 3, an output example for this is the following:

```
0:10
```

#### 6.13 Program gto_comparative_map

The gto_comparative_map creates a visualization for comparative maps.
For help type:

```
./gto_comparative_map -h
```

In the following subsections, we explain the input and output paramters.

##### Input parameters

The gto_comparative_map program needs an input file with the plot positions, respecting a defined structure.
The attribution is given according to:

```
Usage: ./gto_comparative_map [options] [[--] args]
   or: ./gto_comparative_map [options]

It creates a visualization for comparative maps.

    -h, --help             Show this help message and exit

Basic options
    <FILE>                 Contigs filename with positions (.pos),

Optional

    -h                     Give this help,
```

```
    -V                      Display version number,
    -v                      Verbose mode (more information),
    -l <link>               Link type between maps [0;4],
    -w <width>              Chromosome width,
    -s <space>              Space between chromosomes,
    -m <mult>               Color id multiplication factor,
    -b <begin>              Color id beggining,
    -c <minimum>            Minimum block size to consider,
    -i                      Do NOT show inversion maps,
    -r                      Do NOT show regular maps,
    -o <FILE>               Output image filename with map,

Example: ./gto_comparative_map -o map.svg map.config
```

An example of such an input file is:

```
#SCF    5000000 5000000
aaa     1       1000000 1       1000000 bbbb    3000000 4000000 3000000 4000000
bbb     1500000 2000000 1500000 2000000 cccc    1500000 2000000 1500000 2000000
aaa     2000000 3000000 2000000 3000000 bbbb    3000000 2000000 3000000 2000000
```

##### Output

The output of the `gto_comparative_map` program is a executing report, and a svg plot with the maps. Using the input above, an output example for this is the following:

```
==[ PROCESSING ]====================
Printing plot ...
Found 2 regular regions.
Found 1 inverted regions.
Done!

==[ STATISTICS ]====================
Total cpu time: 0 second(s).
```

In the Figure 6.1 is represented the plot for the execution above.

#### 6.14 Program gto_max

The `gto_max` computes the maximum value in each row between two files.
For help type:

```
./gto_max -h
```

In the following subsections, we explain the input and output paramters.

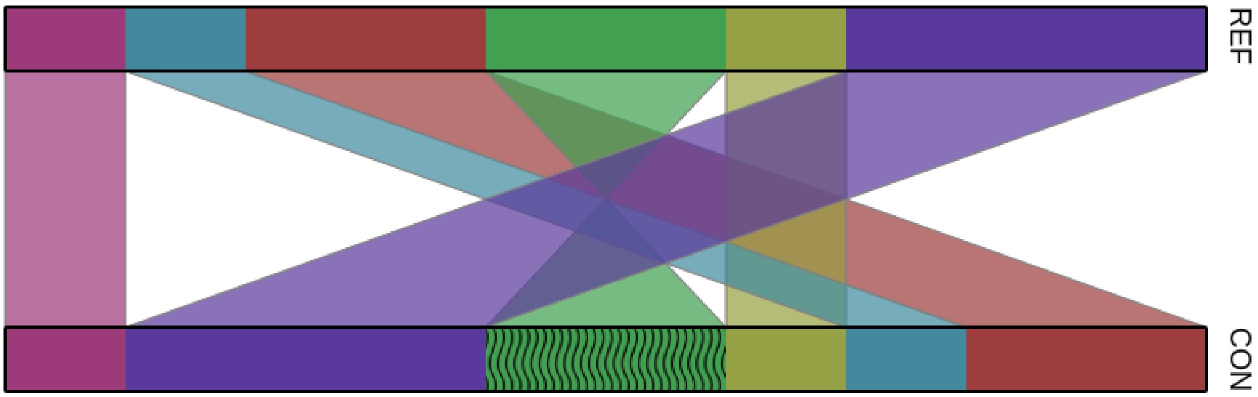

Figure 6.1: `gto_comparative_map` execution plot.

##### Input parameters

The `gto_max` program needs two streams for the computation, namely the input, which are two decimal files.

The attribution is given according to:

```
Usage: ./gto_max [options] [[--] args]
   or: ./gto_max [options]

It computes the maximum value in each row between two files.

    -h, --help                Show this help message and exit

Basic options
    -f, --first_file=<str>    File to compute the max
    -s, --second_file=<str>   The second file to do the max computation
    > output.num              Output numeric file (stdout)

Example: ./gto_max -f input1.num -s input2.num > output.num
```

An example of such an input files are:

File 1:

```
0.123
3.432
2.341
1.323
7.538
4.122
0.242
0.654
5.633
```

File 2:

```
2.123
5.312
2.355
0.124
1.785
3.521
0.532
7.324
2.312
```

##### Output

The output of the gto_max program is the numeric file with the maximum value for each row between both input files.
Executing the application with the provided input, the output of this execution is:

```
2.123000
5.312000
2.355000
1.323000
7.538000
4.122000
0.532000
7.324000
5.633000
```

#### 6.15 Program gto_min

The gto_min computes the minium value in each row between two files.
For help type:

```
./gto_min -h
```

In the following subsections, we explain the input and output paramters.

##### Input parameters

The gto_min program needs two streams for the computation, namely the input, which are two decimal files.
The attribution is given according to:

```
Usage: ./gto_min [options] [[--] args]
   or: ./gto_min [options]

It computes the minimum value in each row between two files.

    -h, --help                Show this help message and exit
```

```
Basic options
    -f, --first_file=<str>    File to compute the max
    -s, --second_file=<str>   The second file to do the max computation
    > output.num              Output numeric file (stdout)

Example: ./gto_min -f input1.num -s input2.num > output.num
```

An example of such an input files are:

File 1:

```
0.123
3.432
2.341
1.323
7.538
4.122
0.242
0.654
5.633
```

File 2:

```
2.123
5.312
2.355
0.124
1.785
3.521
0.532
7.324
2.312
```

##### Output

The output of the `gto_min` program is the numeric file with the minimum value for each row between both input files.

Executing the application with the provided input, the output of this execution is:

```
0.123000
3.432000
2.341000
0.124000
1.785000
3.521000
0.242000
0.654000
2.312000
```

### Bibliography

[1] E. R. Mardis, "Dna sequencing technologies: 2006–2016," *Nature protocols*, vol. 12, no. 2, p. 213, 2017.

[2] C. Brouwer, T. D. Vu, M. Zhou, G. Cardinali, M. M. Welling, N. van de Wiele, and V. Robert, "Current opportunities and challenges of next generation sequencing (ngs) of dna; determining health and disease," *British Biotechnology Journal*, vol. 13, no. 4, 2016.

[3] L. Liu, Y. Li, S. Li, N. Hu, Y. He, R. Pong, D. Lin, L. Lu, and M. Law, "Comparison of next-generation sequencing systems," *BioMed Research International*, vol. 2012, 2012.

[4] H. Zhang, "Overview of sequence data formats," in *Statistical Genomics.* Springer, 2016, pp. 3–17.

[5] P. J. Cock, C. J. Fields, N. Goto, M. L. Heuer, and P. M. Rice, "The sanger fastq file format for sequences with quality scores, and the solexa/illumina fastq variants," *Nucleic acids research*, vol. 38, no. 6, pp. 1767–1771, 2009.

[6] A. P. Droop, "fqtools: an efficient software suite for modern fastq file manipulation," *Bioinformatics*, vol. 32, no. 12, pp. 1883–1884, 2016.

[7] A. Gordon, G. Hannon *et al.*, "Fastx-toolkit," *FASTQ/A short-reads preprocessing tools (unpublished) http://hannonlab. cshl. edu/fastx_toolkit*, vol. 5, 2010.

[8] E. Afgan, D. Baker, B. Batut, M. Van Den Beek, D. Bouvier, M. Čech, J. Chilton, D. Clements, N. Coraor, B. A. Grüning *et al.*, "The galaxy platform for accessible, reproducible and collaborative biomedical analyses: 2018 update," *Nucleic acids research*, vol. 46, no. W1, pp. W537–W544, 2018.

[9] M. A. DePristo, E. Banks, R. Poplin, K. V. Garimella, J. R. Maguire, C. Hartl, A. A. Philippakis, G. Del Angel, M. A. Rivas, M. Hanna *et al.*, "A framework for variation discovery and genotyping using next-generation dna sequencing data," *Nature genetics*, vol. 43, no. 5, p. 491, 2011.

[10] S. Kumar, G. Stecher, and K. Tamura, "Mega7: molecular evolutionary genetics analysis version 7.0 for bigger datasets," *Molecular biology and evolution*, vol. 33, no. 7, pp. 1870–1874, 2016.

[11] W. Shen, S. Le, Y. Li, and F. Hu, "Seqkit: a cross-platform and ultrafast toolkit for fasta/q file manipulation," *PLoS One*, vol. 11, no. 10, p. e0163962, 2016.

[12] J. Goecks, A. Nekrutenko, and J. Taylor, "Galaxy: a comprehensive approach for supporting accessible, reproducible, and transparent computational research in the life sciences," *Genome biology*, vol. 11, no. 8, p. R86, 2010.

[13] D. Blankenberg, A. Gordon, G. Von Kuster, N. Coraor, J. Taylor, A. Nekrutenko, and G. Team, "Manipulation of fastq data with galaxy," *Bioinformatics*, vol. 26, no. 14, pp. 1783–1785, 2010.

[14] G. A. Van der Auwera, M. O. Carneiro, C. Hartl, R. Poplin, G. Del Angel, A. Levy-Moonshine, T. Jordan, K. Shakir, D. Roazen, J. Thibault *et al.*, "From fastq data to high-confidence variant calls: the genome analysis toolkit best practices pipeline," *Current protocols in bioinformatics*, vol. 43, no. 1, pp. 11–10, 2013.

[15] K. Tamura, D. Peterson, N. Peterson, G. Stecher, M. Nei, and S. Kumar, "Mega5: molecular evolutionary genetics analysis using maximum likelihood, evolutionary distance, and maximum parsimony methods," *Molecular biology and evolution*, vol. 28, no. 10, pp. 2731–2739, 2011.

[16] M. Hosseini, D. Pratas, and A. Pinho, "A survey on data compression methods for biological sequences," *Information*, vol. 7, no. 4, p. 56, 2016.

[17] Y. Liu, H. Peng, L. Wong, and J. Li, "High-speed and high-ratio referential genome compression," *Bioinformatics*, vol. 33, no. 21, pp. 3364–3372, 2017.

[18] I. Ochoa, M. Hernaez, and T. Weissman, "idocomp: a compression scheme for assembled genomes," *Bioinformatics*, vol. 31, no. 5, pp. 626–633, 2014.

[19] D. Pratas, A. J. Pinho, and P. J. Ferreira, "Efficient compression of genomic sequences," in *2016 Data Compression Conference (DCC)*. IEEE, 2016, pp. 231–240.

[20] S. Deorowicz, A. Danek, and M. Niemiec, "Gdc 2: Compression of large collections of genomes," *Scientific reports*, vol. 5, p. 11565, 2015.

[21] M. Hernaez, D. Pavlichin, T. Weissman, and I. Ochoa, "Genomic data compression," *Annual Review of Biomedical Data Science*, vol. 2, 2019.

[22] Ö. Nalbantoglu, D. Russell, and K. Sayood, "Data compression concepts and algorithms and their applications to bioinformatics," *Entropy*, vol. 12, no. 1, pp. 34–52, 2010.

[23] M. Hosseini, D. Pratas, and A. J. Pinho, "Ac: A compression tool for amino acid sequences," *Interdisciplinary Sciences: Computational Life Sciences*, pp. 1–9, 2019.

[24] D. Pratas, M. Hosseini, and A. J. Pinho, "Compression of amino acid sequences," in *International Conference on Practical Applications of Computational Biology & Bioinformatics*. Springer, 2018, pp. 105–113.

[25] W. Huang, L. Li, J. R. Myers, and G. T. Marth, "Art: a next-generation sequencing read simulator," *Bioinformatics*, vol. 28, no. 4, pp. 593–594, 2011.

[26] A. Price and C. Gibas, "Simulome: a genome sequence and variant simulator," *Bioinformatics*, vol. 33, no. 12, pp. 1876–1878, 2017.

[27] G. Baruzzo, K. E. Hayer, E. J. Kim, B. Di Camillo, G. A. FitzGerald, and G. R. Grant, "Simulation-based comprehensive benchmarking of rna-seq aligners," *Nature methods*, vol. 14, no. 2, p. 135, 2017.

[28] M. Escalona, S. Rocha, and D. Posada, "A comparison of tools for the simulation of genomic next-generation sequencing data," *Nature Reviews Genetics*, vol. 17, no. 8, p. 459, 2016.

[29] D. Pratas, A. J. Pinho, and J. M. Rodrigues, "Xs: a fastq read simulator," *BMC research notes*, vol. 7, no. 1, p. 40, 2014.
